# Neurotropic and non-neurotropic equid alphaherpesvirus 1 (EHV1) mobilize most histones within viral replication compartments

**DOI:** 10.1101/2025.08.01.668067

**Authors:** Kristen L Conn

**Affiliations:** Department of Veterinary Microbiology, University of Saskatchewan, Saskatoon, Saskatchewan, Canada

**Keywords:** Alphaherpesvirus, EHV1, linker histone, core histone, histone variants, FRAP, histone dynamics, histone exchange, viral chromatin

## Abstract

Equid alphaherpesvirus 1 (EHV1) is a DNA virus that causes severe disease outcomes in equids. Some EHV1 strains are neurotropic and cause disease in the central nervous system, whereas others are non-neurotropic and can cause negative reproductive outcomes. The molecular mechanisms that govern pathotype of individual EHV1 strains are not understood. However, EHV1 replication in the presence of epigenetic inhibitors suggests that neurotropic and non-neurotropic EHV1 are differentially susceptible to epigenetic silencing. Aside from this evidence, little is known about EHV1 chromatin or its regulation. Here, we used fluorescence recovery after photobleaching to characterize EHV1 lytic chromatin dynamics. Infection with neurotropic or non-neurotropic EHV1 mobilized all histones. Canonical (H2A, H2B, H3.1, H4) or variant (H2A.B, H2A.Z, H2A.X, macroH2A, H3.3) core and linker H1.2 histones were equally mobilized by either strain. Thus, there were no vast differences in histone mobility during neurotropic or non-neurotropic EHV1 infection. All histones except for H2A.B were more mobile within EHV1 replication compartments (RCs) than the surrounding infected-cell chromatin. The differential mobility of histones within domains enriched for viral or cellular chromatin is consistent with distinct mechanisms to assemble and regulate the chromatin associated with viral or host DNA. Histones were further mobilized within RCs in cells in which infection had further progressed. Such mobilization indicates that increased levels of EHV1 transcription, DNA replication, or protein expression directly or indirectly mobilize histones. The high histone mobility within EHV1 RCs is consistent with assembly of EHV1 genomes in very dynamic and unstable nucleosomes. These data support a model in which EHV1 limits genome silencing by preventing stable chromatin assembly, or destabilizing the chromatin assembled, with viral genomes during lytic infection. We propose that manipulation of histone dynamics represents a novel mechanism of epigenetic regulation adopted by alphaherpesviruses to maintain genome accessibility and prevent gene silencing.

**Author summary:** DNA viruses are subjected to epigenetic regulation that silences or promotes gene expression. Multiple epigenetic mechanisms contribute to stabilize chromatin to silence gene expression or destabilize it to promote gene expression. Knowledge of the mechanisms whereby viruses prevent or overcome genome silencing and promote expression of their genes is important to understand how viruses, including alphaherpesviruses, take over the host cell to establish productive infection. Here we show that EHV1 broadly mobilizes histones within nuclear domains enriched in viral chromatin. Histone mobilization destabilizes chromatin and is consistent with the assembly of EHV1 genomes in dynamic, unstable nucleosomes. The manipulation of histone mobility is a phenomenon first described for the alphaherpesvirus herpes simplex virus 1 (HSV1). The conserved approach to dysregulate chromatin dynamics and mobilize histones represents a unique means whereby herpesviruses destabilize chromatin. Understanding the mechanisms that mobilize histones during infection will increase our general understanding of epigenetic regulation, which is important in the pathogenesis of infectious diseases and also of developmental or genetic ones. Moreover, knowledge of the processes whereby herpesviruses destabilize chromatin will support the development of novel therapeutics to maintain viral genomes in stable, silenced chromatin to prevent productive infection and development of associated diseases.

## Introduction

EHV1 is prevalent and highly infectious (1, 2). It initially infects and establishes primary lytic replication within epithelial cells lining the upper respiratory tract. Infection of monocytes, including CD172a^+^, recruited to EHV1-infected epithelial cells enables EHV1 to disseminate throughout the body in a cell-associated viremia (3, 4). Subsequent cell-to-cell contacts between infected monocytes and endothelial cells lining blood vessels transfers EHV1 to the endothelial cells where it establishes secondary lytic replication (5). This replication causes severe disease outcomes reflective of the affected organ(s).

Infection of cells lining vasculature within the central nervous system (CNS), for example, causes neurological symptoms including equine herpes myeloencephalopathy (EHM) (6). Whereas infection of cells lining vasculature within the uterus is associated with negative reproductive outcomes, including abortion and neonatal foal death (6). The molecular mechanisms underlying EHV1 pathotype are not yet well understood. One correlate is a single nucleotide polymorphism in the gene encoding the DNA polymerase (ORF 30) (7). Neurotropic strains typically encode a D at amino acid (a.a.) residue 752, whereas non-neurotropic strains typically encode an N (8, 9). However, this polymorphism is not strictly associated with, nor is it predictive, of neurotropism (10–12). Neurotropic EHV1 infection is characterized by more pronounced and extended cell-associated viremia subsequent to increased replication within epithelial cells, greater infection of, and faster replication kinetics within, immune cells, and increased transfer to endothelial cells (3, 13–15). The differences between neurotropic and non-neurotropic EHV1 replication may, at least in part, relate to chromatin regulation of viral gene expression. Within CD172a^+^ cells, neurotropic EHV1 strains are less sensitive to chromatin-mediated gene silencing than are non-neurotropic ones (13, 14). However neurotropic strains are sensitive to chromatin-mediated gene silencing within epithelial cells (16). To understand how chromatin regulation of EHV1 gene expression contributes to replication kinetics, and potentially strain pathotype, more knowledge of EHV1 chromatin is required.

Chromatin physically and functionally regulates DNA access for gene expression. The basic unit of chromatin is the nucleosome, a core histone octamer composed of two molecules each of histones H2A, H2B, H3, and H4 wrapped in approximately 147bp of DNA (17). Linker H1 histones bind to the DNA at nucleosome entry-exit sites to stabilize and compact chromatin. Histone-histone and histone-DNA interactions within and between nucleosomes regulate chromatin stability and compaction to control DNA access. Stabilization of such interactions promotes chromatin compaction to decrease DNA accessibility, whereas their destabilization promotes decompaction to increase DNA accessibility. Multiple factors contribute to regulate nucleosome stability, including the association of chromatin-binding or –regulatory proteins, posttranslational modifications (PTMs) of the histones assembled in nucleosomes, and the assembly of histone variants within nucleosomes. Histones are broadly categorized as canonical or variant based on their cell-cycle-related expression and primary mechanism of chromatin assembly (18–20). Variant histones have unique a.a. sequences that structurally impact nucleosome stability and functionally provide alternate residues for PTMs or interactions with other chromatin-regulatory or –binding proteins (21, 22). Histones H2A and H3 have distinct variants whereas H2B and H4 have no somatic variants in equids.

Histone H2A has the most numerous and diverse somatic variants. H2A.X most resembles canonical H2A, with 82% sequence identity. Even so, H2A.X-containing nucleosomes are less stable than H2A-containing ones due to increased DNA unwrapping at nucleosome entry-exit sites (23). H2A.X is well characterized for its role in the DNA damage response (DDR), where its phosphorylated form (γ-H2A.X) further destabilizes nucleosomes and decreases H1 association to facilitate DNA access for repair (23). H2A.Z shares only 60% sequence identity with canonical H2A and structurally differs in several key regions that decrease nucleosome stability and increase DNA unwrapping at nucleosome entry-exit sites (24). Consequently, H2A.Z-containing nucleosomes are also more unstable and accessible than H2A-containing ones (24, 25). H2A.Z regulates chromatin for multiple cellular processes, including transcription, heterochromatin formation and maintenance, DNA repair, and DNA replication(26). MacroH2A is a more distinct variant that is only 64% similar to H2A in its amino (N)-terminus and has an additional unique linker region connecting a macrodomain to its carboxyl (C)-terminus. MacroH2A mediates stronger intranucleosomal interactions to increases nucleosome stability, compaction, and inaccessibility (27–29). Consequently, macroH2A nucleosomes repress transcription and stabilize heterochromatin (30–32). H2A.B (H2A.Bbd) is the most divergent variant and shares only 48% sequence identity with canonical H2A. The H2A C-terminus and docking domain are truncated in H2A.B and it has a unique N-terminal arginine-rich region (33, 34). H2A.B nucleosomes wrap only 118bp of DNA, have weaker intranucleosomal interactions, and undergo transient DNA unwrapping at nucleosome entry-exit sites (34–36). H2A.B nucleosomes are therefore highly unstable and accessible, and are enriched in regions of high transcriptional activity and DNA synthesis (37–40).

H3 has two somatic variants, H3.3 and centromere-specific CENPA (centromeric protein A; CenH3). H3.3 is more than 96% identical to canonical H3.1 and H3.2, differing by only 5 or 4 a.a., respectively. H3.3 nucleosomes have similar stability as those assembled with canonical H3, however, H3.3 synergizes with co-assembled H2A variants, such as H2A.Z, to further regulate nucleosome stability and DNA association (25). H3.3 accumulates in transcriptionally active regions and telomeric heterochromatin (41–43). The more diverse CENPA is assembled in centromeric nucleosomes and shares only 45% sequence identity with canonical H3 (44). CENPA nucleosomes loosely wrap only 121bp of DNA, have transient DNA unwrapping at nucleosome entry-exit sites, and form an untwisted chromatin structure to increase CENPA-nucleosome accessibility and instability (45–47).

Histones continually exchange within chromatin to facilitate structural and functional regulation of chromatin at any given locus. Intrinsic histone exchange generally relates to their stability of association within chromatin. For example, H1 chromatin exchange is the fastest, while H2A-H2B dimers peripheral in the nucleosome undergo faster exchange than the H3-H4 dimers central in the nucleosome (48–58). For any given histone type, variants have inherently distinct exchange rates consistent with their effects on nucleosome stability such that destabilizing histones typically exchange faster than stabilizing ones (24). Nonetheless, even stabilizing histones assembled in condensed chromatin undergo exchange. Histone exchange is also regulated by external factors, including PTMs of the histones assembled in nucleosomes, the association of chromatin-interacting or –regulatory proteins, and processes that require DNA access (59).

Nucleosomes are partially or fully disassembled to enable DNA access and are then reassembled. Processes such as transcription that require access to the DNA therefore enhance histone exchange. Highly transcriptionally active chromatin regions have very fast rates of histone exchange to accommodate the DNA accessibility necessary for high levels of transcription. Accordingly, the most transcriptionally active chromatin, including nucleolar chromatin, is assembled in highly dynamic and unstable nucleosomes. Such dynamic chromatin is challenging to study due to inherent nucleosome instability and associated DNA hyperaccessibility. Consequently, the most dynamic and accessible chromatin appears as “nucleosome free” when evaluated by most common chromatin interrogation methods.

Histone exchange (chromatin dynamics) informs on viral chromatin. For example, the alphaherpesvirus HSV1 broadly dysregulates histone exchange. With exception of H2A.B, all evaluated linker and core histones are mobilized (have increased exchange) during lytic HSV1 infection (48–51, 60). Mobilization of histones from the cellular chromatin provides a source for the histones that assemble in HSV1 chromatin and also relates to composition of the viral chromatin (50, 51). Histones are most mobile within HSV1 replication compartments (RCs), which are nuclear domains enriched in viral chromatin (60). The high mobility of histones within HSV1 RCs is consistent with the assembly of viral genomes in highly dynamic, unstable, and accessible chromatin. Thus, measured histone mobility relates to the remarkable hyperaccessibility of the unique chromatin assembled with lytic HSV1 genomes (51, 61–64).

To initiate our investigation of EHV1 chromatin, we used fluorescence recovery after photobleaching (FRAP) as it is the only technique to directly measure histone mobility. Infection with neurotropic or non-neurotropic (abortogenic) EHV1 equally mobilized all evaluated canonical (H2A, H2B, H3.1, and H4) and variant (H2A.Z, H2A.X, H2A.B, macroH2A, and H3.3) core histones and linker histone H1.2. We show that all histones except for H2A.B were most mobile within nuclear domains enriched for EHV1 chromatin, the RCs. Mobilization of histones within RCs altered their chromatin residency to increase the net level of histones unbound from chromatin at any given time. Moreover, mobilization increased histone low-affinity chromatin exchange. The degree of histone mobilization within RCs apparently related to infection progression as histones were more mobile within medium-to large-sized ones than within small ones. Furthermore, within medium-to large-sized RCs histones were more mobile than within the nucleoli of mock-infected cells. These data indicate that lytic EHV1 chromatin is more dynamic or unstable than nucleolar chromatin, which is considered most unstable. Although histones were primarily mobilized within domains enriched for viral chromatin, H2A, H3.3, and H4 were also mobilized, albeit to a lesser degree, within the cellular chromatin surrounding EHV1 RCs. Conversely, linker histone H1.2 was less mobile within the infected-cell chromatin. Histone mobilization is a novel consequence of alphaherpesvirus infection that represents a unique and conserved chromatin regulatory mechanism to destabilize viral chromatin.

## Results

### EHV1 mobilizes core histones H2B and H4

We first evaluated mobility of core histones H2B and H4 as they have no somatic variants in equids that could be differentially mobilized by infection. Furthermore, H2B represents the more mobile H2A-H2B heterodimers flanking the nucleosome, while H4 represents the more stable H3-H4 heterodimers central to the nucleosome (17, 57). Transiently expressed GFP-H2B or –H4 had distinct granular localization with areas relatively enriched or depleted for GFP, consistent with GFP-histone assembly in chromatin (Fig 1, Mock). Most cells had nucleoli depleted for GFP-H2B or –H4, consistent with nucleolar chromatin instability (33, 65, 66). In cells infected with either abortogenic or neurotropic EHV1, viral RCs were evident as nuclear domains generally depleted for GFP-H2B or –H4, similar to the reported depletion of histones within HSV1 RCs (Fig 1, EHV1) (49, 60). As expected for field isolated strains, infection progression, as evaluated by RC number and size, was variable. Most infected cells had GFP-H2B or –H4 depleted regions clearly identifiable as RCs by size or shape (Fig 1, EHV1 “Large RC”; S1 Table). However, some infected cells had smaller GFP-H2B or –H4 depleted regions that more resembled the depleted regions within mock-infected cells (Fig 1, EHV1 “Small RC”). Regardless of the number or size of EHV1 RCs, nucleoli remained as discrete GFP-H2B or –H4 depleted domains.

**Fig 1.**
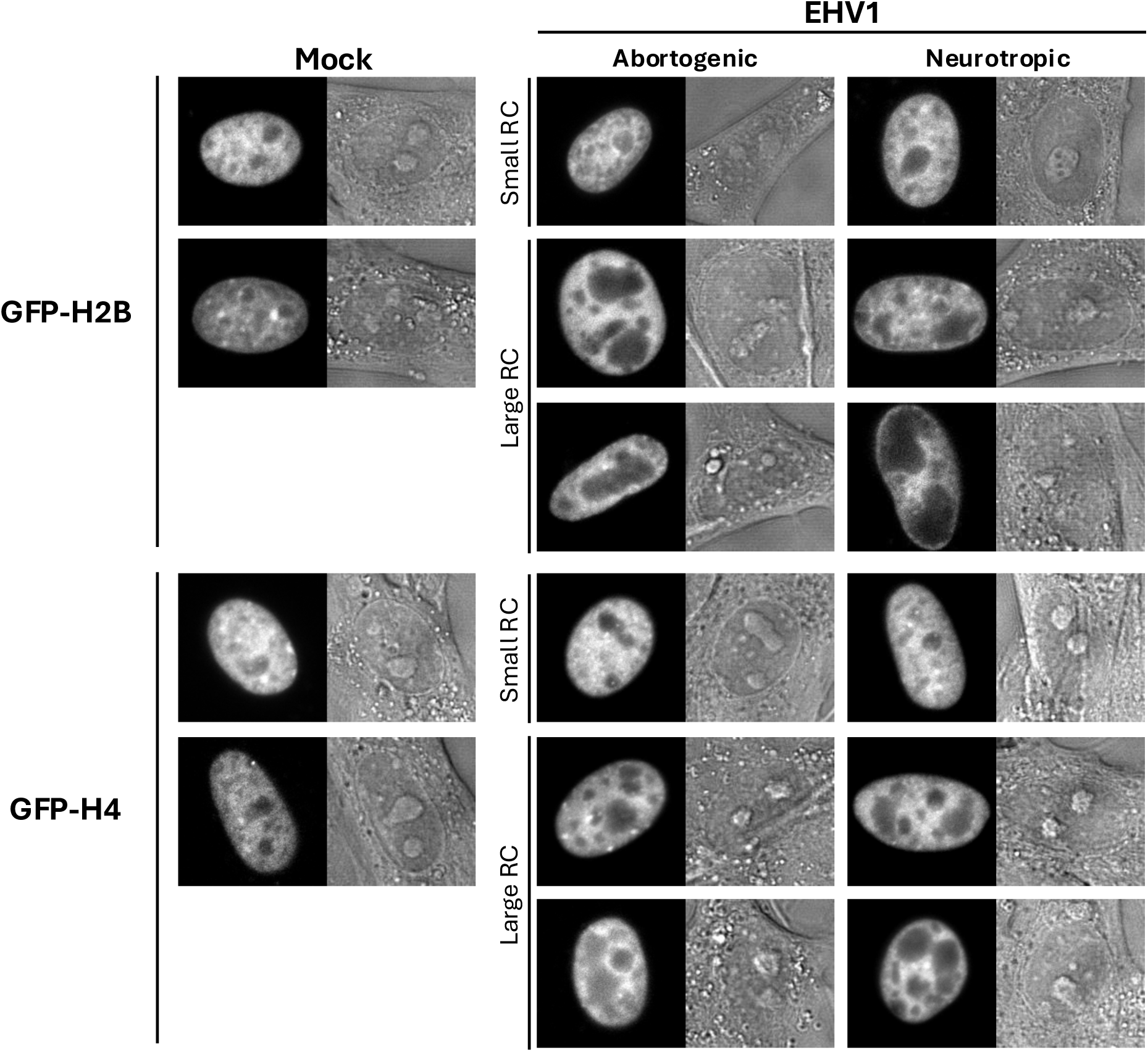
GFP-H2B or –H4 fusion proteins are depleted from EHV1 replication compartments. Digital fluorescent (left panels) and differential interference contrast (DIC; right panels) micrographs show the nucleus of EDerm cells expressing GFP-H2B or – H4. Cells were transfected with plasmids expressing GFP-H2B or –H4. At least 40h after transfection, cells were mock-infected or infected with 10 plaque forming units (PFU) per cell with an abortogenic or neurotropic field isolated EHV1 strain. Live cells were imaged between 5 and 6 hours post infection (hpi). Note the presence of small pools of GFP-H2B or –H4 within EHV1 replication compartments (RCs), similar to the reported localization of GFP-H2B or –H4 within HSV1 RCs(49). Note that nucleoli similarly have small pools of GFP-H2B or –H4 within them.

We evaluated histone dynamics at 5 hours after infection when most cells had identifiable RCs of various sizes (S1 Table). Thus, we evaluated histone mobility in a heterogeneous population of infected cells with variable levels of viral transcription, IE (immediate early), E (early), or L (late) protein expression, and DNA replication. As a surrogate measure for EHV1 chromatin dynamics, we measured histone mobility within RCs, as these nuclear domains are enriched for viral chromatin (Fig 2A). We also measured histone mobility within regions of infected-cell chromatin to test the effect, if any, of infection on cellular chromatin dynamics (Fig 2A). EHV1 RCs are highly transcriptionally active domains, therefore, as a comparator for histone dynamics within a transcriptionally active nuclear domain we measured histone mobility within mock-infected nucleoli (Fig 2A). Equal volume areas were selected for photobleaching within mock-infected nucleoli and the surrounding cell chromatin of the same mock-infected cell, or within an EHV1 RC and the surrounding infected-cell chromatin of the same EHV1-infected cell (Fig 2A). Fluorescence recovery within the photobleached regions, which represents GFP-histone mobility, was then measured over time. Most core histones are typically stably assembled in chromatin and their exchange occurs in the magnitude of hours (49–51, 57, 58, 60, 67, 68). This histone population is unlikely to undergo chromatin exchange in a timescale relevant for assembly in lytic EHV1 chromatin. However, smaller populations of core histones are, at any given time, not assembled in chromatin and available within the freely diffusing histone pool, or more transiently associated with chromatin and undergoing fast chromatin exchange (Fig 2B). We therefore measured histone mobility with a focus on these dynamic histone populations most likely available to assemble in lytic EHV1 chromatin (Fig 2B).

**Fig 2.**
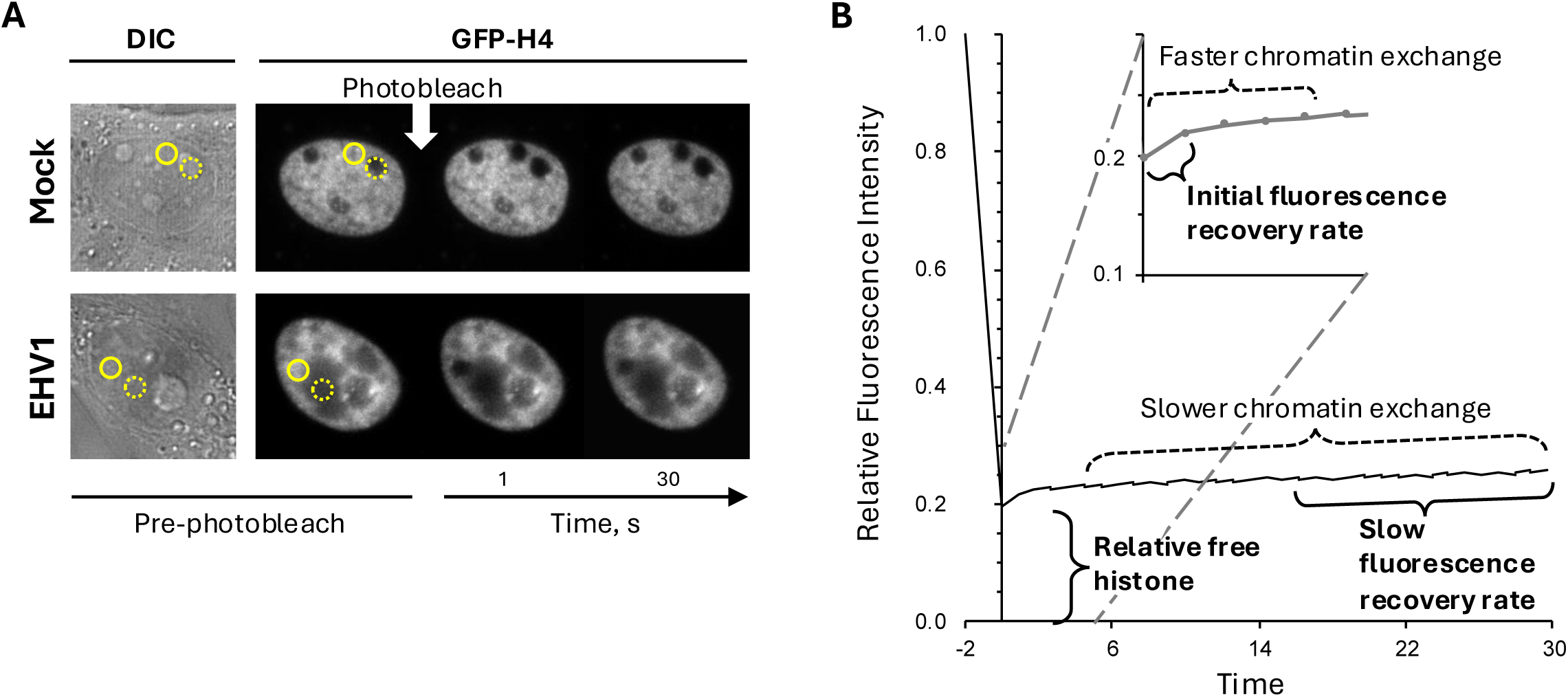
Representative FRAP of GFP-core histone fusion proteins. EDerm cells were transfected with plasmids expressing GFP fused to H4. Transfected cells were mock-infected or infected with 10 PFU/cell of EHV1 at least 40h after transfection. GFP-H4 FRAP was evaluated between 5 and 6hpi. (**A**) Digital DIC (left panel) and fluorescent (right panels) micrographs of the nucleus of cells expressing GFP-H4 before and at 1 or 30s after photobleaching. Selected equal volume regions within the cell chromatin (solid circle) and the nucleolus or EHV1 replication compartment (RC) (dashed circle) were photobleached and fluorescence recovery within the photobleached regions was measured over time. In infected cells, the region selected within the cell chromatin is enriched for cellular chromatin relative to EHV1 chromatin, whereas the region selected within the RC is enriched for EHV1 chromatin relative to cellular chromatin. Fluorescence within the photobleached regions recovers as the bleached GFP-histones within them exchange with the non-bleached fluorescent GFP-histones outside them. (**B**) Line graph of a representative GFP-H4 FRAP in the cell chromatin (an area such as that denoted by the solid circle in panel A, mock). The fluorescence intensity of the photobleached region at a given time is normalized to the fluorescence intensity of the entire nucleus at that same time, expressed relative to the normalized fluorescence intensity of the same photobleached region prior to photobleaching, and is plotted against time after photobleaching. FRAP is therefore independent of the GFP-histone expression levels within any given cell. The first data point after photobleaching is set at 0s due to differences in the time required to photobleach two regions per nucleus within each individual cell (as depicted in panel A). The first data point after photobleaching is a surrogate measure for the levels of free GFP-histone as only freely diffusing histones can move into the photobleached region in this timeframe. Subsequent fluorescent recovery is biphasic. An initial faster phase represents those histones that are weakly bound (low-affinity interactions) in chromatin and therefore undergoing fast chromatin exchange. As a surrogate measure for this histone population, we calculated the initial rate of normalized fluorescence recovery (the slope between the normalized fluorescence at the first and second data points after photobleaching; shown in the inset). The second slower phase of fluorescence recovery represents those histones that are more stably bound (high-affinity interactions) in chromatin and undergoing slow chromatin exchange. As a surrogate measure for this histone population, we calculated the slow rate of fluorescence recovery (the slope between the normalized fluorescence at the 15 and 30s data points after photobleaching). Those histones available in the free pool or undergoing fast chromatin exchange (weakly bound in chromatin) represent the histone populations that are most likely available for assembly in EHV1 chromatin in a timescale relevant to lytic infection.

GFP-H4 fluorescence recovered faster within mock-infected nucleoli than within the surrounding cellular chromatin (Fig 3). Thus, as expected, GFP-H4 was more mobile within nucleoli where the highly transcribed and unstable nucleolar chromatin is. Infection with abortogenic or neurotropic EHV1 mobilized GFP-H4 such that fluorescence recovered faster in infected-than in mock-infected cell chromatin (Fig 3). Moreover, GFP-H4 fluorescence recovered even faster within the EHV1 RCs than within the surrounding infected-cell chromatin (Fig 3). These data show that EHV1 infection globally mobilized H4 and mobilized it to a greater degree in nuclear domains enriched for viral chromatin. H4 was more mobile within RCs than within mock-infected nucleoli, suggesting that H4 interactions in EHV1 chromatin are more dynamic or unstable than its interactions in nucleolar chromatin.

**Fig 3.**
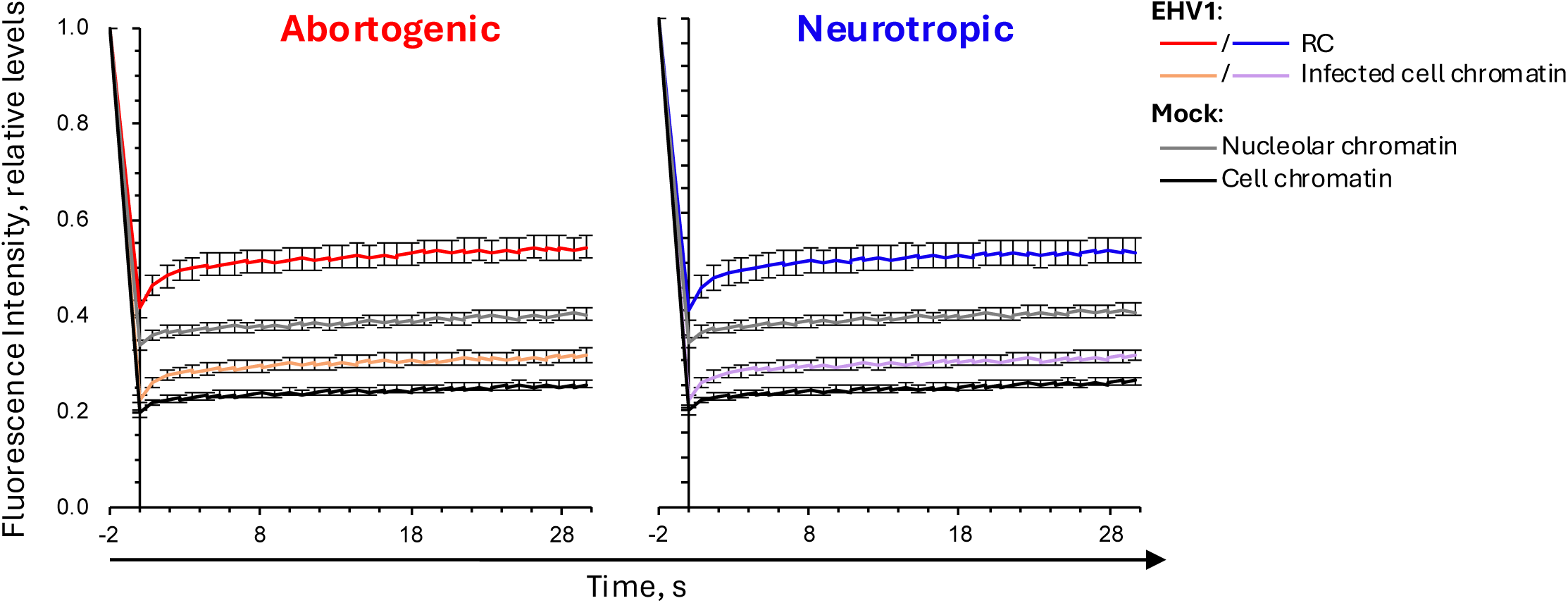
GFP-H4 fluorescence recovers faster in EHV1-infected than in mock-infected cells. EDerm cells were transfected with plasmids encoding GFP-H4. At least 40h after transfection, cells were mock infected (squares) or infected (circles) with 10 PFU/cell of abortogenic or neurotropic EHV1, as indicated. The mobility of GFP-H4 was evaluated between 5 and 6hpi by FRAP. Line graphs present the normalized fluorescence intensities of the photobleached nuclear regions expressed as a ratio to their normalized fluorescence intensities prior to photobleaching plotted against time after photobleaching. FRAP of GFP-H4 in mock-infected cell-or nucleolar-chromatin are plotted in both graphs for comparison. Error bars, standard errors of the means (SEM); n≥39 cells per treatment from 4 independent experiments.

GFP-H2B or –H4 were similarly mobilized during infection with abortogenic or neurotropic EHV1 (Fig 3; data not shown). Given the heterogeneity of infection progression with field isolated strains, we considered that the levels of EHV1 transcription, protein expression, DNA replication, or cellular responses to them, may affect H2B or H4 mobility. We therefore pooled the infected cell FRAP data for each histone and then segregated it by the presence or absence of identifiable RCs rather than by infecting strain. Cells with GFP-H2B or –H4-depleted regions identifiable as RCs by shape or size were grouped as “large” RC, whereas cells with depleted regions that more resembled mock-infected cells were grouped as “small” RC (for examples see Fig 1). Seventy-six or 70% of GFP-H2B expressing cells or 77 or 67% of GFP-H4 expressing ones infected with abortogenic or neurotropic EHV1, respectively, had identifiable “large” RCs (S1 Table).

Fluorescence recovery of GFP-H2B or –H4 was still enhanced within “small” RCs, although fluorescence recovered slower within these domains than within mock infected nucleoli (Fig 4A). Fluorescence recovery in the surrounding infected-cell chromatin was also slower, and similar to that within the mock-infected cell chromatin (Fig 4A). GFP-H2B or –H4 fluorescence recovery was most enhanced within the “large” RCs, where it recovered much faster than within mock-infected nucleoli (Fig 4B). Mobilization of H2B or H4 in cells with “large” RCs also mobilized these histones within the surrounding infected-cell chromatin such that fluorescence recovered faster than it did within the mock-infected cell chromatin (Fig 4B).

**Fig 4.**
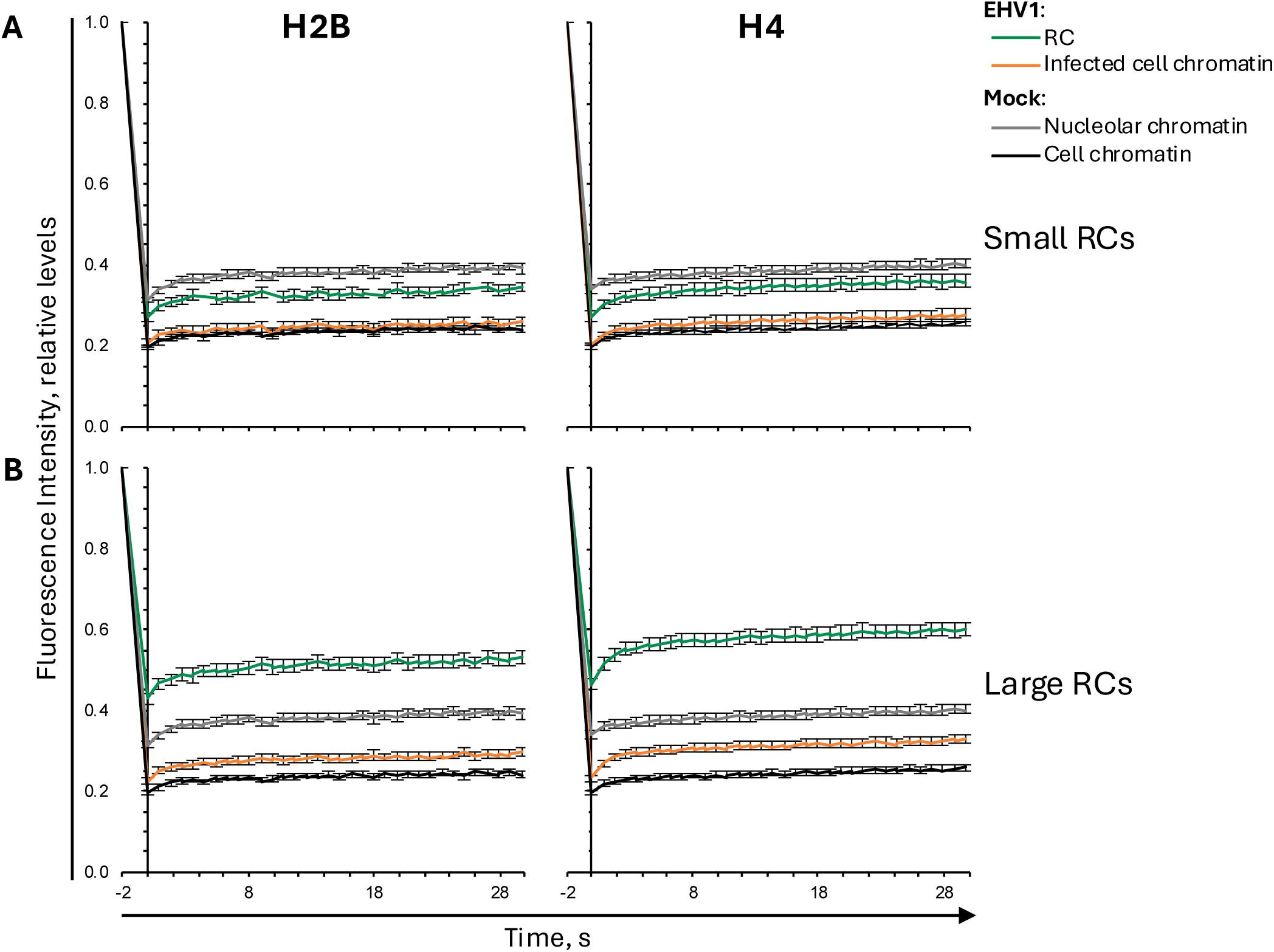
H2B or H4 are most mobile within EHV1 RCs. EDerm cells were transfected with plasmids encoding GFP-H2B or –H4. Transfected cells were mock-infected or infected with 10 PFU/cell of abortogenic or neurotropic EHV1 at least 40h after transfection. Nuclear mobility of GFP-H2B or –H4 was evaluated by FRAP between 5 and 6hpi. FRAP data for EHV1 infected cells were pooled for each histone and segregated by the absence (A; Small RC) or presence (B; Large RC) of clearly identifiable RCs. Line graphs present GFP-H2B or –H4 FRAP in the RCs or infected-cell chromatin of EHV1-infected cells, or the nucleolar-or cell-chromatin of mock-infected cells. GFP-H4 FRAP data presented in Figure 3 is re-analyzed and re-plotted. FRAP of GFP-H2B or –H4 in the nucleolar-or cell-chromatin of mock-infected cells are plotted in both graphs for comparison. Error bars, SEM; n≥38 cells per treatment from 4 independent experiments.

These data show that EHV1 mobilized H2B and H4. These histones were more mobile within RCs, illustrating that H2B and H4 were most dynamic within nuclear domains enriched for viral chromatin. H2B and H4 were mobilized to a greater degree in RCs of cells in which infection was further progressed. Therefore, increased levels of EHV1 transcription, IE, E, or L protein expression, or DNA replication may directly or indirectly enhance H2B and H4 mobility. Within “large” RCs, H2B and H4 were more mobile than within mock-infected nucleoli, suggesting that EHV1 chromatin may be more dynamic or unstable than nucleolar chromatin.

### The most dynamic H2B and H4 populations are mobilized by EHV1

To investigate the H2B and H4 histone populations mobilized during EHV1 infection, we examined GFP-H2B and –H4 fluorescence recovery kinetics. As a surrogate measure for the histones not assembled in chromatin and freely diffusing in the nucleoplasm (the “free” pool), we used the first value after photobleaching (Fig 2B). Only freely diffusing histones can move into photobleached regions within this time frame. In mock-infected cells, nucleolar H2B or H4 free pools were 160 ± 4% or 172 ± 5%, respectively, of the free pools within the surrounding cellular chromatin, consistent with the dynamic instability of nucleolar chromatin (*P*<0.01; Fig 5A, Table 1). Within infected cells, H2B or H4 free pools tended to increase within domains enriched for infected-cell chromatin, to 111 ± 3% or 114 ± 3%, respectively, of their levels within the mock-infected cell chromatin (*P*=ns or <0.01, respectively; Fig 5A, Table 1). However, H2B or H4 free pools increased to a greater degree in domains enriched for viral chromatin. In “small” RCs, free H2B or H4 levels increased to 133 ± 6% or 144 ± 8%, respectively, relative to mock-infected cell chromatin (*P*<0.01; Fig 5A, Table 1). Free H2B or H4 pools increased even further within “large” RCs, to 217 ± 6% or 234 ± 6%, respectively (*P*<0.01; Figure 5A, Table 1). The levels of free H2B or H4 within “large” RCs were significantly greater than within mock-infected nucleoli, further supporting that H2B and H4 are most dynamic within “large” EHV1 RCs (*P*<0.01; Fig 5A, Table 1).

**Fig 5.**
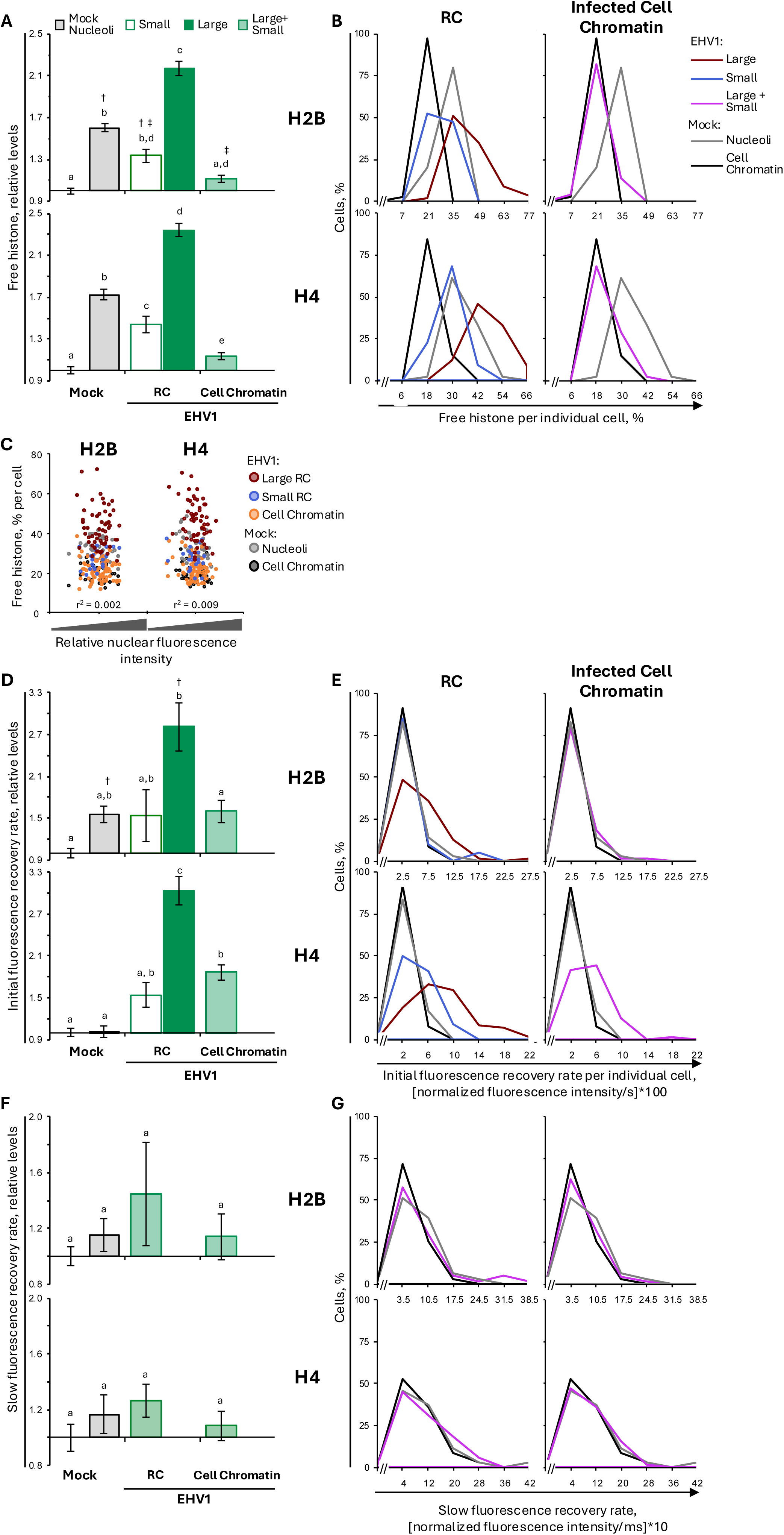
EHV1 mobilizes unbound and weakly bound H2B or H4 within RCs. EDerm cells were transfected with plasmids encoding GFP-H2B or –H4. At least 40h post transfection, cells were mock-infected or infected with 10 PFU/cell of abortogenic or neurotropic EHV1. Nuclear mobilities of GFP-H2B or –H4 were evaluated by FRAP between 5 and 6hpi. FRAP data for EHV1 infected cells were pooled for each histone and segregated by the absence (Small) or presence (Large) of clearly identifiable RCs. (**A**) Bar graphs present the average normalized levels of GFP-H2B or –H4 in the free pools expressed as a ratio to the average normalized level in mock-infected cell chromatin (set at 1). (**B**) Frequency distribution plots show the percentage of free GFP-H2B or –H4 per individual cell. Solid vertical black line indicates one standard deviation (SD) above the average level of free GFP-H2B or –H4 in mock-infected cell chromatin; dashed vertical black line indicates 1 SD above the average level of free GFP-H2B or –H4 in mock-infected nucleolar chromatin. The levels of free GFP-H2B or –H4 per individual cell in the mock-infected cell-or nucleolar-chromatin are plotted in both graphs for comparison. (**C**) Dot plots present the level of GFP-H2B or –H4 in the free pools of individual cells plotted against their normalized total nuclear fluorescence intensity prior to photobleaching. (**D**) Bar graphs present the average initial normalized fluorescence recovery rate for GFP-H2B or –H4 expressed as a ratio to the average initial normalized fluorescence recovery rate in mock-infected cell chromatin (set at 1). (**E**) Frequency distribution plots present the initial normalized fluorescence recovery rate of GFP-H2B or –H4 per individual cell. Solid or dashed vertical black lines, 1 SD above the average initial normalized fluorescence recovery rate for GFP-H2B or –H4 in mock-infected cell-or nucleolar-chromatin, respectively. The initial normalized fluorescence recovery rate for GFP-H2B or –H4 per individual cell in the mock-infected cell-or nucleolar-chromatin are plotted in both graphs for comparison. (**F**) Bar graphs represent the average slow normalized fluorescence recovery rate for GFP-H2B or –H4 expressed as a ratio to the average slow normalized fluorescence recovery rate in mock-infected cell chromatin (set at 1). (**G**) Frequency distribution plots show the slow normalized fluorescence recovery rate for GFP-H2B or –H4 per individual cell. Solid or dashed vertical black lines, 1 SD above the average slow normalized fluorescence recovery rate for GFP-H2B or –H4 in mock-infected cell-or nucleolar-chromatin, respectively. The slow normalized fluorescence recovery rate for GFP-H2B or –H4 in mock-infected cell-or nucleolar-chromatin are re-plotted in both graphs for comparison. Error bars, SEM. n≥38 cells per treatment from 4 independent experiments. Different letters denote *P*<0.01; matching symbols denote *P*<0.05. Statistical significance evaluated by ANOVA with post-hoc Tukey Kramer pair-wise analysis.

**Table 1.**
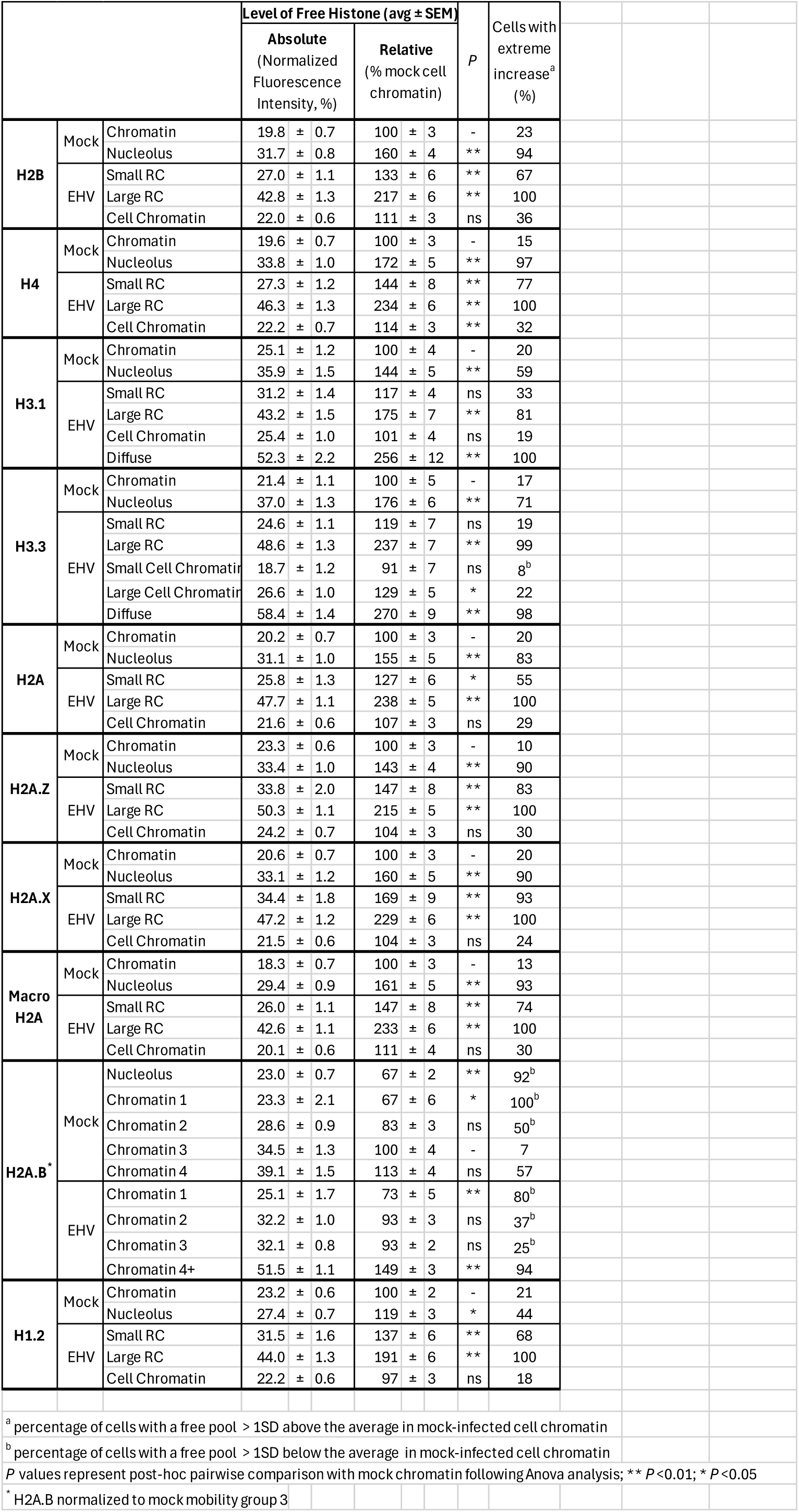
Free histone levels.

It was possible that a subpopulation of infected cells increased their H2B or H4 free pools by an extreme degree while others had little to no change. We therefore evaluated the level of free H2B or H4 per individual cell. The frequency distribution of free H2B or H4 within the cell chromatin of each individual EHV1-or mock-infected cell were unimodal and largely overlapped, consistent with the similar average levels of H2B or H4 in their free pools (Fig 5A, B). The frequency distribution of free H2B or H4 within “small” or “large” RCs per individual cell were also unimodal, indicating that H2B and H4 free pools increased within RCs throughout the infected cell population (Fig 5B). Consistent with the average increases in H2B or H4 free pools within “small” or “large” RCs, their frequency distributions shifted to the right relative to that of mock-infected cell chromatin (Fig 5B). Most of the “small” RCs evaluated had free H2B or H4 levels greater than one standard deviation (SD) above their average levels within mock-infected cell chromatin (67 or 77% of cells, respectively; Fig 5B, Table 1). The free pools of H2B or H4 increased by an extreme degree (>1SD above the mock-infected cell chromatin average) in all evaluated “large” RCs (Fig 5B, Table 1). Furthermore, 70% or 68% of cells had H2B or H4 free pools within “large” RCs greater than 1SD above their average levels within mock-infected nucleoli.

To ensure that H2B or H4 free pools were not increased due to GFP-H2B or –H4 expression levels, the level of free histone per individual cell was plotted relative to its total nuclear fluorescence intensity prior to photobleaching (Fig 5C). Free H2B or H4 levels in any evaluated nuclear domain of either mock-or EHV1-infected cells did not correlate with total nuclear fluorescence. Thus, H2B or H4 free pools within nuclear domains enriched for viral, nucleolar, or cellular chromatin were independent of the overall GFP-histone expression levels.

EHV1 mobilization of H2B or H4 caused a net increase in their free pools within RCs and the H4 free pool within the infected-cell chromatin. To identify whether infection also affected H2B or H4 low-affinity chromatin turnover (fast chromatin exchange) we next evaluated their initial normalized fluorescence recovery rates (Fig 2B). H2B or H4 fast chromatin exchange tended to increase within “small” RCs (to 153 ± 37% or 154 ± 18%, respectively), although these rates were not significantly different from those within mock-infected cell chromatin (Fig 5D, Table 2). Nonetheless, 30 or 55% of cells had an extreme increase in H2B or H4 fast chromatin exchange within “small” RCs (>1SD higher than the mock-infected cell chromatin average), which is larger than expected in a normal population were they not mobilized (Fig 5E, Table 2). The fast chromatin exchange of H2B or H4 further increased within “large” RCs (to 281 ± 35% or 304 ± 20%, respectively; *P*<0.01; Fig 5D, Table 2). H2B or H4 fast chromatin exchange increased within “large” RCs throughout the infected cell population, with an extreme increase (>1SD above the mock-infected cell chromatin average) in 56 or 84% of cells, respectively (Fig 5E, Table 2). H4 mobilization within EHV1-infected cells also affected its fast chromatin exchange within the surrounding infected-cell chromatin, where it increased to 187 ± 11% (*P*<0.01; Fig 5D, E, Table 2).

**Table 2.** Initial normalized rates of fluorescence recovery.

|  |  |  | Initial Fluorescence Recovery Rates (avg ± SEM) |  | P | Cells with extreme increase<sup>a</sup> (%) |
| --- | --- | --- | --- | --- | --- | --- |
|  |  |  | Absolute (Normalized Fluorescence Intensity/ms) | Relative (% mock cell chromatin) |  |  |
| H2B | Mock | Chromatin | 27.5 ± 2.5 | 100 ± 7 | - | 11 |
|  |  | Nucleolus | 42.3 ± 4.0 | 156 ± 12 | ns | 29 |
|  | EHV | Small RC | 41.3 ± 6.4 | 153 ± 37 | ns | 30 |
|  |  | Large RC | 61.9 ± 61.9 | 281 ± 35 | ** | 56 |
|  |  | Cell Chromatin | 36.7 ± 2.9 | 160 ± 16 | ns | 28 |
| H4 | Mock | Chromatin | 27.5 ± 1.7 | 100 ± 6 | - | 16 |
|  |  | Nucleolus | 27.9 ± 2.2 | 101 ± 8 | ns | 19 |
|  | EHV | Small RC | 40.8 ± 4.5 | 154 ± 18 | ns | 55 |
|  |  | Large RC | 84.5 ± 6.0 | 304 ± 20 | ** | 84 |
|  |  | Cell Chromatin | 51.4 ± 3.3 | 187 ± 11 | ** | 65 |
| H3.1 | Mock | Chromatin | 73.9 ± 8.9 | 100 ± 10 | - | 13 |
|  |  | Nucleolus | 84.0 ± 10.5 | 110 ± 10 | ns | 15 |
|  | EHV | Small RC | 75.7 ± 12.3 | 85 ± 15 | ns | 7 |
|  |  | Large RC | 90.0 ± 8.2 | 155 ± 18 | ns | 25 |
|  |  | Cell Chromatin | 61.8 ± 6.2 | 94 ± 10 | ns | 12 |
|  |  | Diffuse | 215.3 ± 19.1 | 373 ± 36 | ** | 82 |
| H3.3 | Mock | Chromatin | 44.3 ± 7.1 | 100 ± 11 | - | 10 |
|  |  | Nucleolus | 46.6 ± 5.5 | 118 ± 14 | ns | 14 |
|  | EHV | Small RC | 39.7 ± 6.4 | 162 ± 39 | ns | 13 |
|  |  | Large RC | 75.7 ± 7.7 | 260 ± 40 | * | 23 |
|  |  | Cell Chromatin | 49.8 ± 4.6 | 157 ± 18 | ns | 11 |
|  |  | Diffuse | 194.7 ± 11.5 | 502 ± 40 | ** | 89 |
| H2A | Mock | Chromatin | 24.3 ± 2.0 | 100 ± 5 | - | 11 |
|  |  | Nucleolus | 37.3 ± 2.9 | 162 ± 10 | ns | 36 |
|  | EHV | Small RC | 31.1 ± 4.6 | 110 ± 20 | ns | 32 |
|  |  | Large RC | 72.9 ± 5.6 | 357 ± 27 | ** | 82 |
|  |  | Small Cell Chromatin | 27.6 ± 2.4 | 99 ± 17 | ns | 25 |
|  |  | Large Cell Chromatin | 40.8 ± 2.2 | 202 ± 12 | * | 44 |
| H2A.Z | Mock | Chromatin | 35.0 ± 2.9 | 100 ± 8 | - | 18 |
|  |  | Nucleolus | 37.7 ± 2.8 | 111 ± 9 | ns | 13 |
|  | EHV | Small RC | 53.5 ± 8.2 | 154 ± 28 | ns | 45 |
|  |  | Large RC | 85.0 ± 6.8 | 263 ± 25 | ** | 70 |
|  |  | Cell Chromatin | 49.6 ± 3.4 | 152 ± 13 | ns | 32 |
| H2A.X | Mock | Chromatin | 24.9 ± 1.8 | 100 ± 6 | - | 15 |
|  |  | Nucleolus | 34.5 ± 2.8 | 137 ± 10 | ns | 43 |
|  | EHV | Small RC | 42.1 ± 8.9 | 189 ± 49 | ns | 36 |
|  |  | Large RC | 48.4 ± 4.1 | 212 ± 14 | ** | 58 |
|  |  | Cell Chromatin | 31.8 ± 1.8 | 136 ± 9 | ns | 25 |
| Macro H2A | Mock | Chromatin | 23.1 ± 2.4 | 100 ± 9 | - | 10 |
|  |  | Nucleolus | 28.2 ± 2.6 | 123 ± 11 | ns | 16 |
|  | EHV | Small RC | 26.8 ± 2.5 | 119 ± 13 | ns | 8 |
|  |  | Large RC | 46.9 ± 5.7 | 210 ± 26 | ** | 48 |
|  |  | Cell Chromatin | 26.3 ± 1.8 | 116 ± 9 | ns | 20 |
| H2A.B<sup>*</sup> | Mock | Nucleolus | 211.0 ± 10.9 | 97 ± 5 | ns | 15 |
|  |  | Chromatin 1 | 90.3 ± 13.0 | 42 ± 6 | ** | 100<sup>b</sup> |
|  |  | Chromatin 2 | 176.9 ± 20.0 | 81 ± 9 | ns | 33<sup>b</sup> |
|  |  | Chromatin 3 | 217.3 ± 20.2 | 100 ± 9 | - | 13 |
|  |  | Chromatin 4 | 350.6 ± 31.8 | 161 ± 15 | ** | 71 |
|  | EHV | Chromatin 1 | 85.0 ± 7.1 | 39 ± 3 | ** | 100<sup>b</sup> |
|  |  | Chromatin 2 | 130.1 ± 6.0 | 60 ± 3 | ** | 65<sup>b</sup> |
|  |  | Chromatin 3 | 132.3 ± 6.1 | 61 ± 3 | ** | 66<sup>b</sup> |
|  |  | Chromatin 4+ | 187.5 ± 6.1 | 86 ± 3 | ns | 8<sup>b</sup> |
| H1.2 | Mock | Chromatin | 42.0 ± 1.6 | 100 ± 4 | - | 18 |
|  |  | Nucleolus | 41.4 ± 2.4 | 99 ± 6 | ns | 13 |
|  | EHV | Small RC | 59.6 ± 3.7 | 142 ± 9 | ** | 60 |
|  |  | Large RC | 57.5 ± 4.5 | 138 ± 11 | ** | 57 |
|  |  | Cell Chromatin | 44.0 ± 1.7 | 106 ± 4 | ns | 25 |
| <sup>a</sup> percentage of cells with an initial fluorescence recovery rate > 1SD above the average in mock-infected cell chromatin |  |  |  |  |  |  |
| <sup>b</sup> percentage of cells with an initial fluorescence recovery rate >1SD below the average in mock-infected cell chromatin |  |  |  |  |  |  |
| P values represent post-hoc pairwise comparison with mock chromatin following Anova analysis; ** P<0.01; * P<0.05 |  |  |  |  |  |  |
| <sup>*</sup> H2A.B normalized to mock mobility group 3 |  |  |  |  |  |  |

To investigate whether EHV1 also mobilized those histones more stably assembled in chromatin, we next evaluated slow normalized fluorescence recovery (Fig 2B). This analysis revealed that the high-affinity (stable) chromatin interactions of H2B and H4 were largely unaffected during infection. The H2B or H4 slow chromatin exchange rates tended to increase within RCs (to 144 ± 16% or 127 ± 12%, respectively), however, these rates were not significantly different from those within mock-infected cell chromatin (Fig 5F, G, Table 3). Thus, EHV1 infection primarily mobilized H2B and H4 by increasing their low-affinity chromatin turnover and free pools. These most mobile histone populations would be available to assemble in, and exchange with, EHV1 chromatin on a timescale relevant to lytic infection.

**Table 3.**
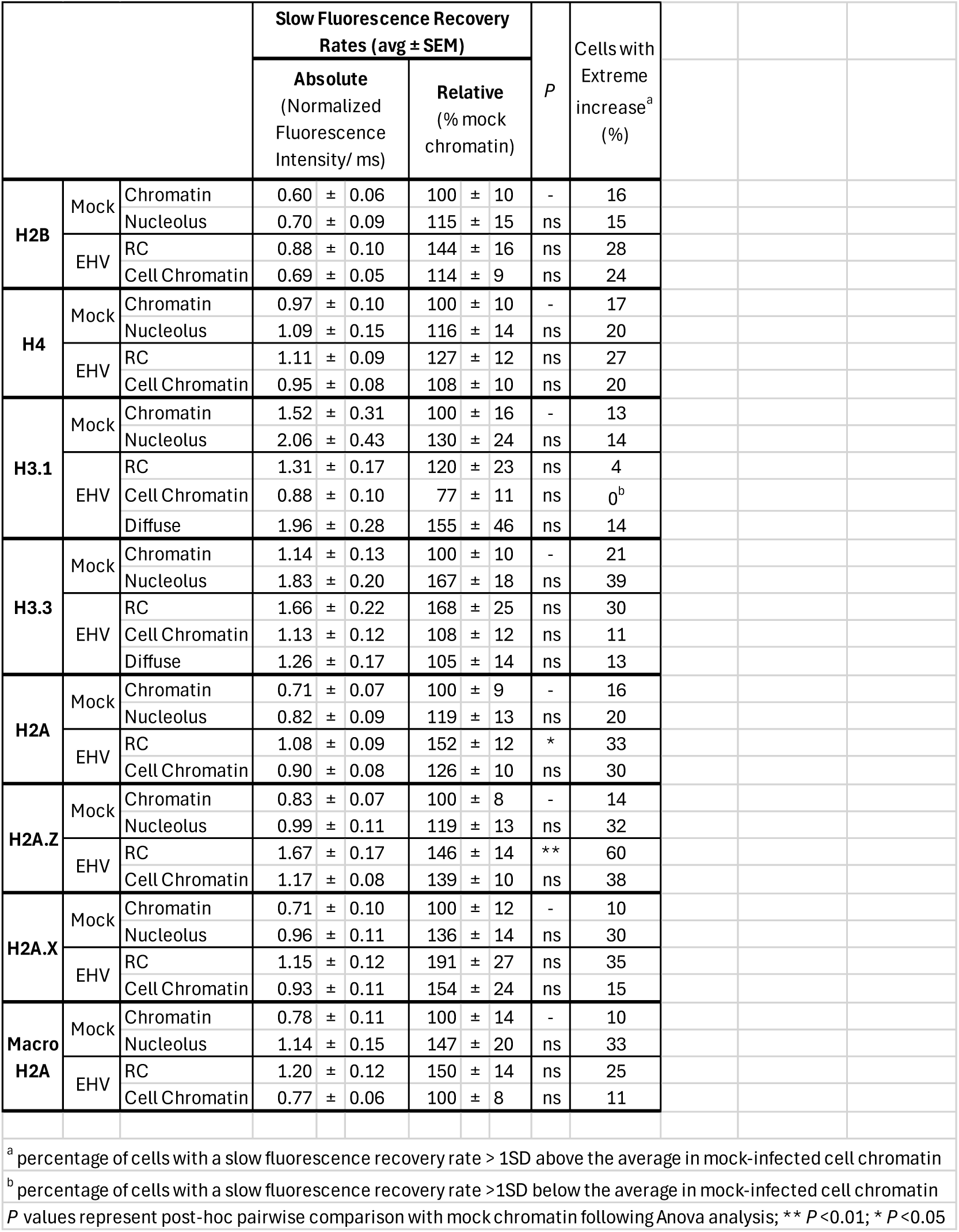
Normalized rates of slow fluorescence recovery.

|  |  |  | Slow Fluorescence Recovery Rates (avg ± SEM) |  | P | Cells with Extreme increase<sup>a</sup> (%) |
| --- | --- | --- | --- | --- | --- | --- |
|  |  |  | Absolute (Normalized Fluorescence Intensity/ ms) | Relative (% mock chromatin) |  |  |
| H2B | Mock | Chromatin | 0.60 ± 0.06 | 100 ± 10 | - | 16 |
|  |  | Nucleolus | 0.70 ± 0.09 | 115 ± 15 | ns | 15 |
|  | EHV | RC | 0.88 ± 0.10 | 144 ± 16 | ns | 28 |
|  |  | Cell Chromatin | 0.69 ± 0.05 | 114 ± 9 | ns | 24 |
| H4 | Mock | Chromatin | 0.97 ± 0.10 | 100 ± 10 | - | 17 |
|  |  | Nucleolus | 1.09 ± 0.15 | 116 ± 14 | ns | 20 |
|  | EHV | RC | 1.11 ± 0.09 | 127 ± 12 | ns | 27 |
|  |  | Cell Chromatin | 0.95 ± 0.08 | 108 ± 10 | ns | 20 |
| H3.1 | Mock | Chromatin | 1.52 ± 0.31 | 100 ± 16 | - | 13 |
|  |  | Nucleolus | 2.06 ± 0.43 | 130 ± 24 | ns | 14 |
|  | EHV | RC | 1.31 ± 0.17 | 120 ± 23 | ns | 4 |
|  |  | Cell Chromatin | 0.88 ± 0.10 | 77 ± 11 | ns | 0<sup>b</sup> |
|  |  | Diffuse | 1.96 ± 0.28 | 155 ± 46 | ns | 14 |
| H3.3 | Mock | Chromatin | 1.14 ± 0.13 | 100 ± 10 | - | 21 |
|  |  | Nucleolus | 1.83 ± 0.20 | 167 ± 18 | ns | 39 |
|  | EHV | RC | 1.66 ± 0.22 | 168 ± 25 | ns | 30 |
|  |  | Cell Chromatin | 1.13 ± 0.12 | 108 ± 12 | ns | 11 |
|  |  | Diffuse | 1.26 ± 0.17 | 105 ± 14 | ns | 13 |
| H2A | Mock | Chromatin | 0.71 ± 0.07 | 100 ± 9 | - | 16 |
|  |  | Nucleolus | 0.82 ± 0.09 | 119 ± 13 | ns | 20 |
|  | EHV | RC | 1.08 ± 0.09 | 152 ± 12 | * | 33 |
|  |  | Cell Chromatin | 0.90 ± 0.08 | 126 ± 10 | ns | 30 |
| H2A.Z | Mock | Chromatin | 0.83 ± 0.07 | 100 ± 8 | - | 14 |
|  |  | Nucleolus | 0.99 ± 0.11 | 119 ± 13 | ns | 32 |
|  | EHV | RC | 1.67 ± 0.17 | 146 ± 14 | ** | 60 |
|  |  | Cell Chromatin | 1.17 ± 0.08 | 139 ± 10 | ns | 38 |
| H2A.X | Mock | Chromatin | 0.71 ± 0.10 | 100 ± 12 | - | 10 |
|  |  | Nucleolus | 0.96 ± 0.11 | 136 ± 14 | ns | 30 |
|  | EHV | RC | 1.15 ± 0.12 | 191 ± 27 | ns | 35 |
|  |  | Cell Chromatin | 0.93 ± 0.11 | 154 ± 24 | ns | 15 |
| Macro H2A | Mock | Chromatin | 0.78 ± 0.11 | 100 ± 14 | - | 10 |
|  |  | Nucleolus | 1.14 ± 0.15 | 147 ± 20 | ns | 33 |
|  | EHV | RC | 1.20 ± 0.12 | 150 ± 14 | ns | 25 |
|  |  | Cell Chromatin | 0.77 ± 0.06 | 100 ± 8 | ns | 11 |

<sup>a</sup> percentage of cells with a slow fluorescence recovery rate > 1SD above the average in mock-infected cell chromatin <sup>b</sup> percentage of cells with a slow fluorescence recovery rate >1SD below the average in mock-infected cell chromatin P values represent post-hoc pairwise comparison with mock chromatin following Anova analysis; \*\* P<0.01; \* P<0.05

Together these data show that H2B and H4 were mobilized to the greatest degree within nuclear domains enriched for EHV1 chromatin. Mobilization altered H2B and H4 chromatin residency to cause a net increase in their free pools and increased their rates of low-affinity chromatin exchange. H2B and H4 were mobilized to a greater degree within the RCs of cells in which infection had further progressed, suggesting that higher levels of EHV1 DNA replication, transcription, protein expression, or cellular responses to them further mobilize H2B or H4. Infection also mobilized H4, but not H2B, within the infected-cell chromatin by increasing its free pool and low-affinity turnover. EHV1 did not significantly mobilize populations of H2B or H4 more stably bound in chromatin and undergoing slow chromatin exchange. Thus, EHV1 further mobilized the dynamic histone populations most likely available to assemble in lytic viral chromatin.

### EHV1 mobilizes canonical H3.1 and variant H3.3 histones

We next measured canonical H3.1 and variant H3.3 mobility to test whether EHV1 differentially mobilizes particular histone types (as does HSV1) (50, 60, 61). GFP-H3.1 or – H3.3 had discrete granular localization consistent with chromatin regions relatively enriched or depleted for either histone and they were relatively depleted from nucleoli (Figure 6). Due to less pronounced H3.1 depletion from nucleoli and other nuclear domains, it generally appeared more diffuse than H3.3 (Fig 6). In most infected cells, H3.1 or H3.3 were largely depleted from EHV1 RCs. However, some cells had less pronounced depletion of GFP-H3.1 or –H3.3 from “large” RCs relative to the surrounding infected-cell chromatin. Moreover, a minor subpopulation of cells had diffuse GFP-H3.1 or –H3.3, with relative depletion of either histone only apparent from nucleoli (Fig 6 EHV1 “Diffuse”; S1 Table).

**Fig 6.**
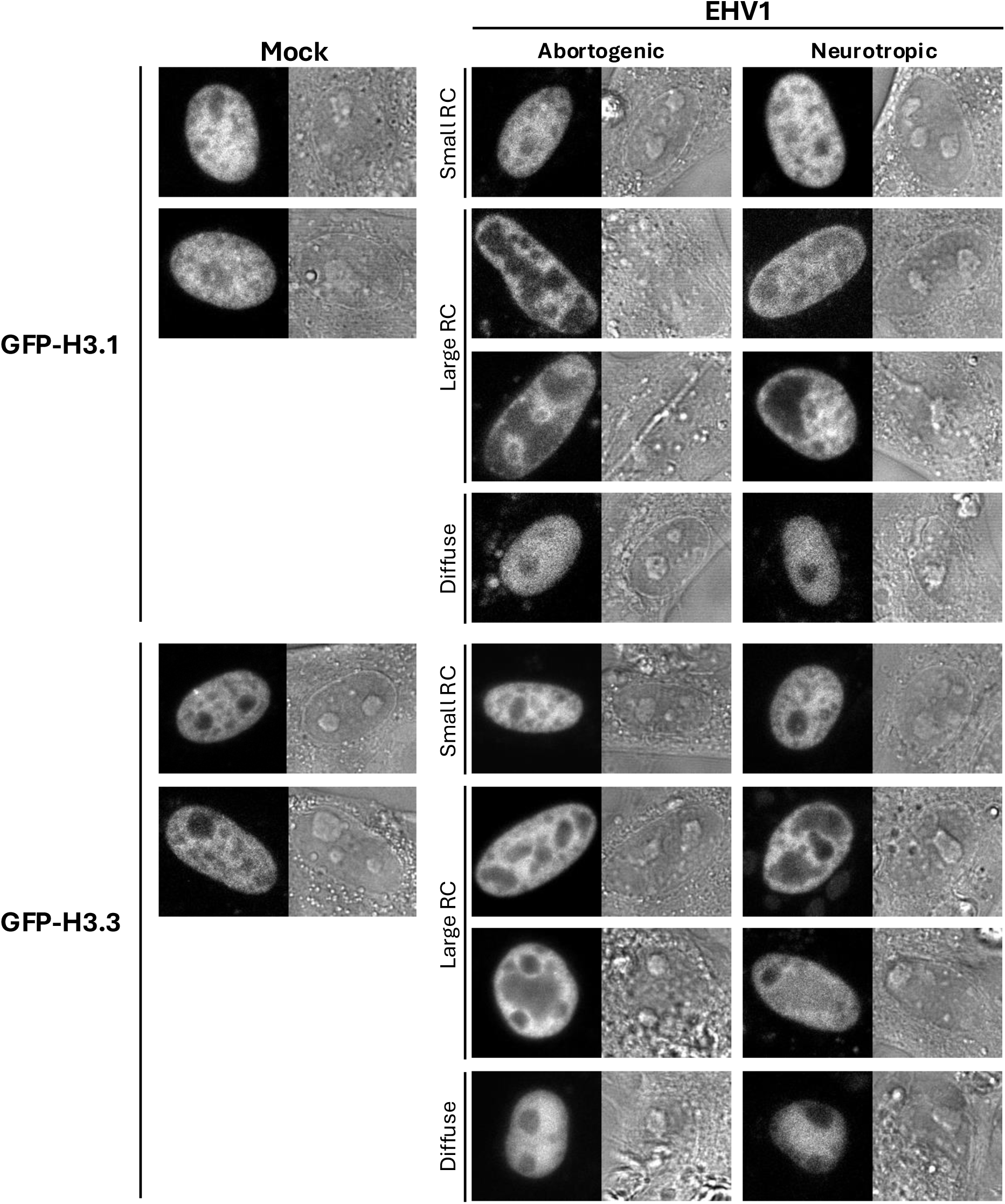
GFP-H3.1 or –H3.3 are variably depleted from EHV1 RCs. Digital fluorescent (left panels) and DIC (right panels) micrographs show the nucleus of EDerm cells expressing GFP-H3.1 or –H3.3. Cells were transfected with plasmids encoding GFP fused to H3.1 or H3.3 and, at least 40h after transfection, were mock-infected or infected with 10 PFU/cell of abortogenic or neurotropic EHV1. Live cells were imaged between 5 and 6hpi.

EHV1 infection mobilized H3.1 and H3.3 within RCs, where the viral chromatin is enriched (S2 Fig). Free H3.1 or H3.3 pools increased to 175 ± 7% or 237 ± 7%, respectively, within “large” RCs (*P*<0.01; Fig 7A, Table 1). H3.1 or H3.3 free pools increased within “large” RCs throughout the cell population, with an extreme increase (>1SD above the mock-infected cell chromatin average) in 81 or 99% of cells, respectively (Fig 7A, B, Table 1). Moreover, H3.1 or H3.3 free pools within “large” RCs were significantly greater than those within mock-infected nucleoli (144 ± 5% or 176 ± 6%, respectively; *P*<0.01), and 34 or 52% of cells had free pools more than 1SD above the average levels within mock-infected nucleoli (Fig 7A, B, Table 1). Such high levels of free H3.1 or H3.3 within “large” RCs were not consequent to GFP-histone expression levels (Fig 7C).

**Fig 7.**
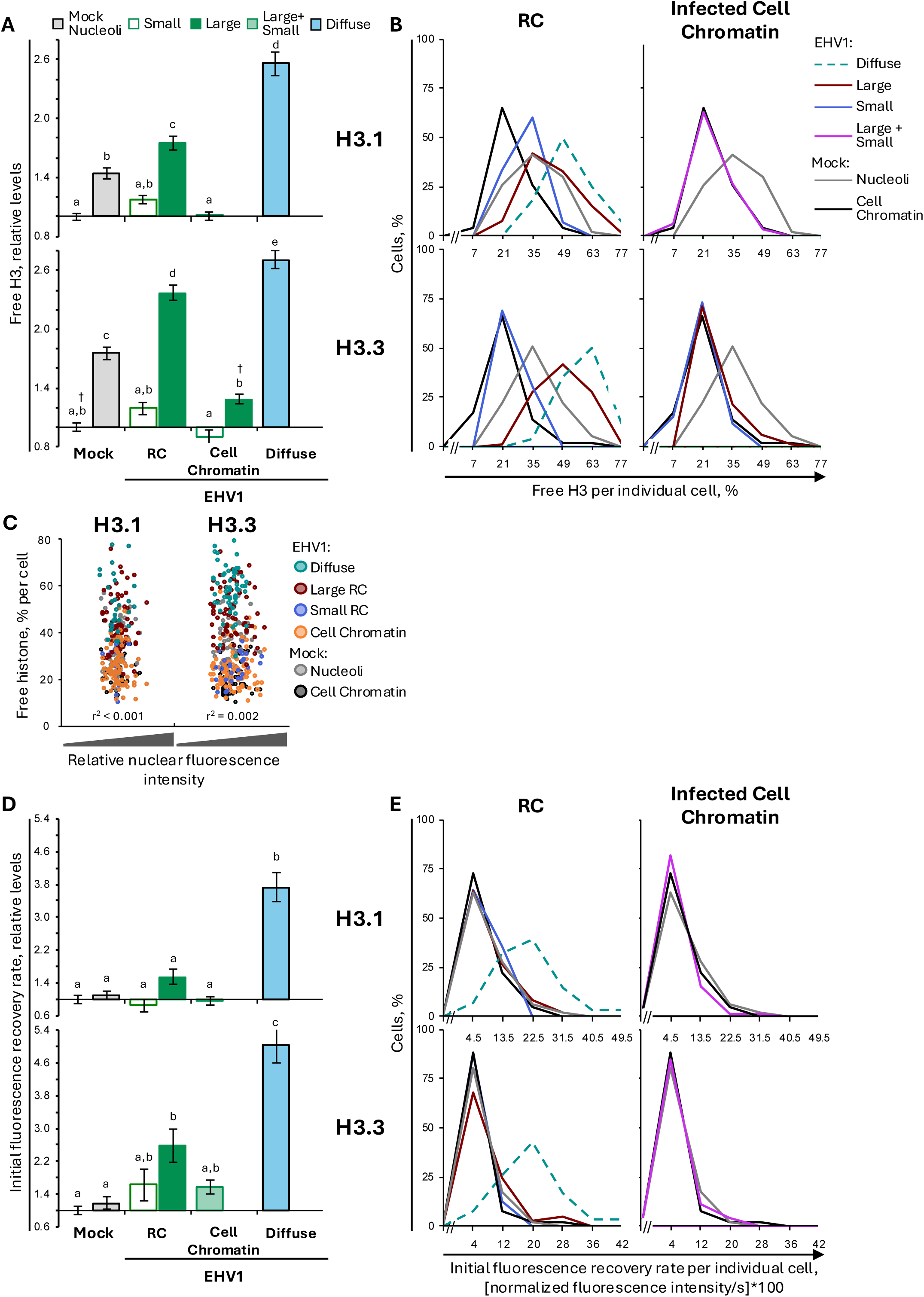
H3.1 and H3.3 are most dynamic within “large” EHV1 RCs and cells with diHuse H3. EDerm cells were transfected with plasmids encoding GFP fused to H3.1 or H3.3. At least 40h post transfection, cells were mock-infected or infected with 10 PFU/cell of abortogenic or neurotropic EHV1. Nuclear mobilities of GFP-H3.1 or –H3.3 were evaluated by FRAP between 5 and 6hpi. FRAP data for EHV1 infected cells were pooled for each histone and segregated by the absence (Small) or presence (Large) of clearly identifiable RCs. (**A**) Bar graphs present the average normalized levels of free GFP-H3.1 or – H3.3 expressed relative to the average normalized levels in mock-infected cell chromatin (set at 1). (**B**) Frequency distribution plots show the percentage of free GFP-H3.1 or –H3.3 per individual cell. Vertical solid or dashed black lines, 1 SD above the average level of free GFP-H3.1 or –H3.3 in mock-infected cell-or nucleolar-chromatin, respectively. The level of free GFP-H3.1 or –H3.3 per individual cell in the mock-infected cell-or nucleolar-chromatin are plotted in both graphs for comparison. (**C**) Dot plots present the level of GFP-H3.1 or –H3.3 in the free pool per individual cell plotted against its normalized total nuclear fluorescence intensity prior to photobleaching. (**D**) Bar graphs represent the average initial normalized fluorescence recovery rate for GFP-H3.1 or –H3.3 expressed relative to the average initial normalized fluorescence recovery rate in mock-infected cell chromatin (set at 1). (**E**) Frequency distribution plots present the initial normalized fluorescence recovery rate for GFP-H3.1 or –H3.3 per individual cell. Solid or dashed vertical black lines, 1 SD above the average initial normalized fluorescence recovery rate for GFP-H3.1 or –H3.3 in mock-infected cell-or nucleolar-chromatin, respectively. The initial normalized fluorescence recovery rate for GFP-H3.1 or –H3.3 per individual cell in the mock-infected cell-or nucleolar-chromatin are plotted in both grapns for comparison. Error bars, SEM. H3.1 n≥46 cells per treatment from 5 independent experiments; H3.3 n≥60 cells per treatment from 6 independent experiments. Different letters denote *P*<0.01; matching symbols denote *P*<0.05. Statistical significance evaluated by ANOVA with post-hoc Tukey Kramer pair-wise analysis.

Although H3.1 and H3.3 were mobilized within “large” RCs, neither histone was substantially mobilized within “small” ones. H3.1 or H3.3 free pools only tended to increase within “small” RCs (to 117 ± 4% or 119 ± 7%, respectively; *P*=ns; Fig 7A, B). Infection also did not substantially mobilize H3.1 or H3.3 within the infected-cell chromatin. Only the H3.3 free pool increased, to 129 ± 5%, within the cell chromatin surrounding “large” RCs (*P*<0.05; Fig 7A, B, Table 1).

It was not possible to discern nuclear domains likely to be enriched for viral-or cell-chromatin within cells with diffuse H3.1 or H3.3. Therefore, H3.1 or H3.3 mobility within these cells represents their net mobility due to cell– and EHV1-chromatin dynamics. H3.1 or H3.3 were nonetheless mobilized within these cells. Mobilization increased H3.1 or H3.3 free pools to 256 ± 12% or 270 ± 9%, respectively, which were significantly larger than their free pools within “large” RCs (*P*<0.01; Fig 7A, B, Table 1). This extreme increase to H3.1 or H3.3 free pools occurred throughout this cell population and did not correlate with overall GFP-histone expression levels (Fig 7B, C). Thus, H3.1 and H3.3 were most mobilized within infected cells with diffuse H3.

Mobilization of H3.1 and H3.3 within “large” RCs and cells with diffuse H3 altered their chromatin residency to cause a net increase in their free pools. In cells with diffuse H3.1 or H3.3 this was accompanied by increased fast chromatin exchange (373 ± 36% or 502 ± 40%, respectively, *P*<0.01; Fig 7D, E, Table 2). In contrast, only H3.3 fast chromatin exchange increased (to 260 ± 40%; *P*<0.05), while that of H3.1 was not significantly altered (155 ± 18%), within “large” RCs (Fig 7D, E, Table 2). However, analysis of the H3.3 fast chromatin exchange rate per individual cell revealed that H3.3 fast chromatin exchange only increased in “large” RCs within a minor subpopulation of cells (23% of cells had a rate >1SD above the mock-infected cell chromatin average; Fig 7E, Table 2). Thus, most cells did not have increased H3.3 fast chromatin exchange within “large” RCs. These data show that H3.1 and H3.3 are differentially mobilized in “large” RCs or cells with diffuse H3 and suggest that different mechanisms contribute to increase H3.1 or H3.3 free pools within either.

In summary, EHV1 did not particularly mobilize H3.1 or H3.3 within “small” RCs or domains enriched for infected-cell chromatin. Only H3.3 was partially mobilized within the cell chromatin surrounding “large” RCs. EHV1 did, however, mobilize canonical H3.1 and variant H3.3 histones within “large” RCs and cells with diffuse H3. This mobilization caused a net increase in H3.1 and H3.3 free pools. However, mobilization differentially affected H3.1 and H3.3 fast chromatin exchange. Whereas the fast chromatin exchange of H3.1 and H3.3 increased within cells with diffuse H3, only H3.3 fast chromatin exchange increased within “large” RCs and only in a minor subpopulation of cells. Thus, the free pools of H3.1 and H3.3 predominantly increased within “large” RCs without substantial alteration to their fast chromatin exchange.

### H3.3 is mobilized to a greater degree than H3.1 in EHV1 “large” RCs

H3.1 and H3.3 were mobilized within “large” RCs and in cells with diffuse H3. In these cells, H3.3 appeared mobilized to a greater degree than H3.1, at least relative to their respective mobilities within mock-infected cell chromatin (Fig 7A, D). H3.3 may thus be more susceptible to histone mobilizing factors or, alternatively, H3.1 may have higher intrinsic mobility and therefore lower potential magnitude for mobilization. To investigate these possibilities, we evaluated mobilization of H3.3 relative to H3.1 (S2 Fig, S2 Table). Surprisingly, H3.3 was less mobile than H3.1 in mock-infected cell chromatin. H3.3 had a relatively smaller free pool and slower fast chromatin exchange than H3.1 (85 ± 5% or 60 ± 10%, respectively, *P*<0.05; S2 Fig; S2 Table). Within nucleoli, however, H3.1 and H3.3 free pools were similar, despite significantly slower H3.3 fast chromatin exchange (56 ± 7% that of H3.1, *P*<0.01; S2 Fig, S2 Table). These data indicate that EDerm cells have a dynamic population of H3.1 that is intrinsically more mobile than H3.3.

Infection mobilized H3.1 and H3.3 such that H3.3 free pools increased significantly relative to those of H3.1 within “large” RCs or cells with diffuse H3 (113 ± 3%, *P*<0.01 or 112 ± 3%, *P*<0.05, respectively; S2 Fig, S2 Table). H3.3 average fast chromatin exchange also increased by a greater magnitude than H3.1 in “large” RCs (260 ± 40% vs 155 ± 18%) and cells with diffuse H3 (502 ± 40% vs 373 ± 26%; Fig 7D, E). However, this resulted in relatively similar H3.3 and H3.1 fast chromatin exchange rates (S2 Fig, S2 Table).

Together these data show that a population of H3.1 is inherently more mobile than H3.3 within EDerm cell chromatin, with a larger unbound population and faster low-affinity chromatin exchange. In “large” RCs H3.3 was mobilized by a greater magnitude, and in diffuse cells H3.3 fast chromatin exchange increased by a greater magnitude. This mobilization resulted in relatively larger free pools of H3.3 than H3.1, despite relatively similar fast chromatin exchange. H3.3 may thus be more susceptible to histone mobilizing factors within “large” RCs or cells with diffuse H3. Furthermore, the relatively larger H3.3 free pool suggests that mobilizing factors may preferentially evict H3.3 from, or prevent it from binding in, EHV1 chromatin.

### EHV1 mobilizes canonical H2A and variant H2A.Z, H2A.X, and macroH2A

Canonical H3.1 and variant H3.3 were differentially mobilized during EHV1 infection. To further interrogate whether EHV1 preferentially mobilizes particular histone types we next evaluated mobility of canonical H2A and variant H2A.Z, H2A.X, and macroH2A during EHV1 infection (the uniquely mobile H2A.B variant is presented and discussed below). GFP-H2A, –H2A.Z, –H2A.X, and –macroH2A had discrete granular nuclear localization consistent with their assembly in chromatin (Fig 8). Regions of varying fluorescence intensity marked chromatin regions relatively enriched or depleted for each histone (Fig 8). The evaluated H2A variants and canonical H2A were typically largely depleted from nucleoli, although H2A and H2A.Z often had larger nucleolar pools (less depletion) than H2A.X or macroH2A (Fig 8). Consistent with other evaluated core histones, H2A, H2A.Z, H2A.X and macroH2A were also largely depleted from EHV1 RCs (Fig 8).

**Fig 8.**
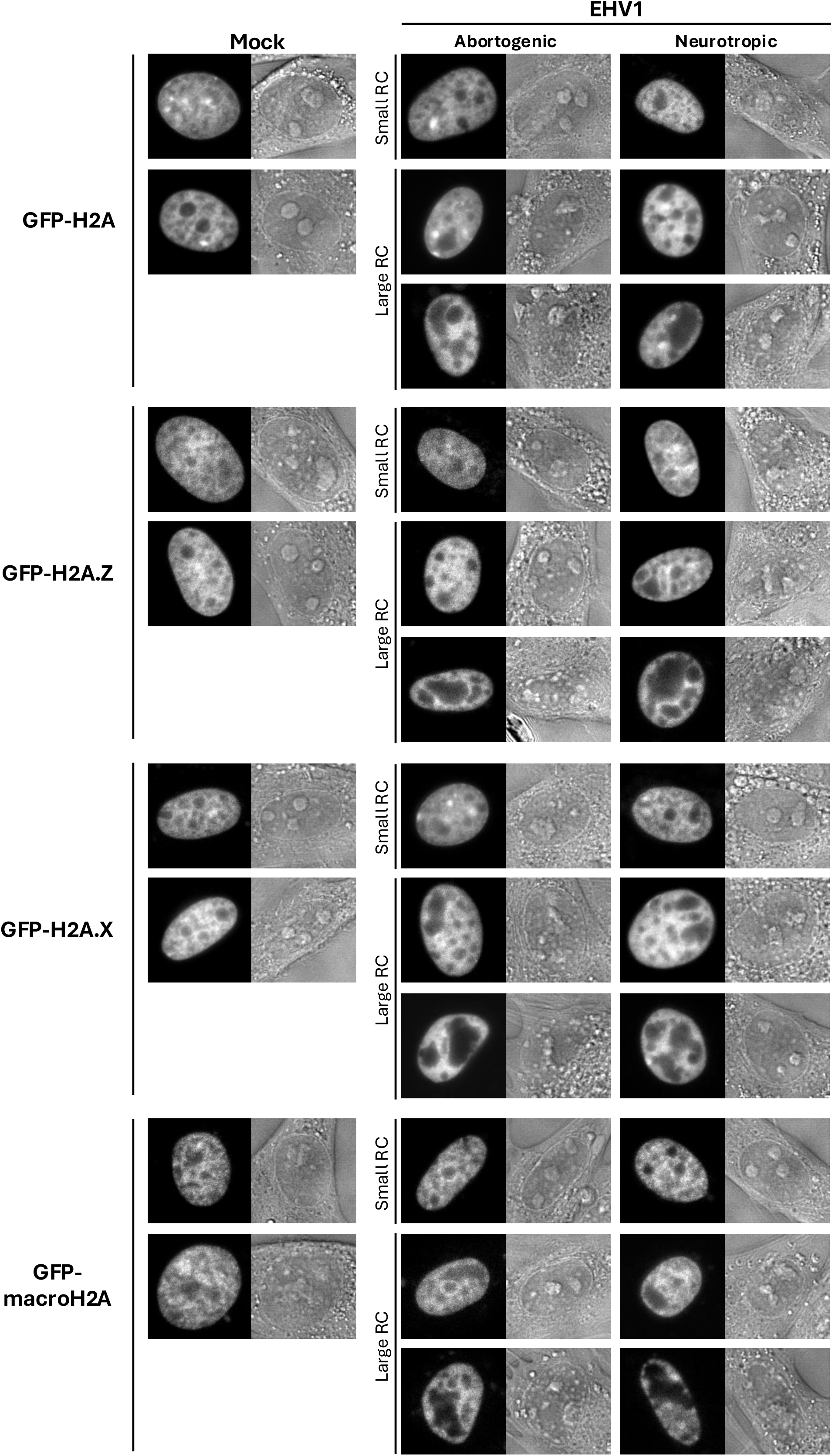
Canonical H2A and variant H2A.Z, H2A.X, and macroH2A are largely depleted from EHV1 RCs. Digital fluorescent (left panels) and DIC (right panels) micrographs show the nucleus of EDerm cells expressing GFP fused to H2A, H2A.Z, H2A.X, or macroH2A. At least 40h after transfection with plasmids encoding the GFP-histone fusion proteins, cells were mock-infected or infected with 10 PFU/cell of abortogenic or neurotropic EHV1. Live cells were imaged between 5 and 6hpi.

Histone H2A variants are the most numerous and functionally diverse to regulate nucleosome stability and DNA accessibility. FRAP analysis revealed that, as expected, canonical and variant GFP-H2A fusion proteins had distinct mobilities within the cell chromatin that corresponded to their effects on nucleosome stability (S3 Fig)(24, 58). GFP-H2A.Z was the most mobile variant with a significantly larger free pool and increased fast chromatin exchange relative to canonical H2A (116 ± 3% and 144 ± 12%, respectively, *P*<0.01; S3 Fig, S2 Table). Conversely, GFP-macroH2A was the least mobile variant. Its free pool or fast chromatin exchange, however, were not significantly different from those of canonical H2A (91 ± 3% or 95 ± 10%, respectively; S3 Fig, S2 Table). H2A, H2A.Z, H2A.X, and macroH2A were all more mobile within nucleoli and, surprisingly, were similarly mobile within this domain (Fig 9, S3 Fig, S2 Table). Only the macroH2A fast chromatin exchange rate was relatively slower than that of H2A within nucleoli (76 ± 7%, *P*<0.05; S3 Fig, S2 Table).

**Fig 9.**
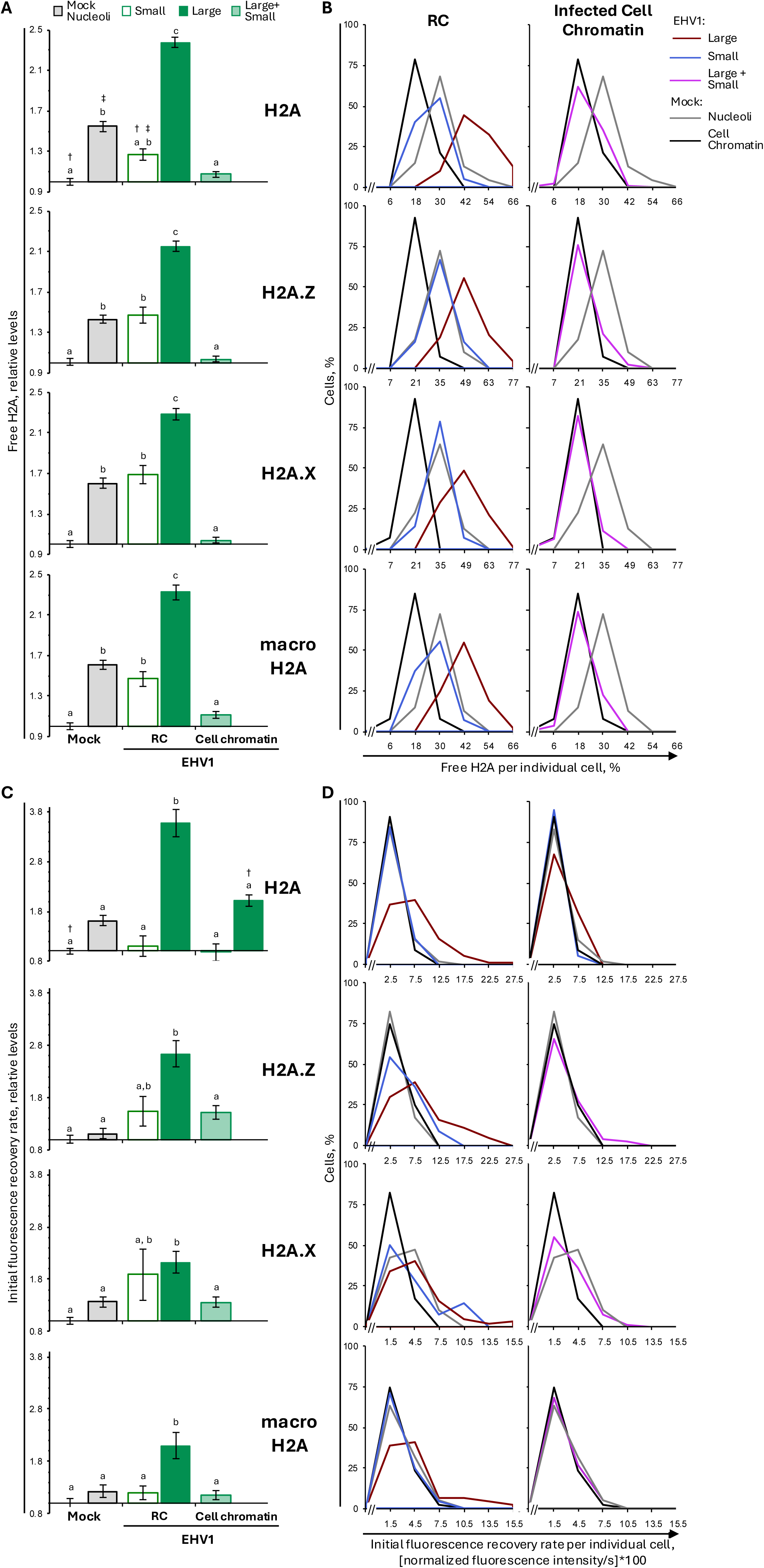
Canonical H2A and variant H2A.Z, H2A.X, and macroH2A are most mobile within “large” EHV1 RCs. EDerm cells were transfected with plasmids expressing GFP-H2A, –H2A.Z, –H2A.X, or –macroH2A at least 40h prior to mock-infection or infection with 10 PFU/cell of abortogenic or neurotropic EHV1. Nuclear mobilities of GFP-H2A, –H2A.Z, –H2A.X or –macroH2A were evaluated by FRAP between 5 and 6hpi. FRAP data for EHV1 infected cells were pooled for each histone and segregated by the absence (Small) or presence (Large) of clearly identifiable RCs. (**A**) Bar graphs represent the average normalized levels of free GFP-H2A, –H2A.Z, –H2A.X or –macroH2A expressed relative to their average normalized levels in mock-infected cell chromatin (set at 1). (**B**) Frequency distribution plots show the percentage of free GFP-H2A, –H2A.Z, –H2A.X or –macroH2A per individual cell. Solid or dashed vertical black lines, 1 SD above the average level of free GFP-H2A, –H2A.Z, –H2A.X or –macroH2A in mock-infected cell-or nucleolar-chromatin, respectively. The level of free GFP-H2A, –H2A.Z, –H2A.X or –macroH2A per individual cell in the mock-infected cell-or nucleolar-chromatin are plotted in both graphs for comparison. (**C**) Bar graphs present the average initial normalized fluorescence recovery rate for GFP-H2A, –H2A.Z, –H2A.X or –macroH2A expressed relative to their average initial normalized fluorescence recovery rate in mock-infected cell chromatin (set at 1). (**D**) Frequency distribution plots present the initial normalized fluorescence recovery rate for GFP-H2A, – H2A.Z, –H2A.X or –macroH2A per individual cell. Solid or dashed vertical black lines, 1 SD above their average initial normalized fluorescence recovery rate in mock-infected cell-or nucleolar-chromatin, respectively. The initial normalized fluorescence recovery rate for GFP-H2A, –H2A.Z, –H2A.X or –macroH2A per individual cell in the mock-infected cell-or nucleolar-chromatin are plotted in both graphs for comparison. Error bars, SEM. H2A n≥47 cells per treatment from 5 independent experiments, H2A.Z, H2A.X, macroH2A n≥40 cells per treatment from 4 independent experiments. Different letters denote *P*<0.01; matching symbols denote *P*<0.05. Statistical significance evaluated by ANOVA with post-hoc Tukey Kramer pair-wise analysis.

Infection mobilized H2A, H2A.Z, H2A.X, and macroH2A within EHV1 RCs (Fig 9, S3 Fig). In “small” RCs, H2A, H2A.Z, H2A.X, or macroH2A free pools increased to 127 ± 6%, 147 ± 8%, 169 ± 9%, or 147 ± 8%, respectively (*P*<0.01, except H2A *P*<0.05; Fig 9A, Table 1). This equated to an extreme increase in free pool levels within the “small” RCs of more than 50% of cells. H2A, H2A.Z, H2A.X, and macroH2A were further mobilized within “large” RCs, where their free pools increased to 238 ± 5%, 215 ± 5%, 229 ± 6%, or 233 ± 6%, respectively (*P*<0.01; Fig9 A, Table 1). This corresponded to an extreme increase in free pools within “large” RCs throughout the entire infected cell population (Fig 9A, B, Table 1). Moreover, greater than 70% of cells had canonical or variant H2A free pools within “large” RCs greater than 1SD above their average levels within mock-infected nucleoli. Consistently, H2A, H2A.Z, H2A.X, and macroH2A free pools within “large” RCs were significantly greater than those within mock-infected nucleoli. These data indicate that H2A, H2A.Z, H2A.X, and macroH2A association in EHV1 chromatin is more dynamic or unstable than their association in nucleolar chromatin, further in support of the dynamic instability of EHV1 lytic chromatin.

EHV1 mobilization of H2A, H2A.Z, H2A.X, and macroH2A within “small” or “large” RCs resulted in a net increase in their free pools within RCs throughout the infected cell population. In “small” RCs, canonical or variant H2A free pools increased without substantially affecting their fast chromatin exchange (Fig 9C, D, Table 2). Only H2A.X fast chromatin exchange increased within “small” RCs in a minor subpopulation of cells, shown by the bimodal frequency distribution of the H2A.X initial fluorescence recovery rate per individual cell (Fig 9D, Table 2). Mobilization of canonical or variant H2A within “large” RCs, however, did increase their fast chromatin exchange. H2A, H2A.Z, H2A.X, or macroH2A fast chromatin exchange increased to 357 ± 27%, 263 ± 25%, 212 ± 14%, or 210 ± 26%, respectively, within “large” RCs (*P*<0.01; Fig 9C, D, Table 2). Thus, mobilization of H2A, H2A.Z, H2A.X, and macroH2A within “large” RCs resulted from changes in their chromatin residency and low-affinity chromatin exchange, whereas mobilization in “small” RCs primarily affected their chromatin residency.

While variant and canonical H2A were mobilized within domains enriched for EHV1 chromatin, they were not significantly mobilized within domains of infected-cell chromatin. Free H2A, H2A.Z, H2A.X, or macroH2A pools were similar in EHV1– and mock-infected cell chromatin (Fig 9A, B, Table 1). Moreover, with exception of canonical H2A, their fast chromatin exchange rates were also largely unaltered. H2A fast chromatin exchange increased to 202 ± 12% in the cell chromatin surrounding “large” RCs (*P*<0.01; Fig 9C, D, Table 2).

In summary, EHV1 mobilized H2A, H2A.Z, H2A.X, and macroH2A. These histones were primarily mobilized within nuclear domains enriched for EHV1 chromatin (RCs).

Mobilization increased their free pools within “small” and “large” RCs but only increased their fast chromatin exchange within “large” ones. These results are consistent with different mechanisms to increase free pools within “small” or “large” RCs. In contrast, only canonical H2A was mobilized within infected-cell chromatin, and only within cells containing “large” RCs. In this case, mobilization increased its fast chromatin exchange without altering its free pool.

### Variant and canonical H2A histones are diHerentially mobilized by EHV1

H2A, H2A.Z, H2A.X, and macroH2A were mobilized within EHV1 RCs relative to their intrinsic mobility within mock-infected cell chromatin. As it was possible that particular H2A types were differentially mobilized by infection, we examined variant H2A mobility relative to that of canonical H2A. In EHV1 “small” RCs, the inherent relationships between variant and canonical H2A mobilities were not well conserved. H2A.X was mobilized to the greatest degree such that its mobility within “small” RCs resembled more that of H2A.Z than H2A (Fig 9, S3 Fig, S2 Table). The H2A.X free pool increased to 133 ± 7% (*P*<0.01) and its fast chromatin exchange rate tended to increase (to 136 ± 29%; *P*=ns) relative to H2A (S3 Fig, S2 Table). H2A.Z was also further mobilized relative to H2A within “small” RCs. H2A.Z fast chromatin exchange increased from 131 ± 9% that of H2A within the mock-infected cell chromatin to 172 ± 26% that of H2A within “small” RCs (S3 Fig, S2 Table). Despite increased fast chromatin exchange, H2A.Z free pools remained relatively similar to those of H2A within “small” RCs or mock-infected cell chromatin (131 ± 8% or 127 ± 3% that of H2A, respectively; S3 Fig, S2 Table). Together, these data suggest that H2A.X and H2A.Z may be preferentially mobilized within “small” RCs or more susceptible to histone mobilizing factors in abundance at earlier stages of infection.

Further mobilization in “large” RCs resulted in relatively similar free pools of H2A, H2A.X, and H2A.Z (S3 Fig, S2 Table). However, H2A fast chromatin exchange increased by a greater magnitude than that of H2A.Z or H2A.X. Thus, H2A.Z fast chromatin exchange was similar to (117 ± 9%; *P*=ns), while that of H2A.X was significantly slower than (66 ± 6%; *P*<0.01), that of H2A within “large” RCs (S3 Fig, S2 Table). Although macroH2A was also further mobilized within “large” RCs, it remained the least mobile variant within them. It’s free pool or fast chromatin exchange rate were 89 ± 2% or 64 ± 8% those of H2A, respectively (*P*<0.01; S3 Fig, S2 Table). These data suggest that canonical H2A may be more susceptible to histone mobilizing factors in abundance within “large” RCs.

In contrast to mobilization of canonical and variant H2A within RCs, infection had little effect on the intrinsic relationships between variant and canonical H2A mobilities within the surrounding infected-cell chromatin. However, as in “large” RCs, H2A.X and macroH2A fast chromatin exchange further decreased relative to H2A (102 ± 7% to 84 ± 5% or 95 ± 10% to 69 ± 5%, *P*<0.05 or <0.01, respectively), consistent with H2A mobilization within the cell chromatin surrounding “large” RCs (S3 Fig, S2 Table). Regardless, the H2A, H2A.Z, H2A.X, and macroH2A free pools were proportionately maintained within infected-cell chromatin.

In summary, infection differentially mobilized canonical H2A and variant H2A.Z, H2A.X, and macroH2A within domains enriched for EHV1 chromatin. H2A.X and H2A.Z were initially mobilized by a larger degree relative to H2A within “small” RCs, suggesting that they may be preferentially mobilized or more susceptible to mobilization factors in abundance at earlier stages of infection. Mobilization of H2A.X and H2A.Z in “small” RCs, however, were distinct in that the H2A.X free pool was most enhanced whereas H2A.Z fast chromatin exchange was most enhanced. The differential mobilization of H2A.X and H2A.Z within “small” RCs indicates that they are mobilized by distinct mechanisms or have differential associations within EHV1 chromatin. At later stages of infection, the further mobilization of H2A, H2A.X, and H2A.Z resulted in relatively similar free pools within “large” RCs. However, despite similar free pools, H2A.X fast chromatin exchange was relatively slower than that of H2A or H2A.Z. Moreover, despite further mobilization of macroH2A within “large” RCs its free pool and fast chromatin exchange remained relatively smaller and slower than those of H2A. The differential relative mobilities of H2A, H2A.Z, H2A.X, and macroH2A within “large” RCs suggests that they have distinct EHV1 chromatin interactions.

### Variant H2A.B is uniquely mobilized during EHV1 infection

H2A.B is the most unique and dynamic H2A variant. It had distinct nuclear localization patterns that ranged from largely homogenous throughout the nucleus to pronounced enrichment within nucleoli (Fig 10). The variable localization of equine H2A.B resembled the cell cycle-related localization of human H2A.B and likewise may relate to cell cycle stage (69). FRAP analysis revealed that H2A.B localization patterns corresponded with distinct mobilities within non-nucleolar chromatin. We arbitrarily numbered the mobility groups from the slowest (1), which corresponded to relatively homogenously distributed H2A.B, to the fastest (4), which corresponded to pronounced nucleolar enrichment (Fig 10). Most cells were within mobility groups 3 (38%) and 2 (31%), with smaller populations within the most extreme mobility groups 1 (13%) and 4 (18%) (Fig 10, S3 Table). The slowest mobility group 1 had the smallest H2A.B free pool and slowest fast chromatin exchange, whereas the fastest mobility group 4 had the largest and fastest, respectively (Fig 11, Tables 1 and 2). The variable mobility of H2A.B within non-nucleolar chromatin did not relate to GFP-H2A.B expression levels measured as the total normalized nuclear fluorescence before photobleaching (S4 Fig).

**Fig 10.**
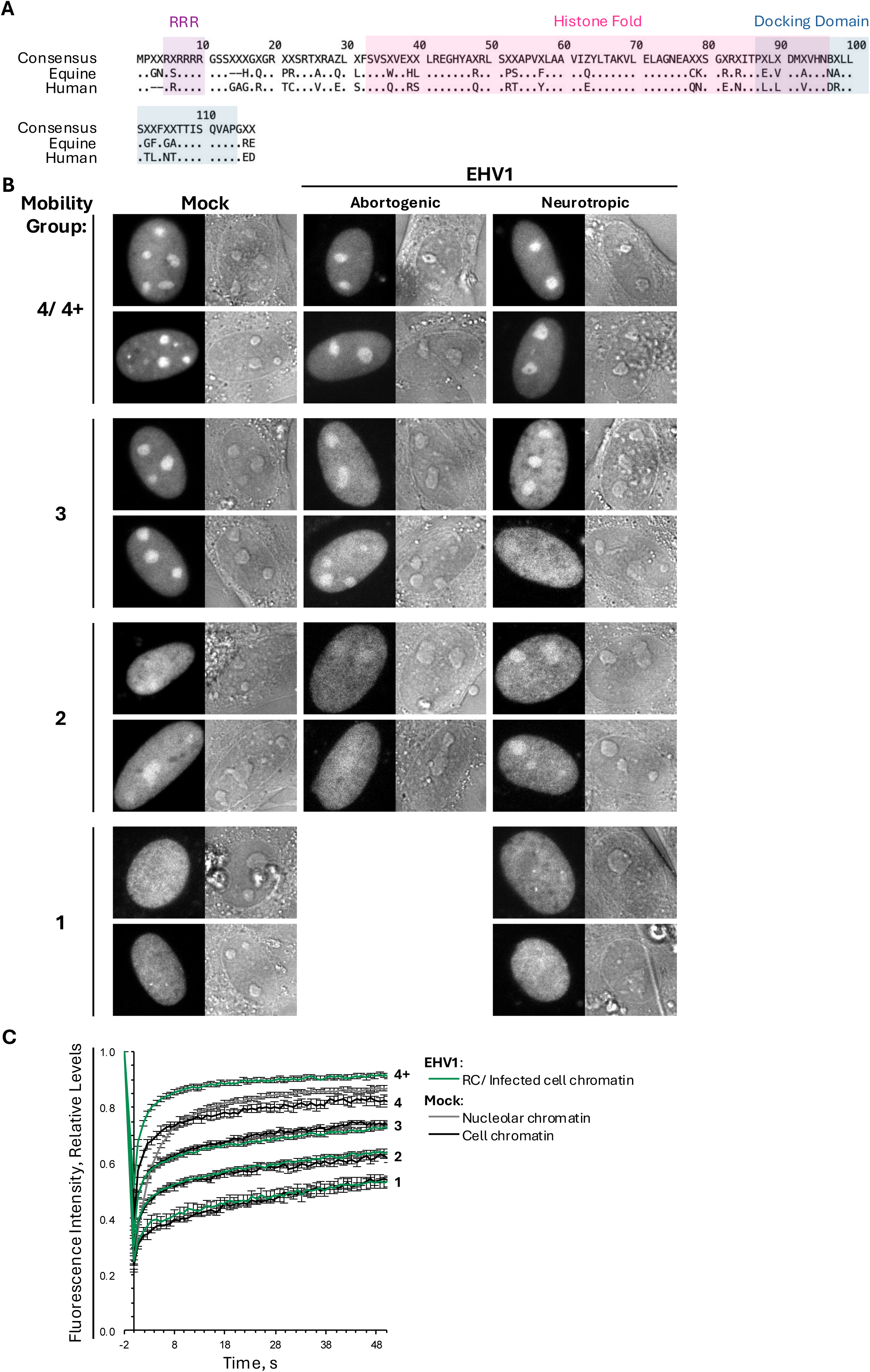
Variant H2A.B is neither depleted nor enriched in EHV1 RCs. EDerm cells were transfected with plasmids encoding GFP fused to H2A.B. At least 40h after transfection, cells were mock-infected or infected with 10 PFU/cell of abortogenic or neurotropic EHV1. Live cells were imaged and GFP-H2A.B mobility evaluated by FRAP between 5 and 6hpi. FRAP data from EHV1-infected cells or the cell chromatin of mock-infected cells were combined and segregated into mobility groups. (**A**) Digital fluorescent (left panels) and DIC (right panels) micrographs show the nucleus of cells expressing GFP-H2A.B. (**B**) Line graph presents FRAP of GFP-H2A.B in mock-or EHV1-infected EDerm cells. Error bars, SEM; n≥38 cells per treatment from 4 independent experiments.

**Fig 11.**
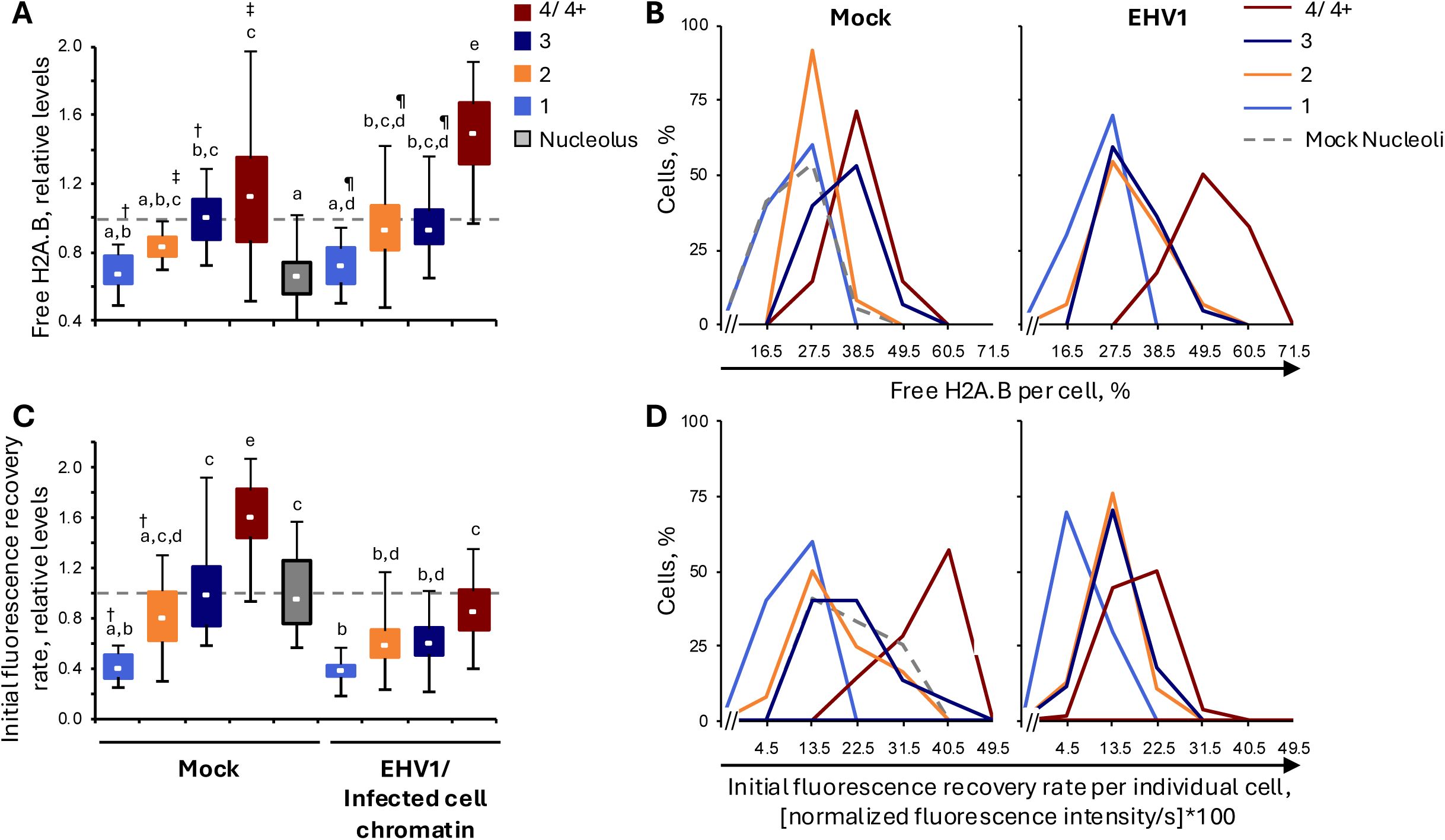
H2A.B is most mobile in a subpopulation of EHV1 infected cells. EDerm cells were transfected with plasmids encoding GFP-H2A.B at least 40h prior to mock-infection or infection with 10 PFU/cell of abortogenic or neurotropic EHV1. Nuclear mobility of GFP-H2A.B was evaluated by FRAP between 5 and 6hpi. FRAP data for EHV1-infected cells or for mock-infected cell chromatin were pooled and segregated by mobility group. (**A**) Bar graph presents the average normalized level of free GFP-H2A.B per mobility group expressed relative to the average normalized level in the mock-infected cell chromatin for mobility group 3 (set at 1). (**B**) Frequency distribution plots show the percentage of free GFP-H2A.B per individual cell segregated by mobility group. Dashed vertical black lines, 1 SD above or below the average level of free H2A.B in the mock-infected cell chromatin for mobility group 3. (**C**) Bar graph presents the average initial normalized fluorescence recovery rate per GFP-H2A.B mobility group expressed relative to the average initial normalized fluorescence recovery rate in the mock-infected cell chromatin of mobility group 3 (set at 1). (**D**) Frequency distribution plots present the initial normalized fluorescence recovery rate for GFP-H2A.B per individual cell segregated by mobility group. Solid or dashed vertical black lines, 1 SD above or below the average initial normalized fluorescence recovery rate in the mock-infected cell chromatin of mobility group 3. Error bars, SEM. n≥38 cells per treatment from 4 independent experiments. Different letters denote *P*<0.01; matching symbols denote *P*<0.05. Statistical significance evaluated by ANOVA with post-hoc Tukey Kramer pair-wise analysis.

H2A.B mobility within nucleoli was similar in all evaluated mock-infected cells (Fig 10B). This suggests that factors to regulate H2A.B chromatin exchange primarily do so within non-nucleolar chromatin and have little impact on H2A.B exchange within nucleolar chromatin. The similar mobility of H2A.B within nucleoli, regardless of its mobility within the cell chromatin, further supports the independence of mobility and GFP-H2A.B expression levels. Furthermore, these data show that H2A.B chromatin exchange within nucleoli is independent of nucleoli number or relative H2A.B enrichment within them. The H2A.B free pool within nucleoli was similar to that within the surrounding cell chromatin for mobility groups 1 and 2 (67 ± 2%, vs 67 ± 6% and 83 ± 3%, respectively, of the H2A.B free pool within mobility group 3; Fig 11, Table 1). H2A.B fast chromatin exchange within nucleoli, however, resembled more that of mobility group 3, which had a significantly larger H2A.B free pool (Fig 11, Tables 1 and 2). Thus, given the similar fast chromatin exchange of H2A.B within nucleoli and the non-nucleolar chromatin within mobility group 3, these data suggest that H2A.B chromatin residency within nucleoli is greater to account for its relatively smaller free pool.

H2A.B is the most dynamic H2A variant and its chromatin exchange kinetics resemble more linker, rather than other core histones (54, 55, 57). Consistently, within non-nucleolar cell chromatin all H2A.B mobility groups had larger free pools (with exception of mobility group 1) and increased fast chromatin exchange relative to canonical H2A (S3 Fig, S2 Table). Within nucleoli, however, the H2A.B free pool was significantly smaller (74 ± 2%) and its fast chromatin exchange significantly faster (567 ± 30%) relative to canonical H2A (*P*<0.01, S3 Fig, S2 Table). Thus, H2A.B chromatin turnover is exceptionally rapid relative to that of H2A within nucleoli, yet H2A.B is also more likely than H2A to be assembled in nucleolar chromatin at any given time.

EHV1 infection surprisingly did not alter H2A.B nuclear localization. H2A.B still had varying localization patterns that ranged from largely homogenous throughout the nucleus to pronounced nucleolar enrichment (Fig 10). Significant H2A.B depletion from, or enrichment in, EHV1 RCs was not observed. Nuclear domains likely to be enriched in viral-or cell-chromatin were therefore indistinguishable and measured H2A.B mobility represents its net mobility due exchange with cell– and EHV1-chromatin. H2A.B also consistently segregated into distinct mobility groups within infected cells (Fig 10B, S5 Fig). The slowest H2A.B mobility groups within infected cells were similar to the slowest mobility groups within mock-infected cell chromatin (1-3; Fig 10B, S5 Fig). Only the fastest mobility group within EHV1-infected cells, designated as 4+, was mobilized relative to mock-infected cells. This most mobile H2A.B group was evident regardless of whether cells were infected with abortogenic or neurotropic EHV1, although the relative populations of cells within this, or the slower mobility groups 1-3, varied (S5 Fig, S3 Table). Most cells infected with abortogenic EHV1 segregated into mobility group 4+, with no cells within the slowest mobility group 1 (S5 Fig, S3 Table). Most cells infected with neurotropic EHV1, meanwhile, segregated into mobility groups 2 and 3. These data suggest that H2A.B mobilization may relate to infection progression as the abortive EHV1 strain tended to have slightly faster replication kinetics than the neurotropic one in EDerm cells. Alternatively, other factors, such as specific virus-host protein-protein interactions, may contribute to regulate H2A.B chromatin exchange within EHV1-infected cells.

H2A.B mobility group 4+ was the most mobile. Fluorescence within these cells recovered faster than within any other mobility group of mock-or EHV1-infected cells or within mock-infected nucleoli (Fig 10B). H2A.B mobility group 4+ had the largest free pool (149 ± 3%) and the greatest fast chromatin exchange (86 ± 3%) within EHV1-infected cells (expressed relative to H2A.B mobility group 3 in mock-infected cell chromatin; Fig 11, Tables 1 and 2). The H2A.B free pool within mobility group 4+ was significantly increased relative to that within mock-infected mobility group 4 (149 ± 3% vs 113 ± 4%), although its fast chromatin exchange rate was significantly slower (86 ± 3% vs 161 ± 15%, *P*<0.01; Fig 11, Tables 1 and 2). Thus H2A.B mobilization within this subpopulation of infected cells (4+) increased its net level in the free pool despite relatively slower fast chromatin exchange. These data suggest that within this population of infected cells H2A.B is less likely to re-bind chromatin once unbound but has slower turnover when bound.

The inability to distinguish EHV1 RCs in H2A.B expressing cells precluded comparative analysis of H2A.B and H2A mobilization within domains enriched for viral or infected-cell chromatin. As a surrogate analysis, H2A.B mobility was expressed relative to H2A mobility within infected-cell chromatin or “large” RCs. The intrinsic property of H2A.B as the most dynamic H2A variant was maintained relative to H2A within infected-cell chromatin (S3 Fig, S2 Table). All H2A.B mobility groups had larger free pools (with exception of mobility group 1) and enhanced fast chromatin exchange relative to H2A within the infected-cell chromatin (S3 Fig, S2 Table). With respect to H2A mobility within “large” RCs, however, only H2A.B mobility group 4+ had a larger free pool (108 ± 2%, *P*<0.05) and enhanced fast chromatin exchange (257 ± 8%, *P*<0.01) relative to H2A (S3 Fig, S2 Table). Mobility groups 1 to 3 had significantly smaller free pools and, with exception of mobility group 1, significantly increased fast chromatin exchange relative to H2A within “large” RCs (S3 Fig, S2 Table). The relative mobilities of H2A.B mobility groups 1 to 3 and H2A within “large” RCs were similar to the relative mobilities of H2A.B and H2A within mock-infected nucleolar chromatin.

These data highlight the unique chromatin exchange of H2A.B. EHV1 did not notably alter H2A.B nuclear localization and only mobilized H2A.B within a subpopulation of infected cells (4+). Within this subpopulation, H2A.B free pools increased despite a relative decrease in fast chromatin exchange. Thus, in these cells H2A.B is less likely to bind in chromatin but undergoes slower low-affinity turnover when bound.

### Linker histone H1.2 is most mobile within EHV1 RCs

Linker histones are the most dynamic histone and undergo rapid chromatin exchange (48, 53–56). We evaluated H1.2 mobility during EHV1 infection as a representative linker histone given that it is abundantly expressed throughout the cell cycle, assembles in euchromatin, and is the H1 variant most mobilized by HSV1 (48, 56, 70). GFP-H1.2 had discrete granular distribution with relative nucleolar depletion. However, depletion from nucleoli tended to be less pronounced than that of most core histones (Fig 12A). FRAP analysis revealed that, as expected, H1.2 was more mobile than core histones and also more mobile within nucleoli than within the surrounding cell chromatin (Fig 12B).

**Fig 12.**
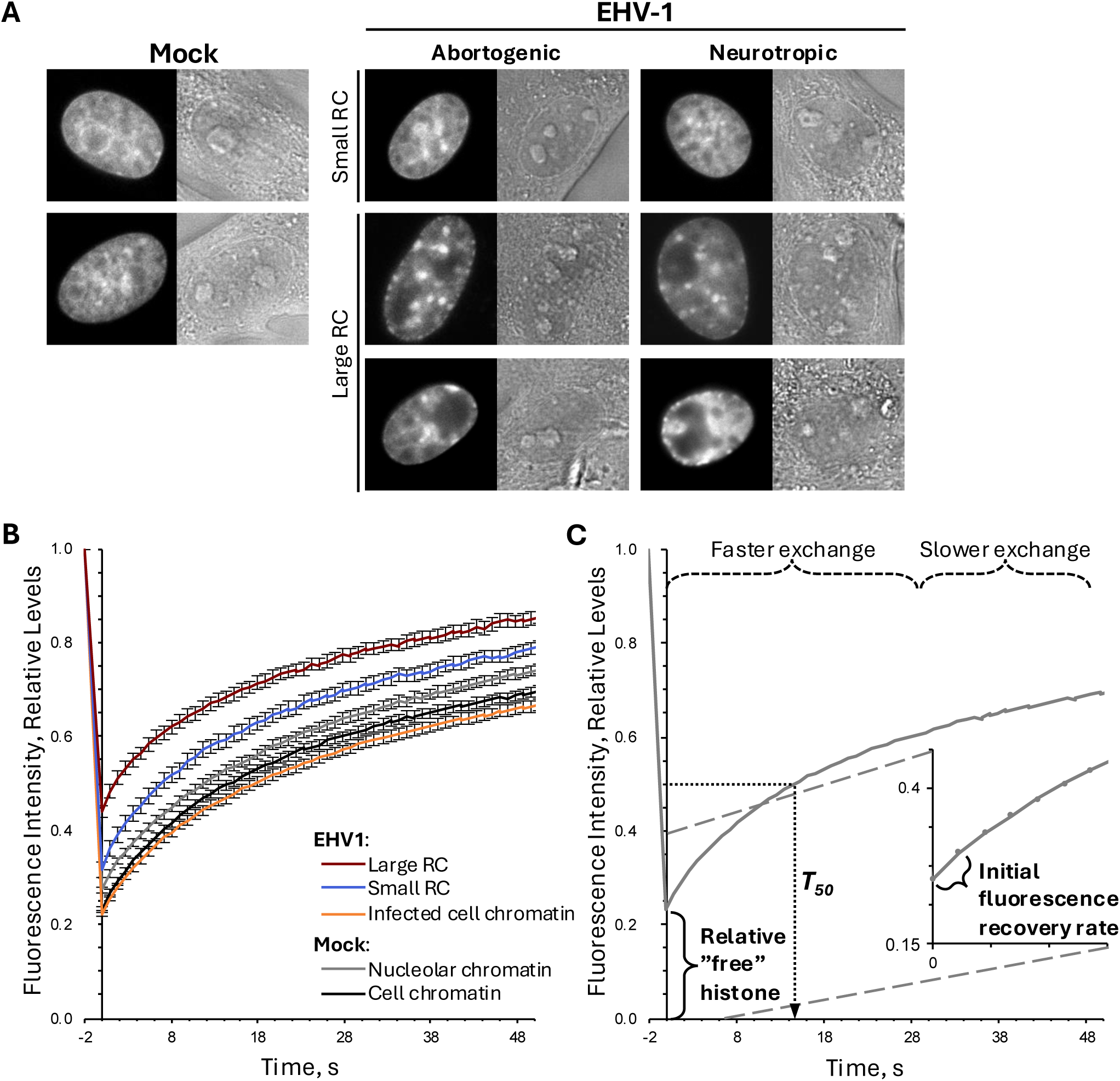
EHV1 mobilizes linker histone H1.2. EDerm cells were transfected with plasmids encoding GFP fused to H1.2 at least 40h before mock-infection or infection with 10 PFU/cell of abortogenic or neurotropic EHV1. Live cells were imaged and GFP-H1.2 mobility evaluated by FRAP between 5 and 6hpi. FRAP data from EHV1 infected cells were combined and segregated by the presence (RC large) or absence (RC small) of clearly identifiable RCs. (**A**) Digital fluorescent (left panels) and DIC (right panels) micrographs show the nucleus of cells expressing GFP-H1.2. (**B**) Line graph presents FRAP of GFP-H1.2 in mock-or EHV1-infected cells. (**C**) Line graph of a representative GFP-H1.2 FRAP in mock-infected cell chromatin (an area such as that denoted by the solid circle in Figure 2A, mock). The surrogate measures for the levels of H1.2 not bound in chromatin (available in the “free” pools) or weakly bound in chromatin (undergoing fast chromatin exchange) are calculated as for the core histones (described in Figure 2). The *T_50_*, a summary measure that primarily reflects low-affinity chromatin interactions, is calculated as the time after photobleaching to recover 50% normalized fluorescence in the photobleached region. Error bars, SEM; n≥38 cells per treatment from 4 independent experiments.

Infection altered H1.2 localization such that it was relatively depleted from viral RCs and accumulated in discrete puncta within domains enriched for infected-cell chromatin (Fig 12A). The redistribution of H1.2 within infected cells was reflected in its mobility. H1.2 was mobilized within EHV1 RCs (Fig 12B). This mobilization apparently related to infection progression as H1.2 was further mobilized in “large” relative to “small” RCs (Fig 12B). Conversely, H1.2 mobility decreased in the infected-cell chromatin such that fluorescence recovered even slower within it than within mock-infected cell chromatin (Fig 12B). The apparent stabilization of H1.2 binding within infected-cell chromatin is consistent with its accumulation (or immobilization) in discrete puncta.

As an overall measure of H1.2 mobility we calculated its *T_50_*, the time to recover 50% of the normalized fluorescence intensity within photobleached regions (Fig 12C). *T_50_* and mobility inversely relate so that a smaller *T_50_* reflects increased mobility, while a larger *T_50_* reflects reduced mobility. The *T_50_* is most influenced by low-affinity chromatin binding and consequently primarily reflects mobility of the most dynamic histone populations. H1.2 mobilization in “small” or “large” RCs decreased its *T_50_* to 52 ± 6% or 26 ±3 %, respectively, of that in mock-infected cell chromatin (*P*<0.01; Fig 13A, B). H1.2 was mobilized within RCs throughout the infected cell population with an extreme degree of mobilization in 61% or 93% of cells within “small” or “large” RCs, respectively (Fig 13B). H1.2 mobilization within RCs caused a net increase in its free pools, to 137 ± 6% or 191 ± 6%, and increased its fast chromatin exchange, to 142 ± 9% or 138 ± 11%, within “small” or “large” RCs, respectively (*P*<0.01; Fig 13C-F, Tables 1 and 2). Interestingly, H1.2 fast chromatin exchange in “small” or “large” RCs were similar despite a significantly larger H1.2 free pool within the “large” ones (Fig 13C, E). Thus, as infection progresses H1.2 may be less likely to assemble in, or be displaced from, EHV1 chromatin to account for its increased free pool within “large” RCs.

**Fig 13.**
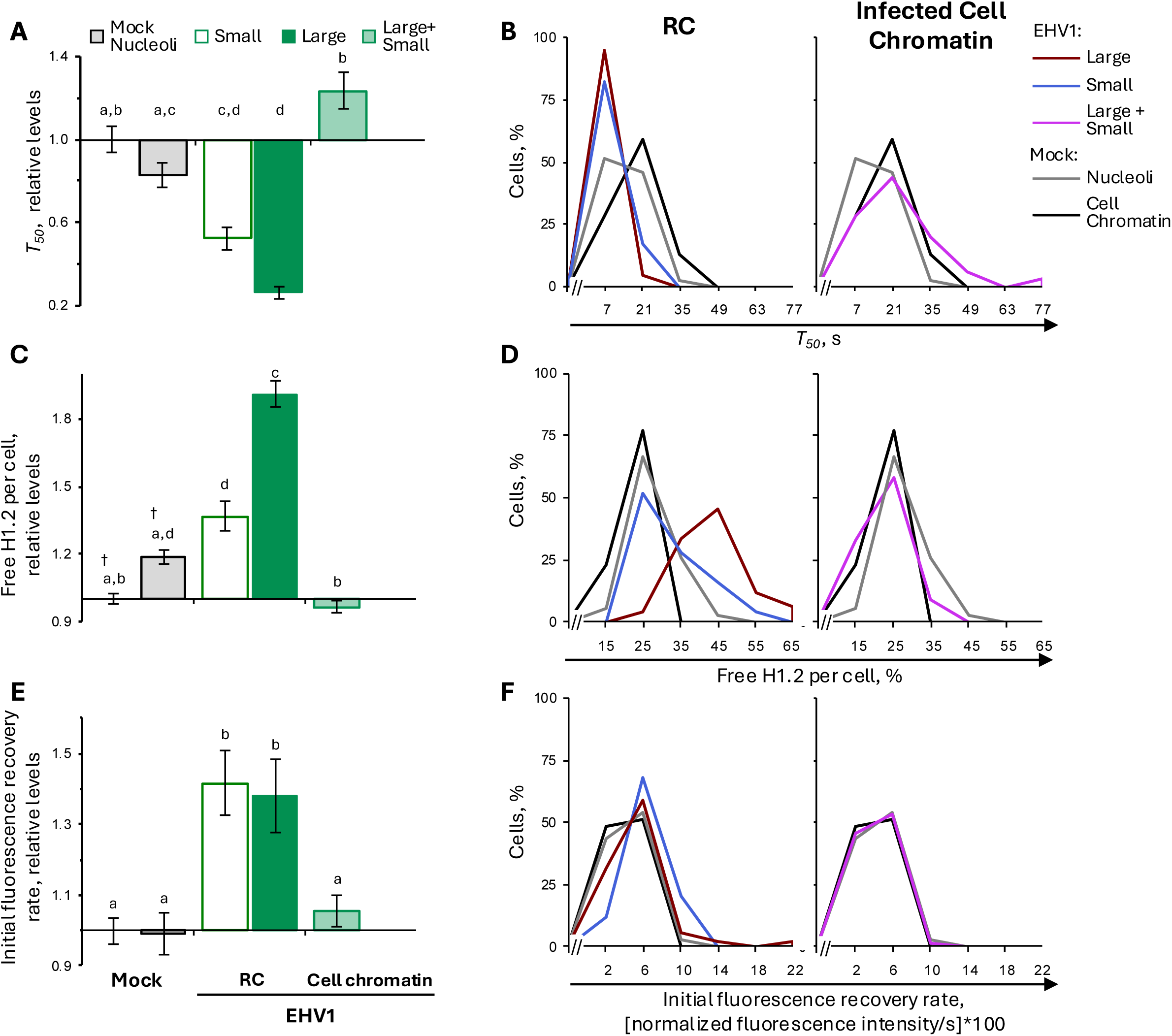
H1.2 is most dynamic in EHV1 RCs. EDerm cells were transfected with plasmids encoding GFP-H1.2 at least 40h prior to mock-infection or infection with 10 PFU/cell of abortogenic or neurotropic EHV1. Nuclear mobility of GFP-H1.2 was evaluated by FRAP between 5 and 6hpi. FRAP data for EHV1 infected cells were pooled and segregated by the absence (Small) or presence (Large) of clearly identifiable RCs. (**A**) Bar graph presents the average *T_50_* for GFP-H1.2 expressed relative to the average *T_50_* in mock-infected cell chromatin (set at 1). (**B**) Frequency distribution plots present the *T_50_* for GFP-H1.2 per individual cell. Solid or dashed vertical black lines, 1 SD below (RC) or above (Infected Cell Chromatin) the average *T_50_* for GFP-H1.2 in mock-infected cell-or nucleolar-chromatin, respectively. The *T_50_* for GFP-H1.2 per individual cell in the mock-infected cell-or nucleolar-chromatin are plotted in both graphs for comparison. (**C**) Bar graph presents the average normalized level of free GFP-H1.2 expressed relative to the average normalized level in mock-infected cell chromatin (set at 1). (**D**) Frequency distribution plots show the percentage of free GFP-H1.2 per individual cell. Solid or dashed vertical black lines, 1 SD above the average level of free GFP-H1.2 in mock-infected cell-or nucleolar-chromatin, respectively. The level of free GFP-H1.2 per individual cell in the mock-infected cell-or nucleolar-chromatin are plotted in both graphs for comparison. (**E**) Bar graph presents the average initial normalized fluorescence recovery rate for GFP-H1.2 expressed relative to the average initial normalized fluorescence recovery rate in mock-infected cell chromatin (set at 1). (**F**) Frequency distribution plots present the initial normalized fluorescence recovery rate of GFP-H1.2 per individual cell. Solid or dashed vertical black lines, 1 SD above the average initial normalized fluorescence recovery rate of GFP-H1.2 in mock-infected cell-or nucleolar-chromatin, respectively. The initial normalized fluorescence recovery rate for GFP-H1.2 per individual cell in the mock-infected cell-or nucleolar-chromatin are plotted in both panels for comparison. Error bars, SEM. n≥38 cells per treatment from 4 independent experiments. Different letters denote *P*<0.01; matching symbols denote *P*<0.05. Statistical significance evaluated by ANOVA with post-hoc Tukey Kramer pair-wise analysis.

In contrast to mobilization within RCs, H1.2 mobility was decreased within domains enriched for infected-cell chromatin (Fig 12B). Reduced mobility of H1.2 tended to increase its *T_50_* to 124 ± 9 %, although significance was not achieved (Fig 13A, B). Regardless, 31% of cells had a *T_50_* greater than 1SD above the average *T_50_* in mock-infected cell chromatin, which is almost double the population expected were H1.2 not immobilized within it (Fig 13B). Although the overall mobility of H1.2 decreased, its free pool and fast chromatin exchange were similar within EHV-or mock-infected cell chromatin (97 ± 3% or 106 ± 4%, respectively; Fig 13 C-F, Tables 1 and 2). These data suggest that decreased H1.2 mobility within infected-cell chromatin reflects stabilization of H1.2 high-affinity, rather than low-affinity, chromatin interactions.

Together, these data show that EHV1 mobilized H1.2 within nuclear domains enriched for viral chromatin (RCs). Mobilization was apparently independent of infection progression, although as infection progressed H1.2 was less likely to be bound in chromatin within RCs. Thus, H1.2 may be less likely assembled in, or displaced from, EHV1 chromatin during robust viral transcription and DNA replication, and when all kinetic protein classes are expressed. Infection decreased H1.2 mobility within the infected-cell chromatin. This de-mobilization did not affect H1.2 chromatin residency or low-affinity chromatin interactions, suggesting that EHV1 may further stabilize higher-affinity H1.2 binding within infected-cell chromatin.

## Discussion

Herein we show that EHV1 mobilized canonical and variant core histones and linker histone H1.2. Histones were most mobile within RCs, which are enriched for viral chromatin. The distinctive mobilities of histones within RCs or the surrounding cellular chromatin indicate that histone associations with viral chromatin are different from their associations with cellular chromatin and suggest that viral and cellular chromatin have unique properties. Moreover, the dissimilar mobility of histones within EHV1 RCs and the surrounding cellular chromatin supports the idea that histone assembly and turnover are differentially regulated in either chromatin. The particular factors that regulate such processes within the viral chromatin are therefore likely enriched in RCs.

The differential mobilization of histones within EHV1 RCs suggests that their mobilization is unlikely a general response to indiscriminately disrupt chromatin. Histone mobility was primarily enhanced through changes in chromatin residency or low-affinity turnover. Mobility of most histones within EHV1 RCs reflected a trend in which chromatin residency was most affected at earlier stages of infection, whereas both chromatin residency and low-affinity turnover were affected at later stages (Figs 5 and 9). Exceptions to this trend were mobilization of H3.1, H3.3, and H1.2 (Figs 7 and 13). These data are consistent with specific mechanisms that regulate chromatin assembly and turnover of individual histones. The magnitude of mobilization for individual histones were also variable. Some, such as H2A.X, H2A.Z, and H1.2, were preferentially mobilized at earlier stages of infection, whereas others, such as H2A, H3.1, and H3.3, were preferentially mobilized at later stages. Thus, the chromatin-binding and –turnover properties of individual histone types were differentially altered relative to each other and their relative mobilities varied over the course of infection. Any given histone may be mobilized to promote its assembly in viral chromatin. Conversely, mobilization may reflect eviction from, or inhibition of assembly in, EHV1 chromatin. The varied mobilities of individual histones over the course of infection is consistent with changes to their associations with viral chromatin and suggest that viral chromatin composition may also be variable.

Our data did not reveal significant differences in histone mobility during infection with neurotropic or non-neurotropic EHV1. Nonetheless, several trends indicate at least some differences in the histone mobilization mechanisms or histone-chromatin interactions of either strain. For example, during neurotropic EHV1 infection H2A.Z and H3.1 fast chromatin exchange tended to be slower in “small” RCs and faster in “large” RCs than during non-neurotropic EHV1 infection. Neurotropic EHV1 also tended to enhance H3.3 and H4 fast chromatin exchange in the cell chromatin surrounding “large” RCs.

Despite such trends, free histone pools were consistently similarly altered during infection with either neurotropic or non-neurotropic EHV1. The more prominent or variable alteration of fast chromatin exchange of particular histones during neurotropic EHV1 infection may reflect more specific nuances of viral chromatin regulation that are not readily apparent by FRAP analysis. Knowledge of histone mobilization mechanisms will support more in-depth characterizations of EHV1 chromatin and its regulation to provide insight into whether chromatin regulation varies between, or contributes to, strain pathotype. Regardless of whether neurotropic or non-neurotropic chromatin regulation is similar or distinct, understanding how viral chromatin is regulated will support the design of novel antiviral therapies to silence EHV1 genomes and prevent productive infection.

Histones were typically mobilized to a greater degree when infection had further progressed (based on identifiable RC size or number). However, given the similar appearance of “small” RCs and histone depleted regions within mock-infected cells, it is possible that that histone mobility within “small” RCs is greater than measured due to inadvertent selection of non-RC nuclear regions. If this were the case, analyses of histone mobility within any given “small” RC per individual cell would be expected to produce a bimodal frequency distribution, with subpopulations that have histone mobilities resembling those within mock-infected cell chromatin or “large” RCs. Only two analyses revealed bimodal distributions for histone mobility within “small” RCs, the fast chromatin exchange of H2B and H2A.X (Figs 5E and 9D). All other analyses of mobility within a “small” RC per individual cell revealed unimodal distributions intermediate to histone mobilities within mock-infected cell chromatin or “large” RCs. These data are consistent with a mixed population of infected cells that have mobilized a particular histone to varying extents within any given “small” RC. This mobilization supports a model in which the magnitude of mobilization relates to EHV1 replication properties that become more prominent as infection progresses. Thus, levels of EHV1 transcription or DNA replication, expression or accumulation of specific viral proteins, or cellular responses to them may further enhance histone mobility. This proposed model is consistent with our understanding of HSV1 mobilization of histones, for which this phenomenon of histone mobilization during viral infection was first described (48). All HSV1 transcription activators VP16, ICP0, and ICP4 mobilize histones, although ICP4 mobilizes them to the greatest extent and is sufficient to do so (48–50, 60). The EHV1 ICP4 homologue, IE1, has some conservation with ICP4 a.a. sequence, particularly within the DNA binding domain, and with some transcription regulatory mechanisms (71–75). However, whether the function of ICP4 to mobilize histones is conserved in IE1 is not yet known. Work is underway to characterize histone mobilization mechanisms and test how they relate to EHV1 infection progression.

Histones were predominantly depleted from RCs. This depletion is consistent with mobilization of histones specifically within RCs as, at any given time, histones were more likely imaged within cellular chromatin domains where they are less mobile and accordingly spend more time. However, histones were not universally depleted from RCs. Minor subpopulations of cells had RCs relatively less depleted for H3.1, H3.3, or macroH2A (Figs 6 and 8). Furthermore, diffuse H2A.B or diffuse H3.1 or H3.3 within a minor subpopulation of cells revealed that in some cases histones may be equally enriched, and spend equal time, within domains containing cellular or viral chromatin (Figs 6 and 10). The high degree of H3.1 or H3.3 mobilization in cells with diffuse H3 was unlikely due to general H3 eviction from all chromatins as nucleoli remained depleted for either histone (Fig 6). Thus, mechanisms to mobilize H3 to such an extreme degree within the viral or infected-cell chromatin did not notably alter its association or turnover within nucleolar chromatin. Likewise, nucleoli remained enriched for H2A.B within cells of mobility groups 2, 3, and 4+ (Fig 10). H2A.B association and turnover within nucleolar chromatin is thus also distinct from that within viral or infected-cell chromatin. While H2A.B is considered the most dynamic core histone, the extreme mobility of H3.1 and H3.3 within cells with diffuse H3 was to a similar degree as H2A.B mobilization within mobility group 4+ (Tables 1 and 2). These data show that, at least under certain circumstances, other core histones can be equally as mobile as H2A.B.

Histones were more dynamic within medium-to large-sized EHV1 RCs than within nucleoli. The higher mobility of histones within RCs indicates that EHV1 chromatin is more unstable or dynamic than nucleolar chromatin. This observation is also consistent with our understanding of the unique HSV1 lytic chromatin. HSV1 nucleosomes are highly unstable. Accordingly, they have very low immunoprecipitation efficiency and viral genomes are hyperaccessible to nucleases (51, 62–64, 76–81). As reported herein for EHV1, canonical (H2A, H2B, H3.1, H4) and variant (H2A.X, macroH2A, H3.3) core histones are also more dynamic within HSV1 RCs than within mock-infected nucleoli (Conn & Schang, unpublished). Although the extreme mobility of histones within HSV1 RCs is most consistent with the assembly of viral genomes in highly dynamic and unstable nucleosomes, whether histone mobility contributes to the remarkable hyperaccessibility of HSV1 chromatin is yet to be tested. HSV1 chromatin composition may contribute more to its unique hyperaccessibility than dynamic histone exchange. Consistently, H2A.B chromatin enrichment is associated with the DNA hyperaccessibility that is characteristic of HSV1 or nucleolar chromatin. Although histone mobilization is consistent with nucleosome instability, whether EHV1 chromatin is also hyperaccessible or enriched in H2A.B is not yet known. H2A.B enrichment within HSV1 chromatin is accompanied by its relative de-mobilization and accumulation in RCs (51). Similarly, H3.1 is relatively de-mobilized and can accumulate in RCs concomitant with its assembly in HSV1 chromatin (50, 82). We did not observe notable enrichment of any histone within EHV1 RCs. Nor was any histone relatively de-mobilized, or not mobilized, within RCs during EHV1 infection. Thus, histone mobility on its own did not highlight preferential assembly of any histone within EHV1 chromatin. This distinction suggests that EHV1 and HSV1 lytic chromatins have distinct properties or some degree of unique regulation. Work is underway to evaluate EHV1 chromatin composition and biophysical properties to further test the relationships between viral nucleosome instability, DNA hyperaccessibility, and histone mobility.

The dysregulation of histone chromatin exchange, with enhanced mobility in viral RCs, is a feature conserved between EHV1 and HSV1 (48–51, 60). However, differences in histone mobilization during infection with either virus are apparent. Most striking is the distinct mobilization of H2A.B. In HSV1-infected cells, H2A.B accumulates within viral RCs and is relatively de-mobilized concomitant with enrichment in transcriptionally active HSV1 chromatin (51). Conversely, H2A.B was neither depleted nor enriched within EHV1 RCs and its mobility was enhanced within a subpopulation of cells (Figs 10 and 11). Human and equine H2A.B are the most dissimilar histones evaluated herein, sharing only 70% a.a. identity (S1 Fig). Thus, structural differences, different PTMs, or different protein-protein interactions may account for distinct mobilizations of H2A.B during EHV1 or HSV1 infections. The apparent maintenance of nucleoli during EHV1 infection is potentially another contributing factor to differential H2A.B mobilization. Herein we show that nucleoli are largely morphologically similar within EHV1-or mock-infected cells (Figs 1, 2, 6, 8, 10, 12). In contrast, HSV1 morphologically and compositionally disrupts nucleoli (83–88). Consequently, many abundant nucleolar resident proteins, including nucleophosmin (NPM1, B23), fibrillarin, nucleolin, and upstream binding factor (UBF), redistribute throughout HSV1-infected nuclei and some accumulate within RCs (83–87, 89–93). Nucleolar protein redistribution kinetics, re-localization patterns, and the roles of specific HSV1 proteins (or not) in their redistribution highlight that multiple mechanisms contribute to disrupt nucleoli during HSV1 infection. At least one HSV1 protein, UL24, has conserved sequence identity with its EHV1 homologue, ORF37, in key regions necessary to disrupt nucleoli (83, 92, 94). Whether this function is conserved in ORF37 and whether individual nucleolar components, aside from H2A.B (Fig 10) or EAP (EBV-encoded small nuclear RNA-associated protein) (95), redistribute throughout the EHV1-infected nucleus are not yet tested. HSV1-mediated nucleoli disruption could displace H2A.B from nucleolar chromatin to promote its assembly in viral chromatin. Conversely, maintenance of nucleoli during EHV1-infection may maintain H2A.B association with nucleolar chromatin so that it is less available for assembly, or less likely to stably assemble, within EHV1 chromatin.

The apparent maintenance of nucleoli during EHV1 infection may relate to yet another difference between EHV1 and HSV1 mobilization of histones. EHV1 did not substantially mobilize histones in the infected-cell chromatin. Only H2A, H3.3, and H4 were partially mobilized within the cellular chromatin surrounding EHV1 RCs (Figs 5, 7, 9). The mobility of H2A, H3.3, and H4 within infected-cell chromatin reflected unique changes to individual chromatin interactions, their chromatin residency (H4 and H3.3, Figs 5A and 7A) or low-affinity chromatin exchange (H4 and H2A, Figs 5D and 9C). The differential mobilization of H2A, H3.3, and H4 within EHV1-infected cell chromatin more likely reflects distinct mechanisms to regulate chromatin exchange of individual histone types rather than a consequence of general nucleosome disruption. In contrast, HSV1 more broadly disrupts the cellular chromatin surrounding RCs. A similar analysis in HSV1-infected cells revealed that all evaluated canonical (H2A, H2B, H3.1, H4) and variant (H2A.X, macroH2A, H3.3) core histones were mobilized to increase their free pools within the infected cell chromatin (Conn & Schang, unpublished). It is interesting to note that histones are broadly mobilized within the cellular chromatin during HSV1 infection despite its progressive compaction and marginalization as infection progresses (96). Two nucleolar resident proteins that redistribute throughout HSV1-infected nuclei, nucleolin and NPM1, are histone chaperones that disrupt nucleosomes to exchange H2A-H2B (nucleolin) or H3-H4 (NPM1) dimers (83, 84, 87, 92, 93, 97–100). Additionally, nucleolin and NPM1 promote nucleosome remodeling and interact with H1 to further decondense chromatin and facilitate histone mobilization (97–99, 101, 102). Thus, increased abundance of nucleolin and NPM1 throughout the HSV1-infected cell chromatin may account for, or contribute to, general mobilization of core histones within it. Conversely, morphological maintenance of nucleoli within EHV1-infected cells suggests that nucleolar resident proteins may not be dispersed, or as drastically dispersed, as they are during HSV1 infection. It remains to be tested whether individual nucleolar proteins, particularly those with histone chaperone activities, relocalize during EHV1 infection and, if so, how they may contribute to histone mobilization.

Herein, we provide a comprehensive analysis of histone mobility in equine cells that is the first to comparatively evaluate mobility within nucleolar and non-nucleolar chromatin. Importantly, as previous investigations of histone mobility used primate-or murine-derived cells, our report of histone mobility within equine-derived cells provides a unique perspective and opportunity to identify processes that may be distinct or differentially regulated in a more evolutionarily distant mammalian species. The mobility of most core histones evaluated herein were as expected based on their known effects on nucleosome stability and, when available, reported mobilities. Histone mobility was also most consistent with our understanding of the relationships between chromatin stability and histone exchange in that histones were more mobile within nucleolar than non-nucleolar chromatin. It is interesting to note that H2A variants, aside from H2A.B, were quite similarly mobile within nucleolar chromatin despite distinct mobilities within non-nucleolar chromatin (S3 Fig, S2 Table). This observation further highlights the unique regulation and properties of nucleolar chromatin. Despite the expected mobilities of most core histones, two examples of remarkable histone dynamics were apparent, the multiple mobilities of H2A.B and the highly dynamic H3.1 population. To our knowledge, this is the first study to comparatively evaluate H2A.B mobility within nucleolar and non-nucleolar chromatin and the first to report distinct H2A.B mobilities within either chromatin, as well as variable H2A.B mobilities within non-nucleolar chromatin (Fig 10, S5 Fig). The distinct mobilities of H2A.B reported herein may not have been noted in other studies due to measuring H2A.B mobility across larger nuclear regions containing variable proportions of nucleolar or non-nucleolar chromatin (36, 51, 69). H2A.B mobility within nucleoli was remarkably consistent across all cells evaluated, indicating that H2A.B exchange in nucleolar chromatin is tightly regulated (Fig 10, S5 Fig). Its distinct and consistent mobility within nucleoli also suggests that factors to mediate or regulate H2A.B turnover within nucleolar chromatin primarily do so within this domain. Conversely, the multiple and variable mobilities of H2A.B within non-nucleolar chromatin are consistent with multiple mechanisms to regulate its assembly or turnover within this chromatin and suggest that H2A.B is subjected to differential regulation within it. The basic arginine repeat region of human H2A.B, which is thought to have regulatory functions, is truncated from 6 to 4 residues within equine H2A.B (S1 Fig). Moreover, equine H2A.B has a serine residue within this region that may be subject to PTM. The sequence differences within the N-terminus of equine H2A.B may contribute to alternate regulation of its chromatin assembly or turnover within non-nucleolar chromatin.

The observation that a population of H3.1 is more dynamic than H3.3 within the cellular chromatin was also unexpected (S2 Fig, S2 Table). This is the first study to comparatively evaluate H3.3 and H3.1 chromatin dynamics using FRAP, which is a sensitive technique to directly evaluate dynamic histone populations. Another study that used FRAP to measure GFP-H3.3 or –H3.1 mobility within Vero (African green monkey) cells shows the same dynamic H3.1 population, although H3.3 and H3.1 mobilities were not directly compared(50). The reported normalized initial fluorescence intensity after photobleaching is approximately 25% or less than 20% for GFP-H3.1 or GFP-H3.3, respectively, within Vero cells, which is comparable to the 25% or 21% reported herein (Table 1) (50). Other methods to indirectly measure H3.1 or H3.3 mobility typically use labelled histones to measure their enrichment in or depletion from specific chromatin regions over time. Such studies do not encounter the most dynamic histone populations as highly dynamic histones are less efficiently fixed and are lost during nuclei preparations. Thus, studies that indirectly measure H3.1 or H3.3 mobility likely have not encountered this most dynamic H3.1 population consequent to methodologies used.

In summary, we show that neurotropic and non-neurotropic EHV1 mobilize histones. Linker histone H1.2 and all canonical and variant core histones, with exception of H2A.B, are mobilized within nuclear domains enriched for EHV1 chromatin. Histones are further mobilized as infection progresses, consistent with increased levels of EHV1 transcription, DNA replication, or protein expression directly or indirectly enhancing histone mobility. The high degree of histone mobilization within EHV1 chromatin domains is consistent with the assembly of viral genomes in dynamic and unstable nucleosomes. Although histones are mobilized within domains enriched in viral chromatin, they are not within domains enriched in cellular chromatin. Histones may therefore be preferentially mobilized within viral chromatin to regulate its stability. The manipulation of histone dynamics is a novel epigenetic mechanism conserved among EHV1 and HSV1 that directly regulates viral chromatin stability and genome accessibility.

## Materials and methods

### Cells and virus

EDerm cells (NBL-6; ATCC) were maintained in Dulbecco’s modified minimum Eagle’s medium (DMEM, Gibco 11885-084) supplemented with 10% fetal bovine serum (FBS, Corning 35-077-CV) at 37°C in 5% CO_2_. Equid alphaherpesvirus 1 (EHV1) strains R08-8428 and D08-8315 were generous gifts from Dr. Vikram Misra (University of Saskatchewan). Strain D08-8315 has ORF 30 G2254 (D752) associated with neurological EHV1 pathotype. This strain was isolated from a 17-year-old female quarter horse with neurological deficits (Prairie Diagnostic Services (PDS) Saskatoon). Strain R08-8428 has ORF 30 A2254 (N752) associated with non-neurotropic EHV1 pathotype. No information regarding this strain isolation is available (PDS Saskatoon).

### Viral stock preparation and titration

Viral stocks were prepared and titrated in EDerm cells. Briefly, cells were seeded such that they were approximately 40% confluent at the time of infection. Cells were typically infected with 0.01 to 0.05 plaque forming units (PFU) per cell diluted in a minimal volume of 4°C DMEM. Following inoculum addition, cells were incubated at 37°C in 5% CO_2_ for 1h with rocking and rotating every 5-10min. The inoculum was then removed and cells were washed twice with 4°C phosphate buffered saline (PBS; 150mM NaCl, 1mM KH_2_PO_4_, 3mM Na_2_HPO_4_, pH 7.4) prior to the addition of fresh 37°C DMEM supplemented with 10% FBS. Cells incubated at 33°C in 5% CO_2_ until greater than 95% cytopathic effect (CPE) was observed. Infected cells were then harvested and pelleted by centrifugation at 3,214xg for 20min at 4°C. Extracellular virions were isolated from the resulting supernatant by centrifugation at 10,000xg for 2h at 4°C. Intracellular virions were released from the cell pellet by the disruption of cell membranes through three rapid freeze-thaw cycles in a dry ice-EtOH bath and 37°C water bath, respectively. Cellular debris was then pelleted by centrifugation at 5,500xg for 30min at 4°C. Intra– and extracellular virions were combined for the EHV1 stocks.

For stock titration, EDerm cells were seeded in 12-well tissue culture plates such that they would be 50-60% confluent at the time of infection. The EHV1 stock was serially diluted 10-fold in 4°C DMEM and cell monolayers overlayed with a minimum volume of the serial dilutions (typically 10^-3^ to 10^-6^). Cells incubated at 37°C in 5% CO_2_ for 1h with rocking and rotating every 5-10min. The dilutions were then removed, cells were washed twice with 4°C PBS and then overlayed with 37°C 0.5% (w/v) methylcellulose in DMEM supplemented with 5% FBS. Infected cells incubated at 37°C in 5% CO_2_ until well-defined plaques were visible in the cell monolayer (typically 3-5 days). Cells were then fixed and stained by addition of 1% (w/v) crystal violet in 17% (v/v) MeOH and incubated at room temperature for a minimum of 24h. Titrations were washed by gentle agitation in a bucket of ambient water and air dried at room temperature prior to counting plaques.

### Plasmids

Constructs encoding green fluorescent protein (GFP) fused to the amino (N)-terminus of histones H2A.X, H2B, H3.3, H3.1, and H4, and GFP fused to the carboxyl (C)-terminus of H2A.Z were generous gifts from Dr. Luis Schang (Cornell University) (49–51, 60). The amino acid sequences of human and equine histones H2A.Z.1 (NP_002097, XP_023493427), H2B (NP_003514, XP_001497452), H3.3 (NP_001365974, NP_001356113), H3.1 (NP_001368928, NP_001356115), and H4 (NP_001029249, XP_001496752) are identical. The DNA sequence encoding H2A.Z.1 (H2A.Zv) was PCR amplified from pH2A.Z.1-GFP using primers H2A.Z F and H2A.Z R (Table 4) with flanking BglII and HindIII restriction sites, respectively. Amplified DNA was ligated in-frame with GFP in pEGFP-C1 (a generous gift from Dr. Luis Schang, Cornell University) to create pEGFP-H2A.Z. Human (NP_002096) and equine (XP_023500737) histone H2A.X differ at amino acid 131. H2A.X human 131S was changed to equine 131A by site-directed mutagenesis of pEGFP-H2A.X using primers H2A.X S131A F and H2A.X S131A R (Table 4).

**Table 4.**
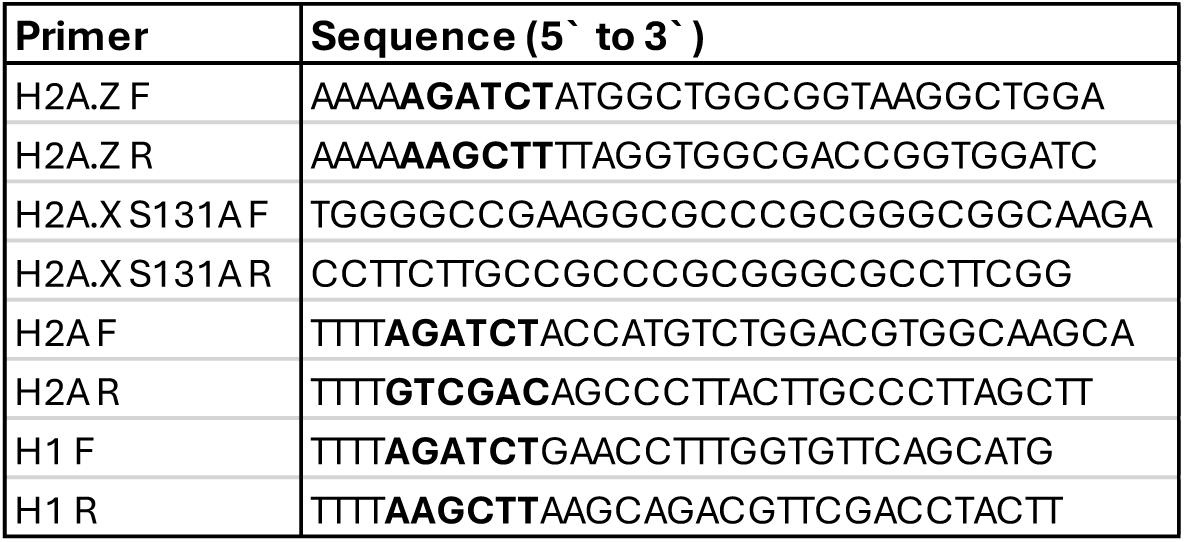
PCR primer sequences. Restriction enzyme recognition sequences are bolded.

Human and equine histone H2A are identical (NP_003501, XP_001505083). The sequence encoding H2A was PCR amplified from U2OS (human osteosarcoma; ATCC HTB-96) genomic DNA using the primers H2A F and H2A R (Table 4). Amplified DNA was ligated in-frame with GFP in pEGFP-C1 using BglII and SalI restriction sites to create pEGFP-H2A. DNA sequences encoding equine macroH2A (XP_023473434.1) and equine H2A.B (XP_023489462.1) were ordered as gBlocks from Integrated DNA Technologies (IDT; Table 5). Flanking BglII and KpnI (macroH2A) or BglII and HindIII (H2A.B) restriction sites were used for in-frame ligation with GFP in pEGFP-C1 to create pEGFP-macroH2A or pEGFP-H2A.B, respectively. The DNA sequence encoding equine linker histone H1.2 (XM_005603676) was PCR amplified from EDerm genomic DNA using the primers H1.2 F and H1.2 R (Table 4). Amplified DNA was ligated in-frame with GFP in pEGFP-C1 using BglII and HindIII restriction sites to create pEGFP-H1.2. All plasmid sequences were confirmed by Sanger sequencing (Sanger Sequencing Platform, CRCHU de Québec-Université Laval, CHUL).

**Table 5.**
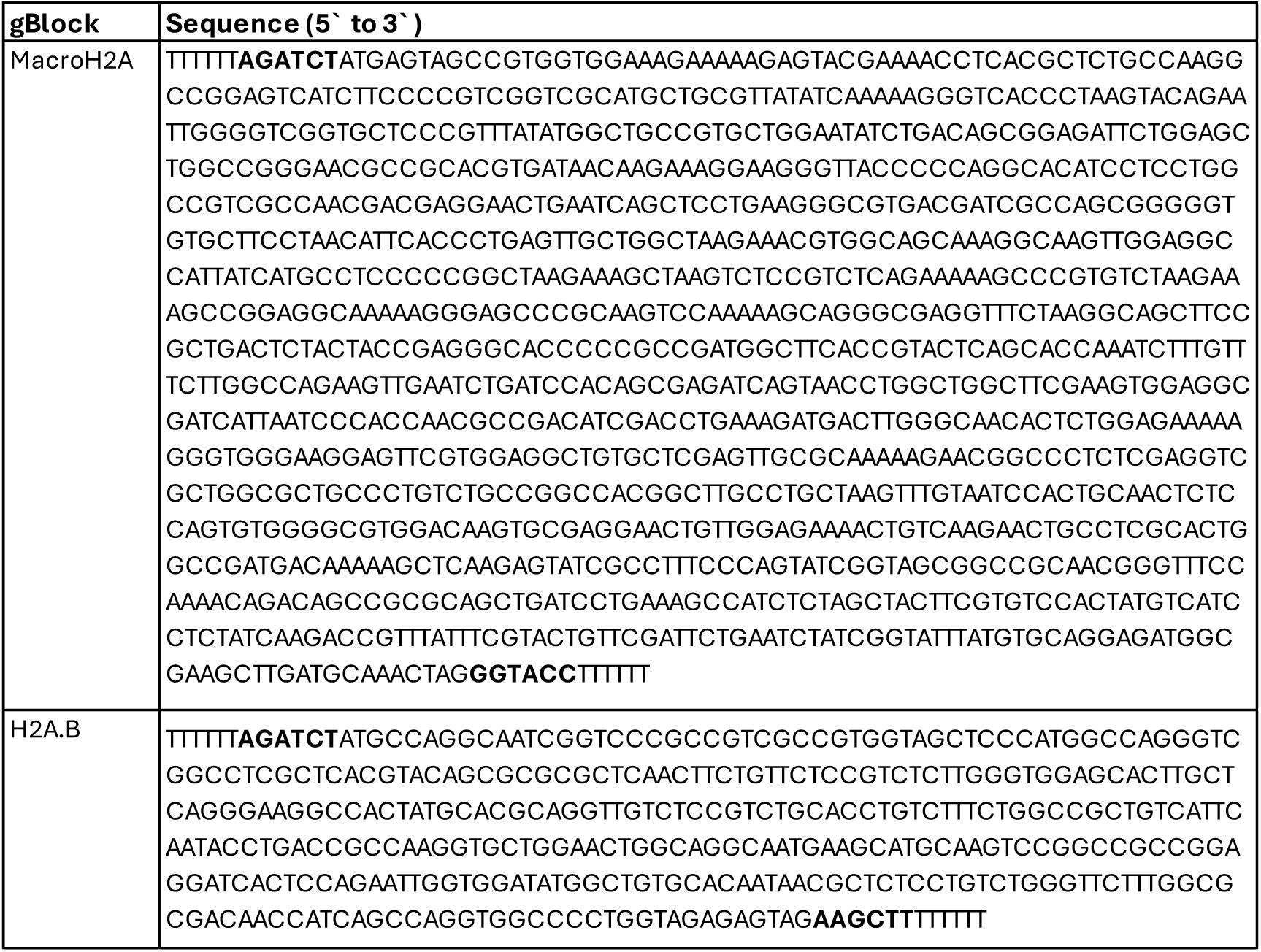
Synthetic DNA sequences. Restriction enzyme recognition sequences are bolded.

Amino acid sequence comparison of equine and human histones H2A.B, H2A.X, macroH2A, and H1.2 are depicted in S1 Fig.

### Transfection

EDerm cells (2.0-2.6×10^5^) were seeded in 6-well tissue culture plates at least 12h prior to transfection. Plasmid DNA was transfected using Lipofectamine 3000 Transfection Reagent (Invitrogen, L3000001). Briefly, for each well to be transfected 4-6µg of plasmid DNA were combined with 4-6µl of P3000 reagent in 100µl of 4°C DMEM in a microfuge tube. In a separate microfuge tube, 1µl of Lipofectamine 3000 was added to 100µl of 4°C DMEM.

Following a 5min incubation at room temperature, the plasmid DNA-P3000 mix was added to the Lipofectamine mix. The combined transfection mix incubated at room temperature for an additional 30min before the volume was brought to 800µl with room temperature DMEM. Cell medium was removed and the cells overlayed with the transfection mix. Cells incubated at 37°C in 5% CO_2_ for 5-6h, then the transfection mix was removed and replaced with fresh 37°C DMEM supplemented with 10% FBS. Transfected cells incubated in 5% CO_2_ at 37°C at least 36h prior to seeding for infection and FRAP analysis. Transiently expressed exogenous histones with N-terminal GFP fusions are nuclear, assemble in chromatin, and have lower expression levels than endogenous histones (48–51, 60).

### EHV1 infection

Transfected EDerm cells were seeded (4×10^5^) on 18×18mm coverslips in 6-well plates for fluorescence recovery after photobleaching (FRAP) analysis. Seeded cells incubated at 37°C in 5% CO_2_ at least 7h prior to infection. Inoculum was prepared by diluting purified EHV1 stocks in 4°C DMEM. For mock infections, 4°C DMEM was used. Cells were overlaid with 400µl of inoculum containing 10 PFU/cell of EHV1 strain R08-8428 (abortogenic) or strain D08-8315 (neurotropic) and incubated at 37°C in 5% CO_2_ for 1h with rocking and rotating every 5-10min. The inoculum was then removed, cells were washed twice with 4°C PBS and overlayed with fresh 37°C DMEM supplemented with 10% FBS. Infected cells incubated at 37°C in 5% CO_2_ until they were subjected to FRAP analysis.

### FRAP

Histone mobility was evaluated between 5 and 6hpi as described previously (48–50, 60). Briefly, a coverslip was mounted on a slide and put on a 37°C stage on a Zeiss LSM 700 inverted confocal microscope(48). Cells were viewed using a Plan-Apochromat 40×/1.4 oil DIC objective lens. FRAP was performed using an Argon laser (488nm) with maximum pinhole size. Whole cell imaging was performed at 2-3% laser intensity while photobleaching was achieved with 30-35 iterations at 100% laser intensity. Two circular regions of equal volume, one within the nucleoli or EHV1 replication compartment of mock-or EHV1-infected cells, respectively, and one within the surrounding cellular chromatin of each were photobleached (as depicted in Fig2A). Thirty to forty-five differential interference contrast (DIC) and fluorescent images were collected at 0.9s intervals from before photobleaching and after photobleaching. At each interval the fluorescence of the entire cell nucleus was measured. The fluorescence of the photobleached regions at each time were normalized to the total nuclear fluorescence at that same time. The normalized fluorescence of each photobleached region at any time is presented as a ratio to the normalized fluorescence of the same region before photobleaching. The fluorescence of the photobleached regions recover as bleached GFP-histones from within them undergo chromatin exchange with the non-bleached fluorescent GFP-histones from outside them. FRAP was measured for 30-45s as we were interested in evaluating the most dynamic histone populations. Moreover, any potential contribution of newly synthesized and nuclear imported GFP-histones to the fluorescence recovery is negated by such a short time. A single slide was used for FRAP analysis for less than 1h. Typically 8-10 cells per condition (mock-, R08-8428-or D08-8315-infection) from at least 4 independent experiments were evaluated.

The normalized fluorescence intensity of the photobleached nuclear regions at the first time after photobleaching was used as a surrogate measure for the level of histones available in the free pool (not bound in chromatin and diffusing in the nucleoplasm). The slope between the normalized fluorescence at the first and second times after photobleaching, representing the initial rate of fluorescence recovery, was used as a surrogate measure for the fast chromatin exchange rate (those histones weakly bound in chromatin and undergoing low-affinity chromatin exchange). The slope of normalized fluorescence recovery from 15-30 s after photobleaching was used as a surrogate measure for the slow chromatin exchange rate (those histones stably bound in chromatin and undergoing high-affinity chromatin exchange). For linker histone H1.2, mobilization was also evaluated by the time to recover 50% of the normalized fluorescence in the photobleached regions (*T_50_*). A higher degree of mobilization results in a shorter *T_50_* as mobilization and *T_50_* inversely relate.

For each individual experiment the level of free histone, calculated fast or slow chromatin exchange rates, or *T_50_* (for H1.2) for each mock-, R08-8428-, or D08-8315-infected cell were normalized to their average values for the mock-infected cell chromatin from the same experiment to account for any potential differences in photobleaching efficiency on any given day.

## Image preparation

Fluorescent images were analyzed using Zeiss Zen black. DIC and fluorescent image brightness and contrast were altered for figure preparation using FIJI(103).

## Statistical analysis

Statistical significance was tested using single-factor ANOVA. For comparisons where ANOVA identified differences, samples were evaluated pairwise post hoc using Tukey Kramer analysis to identify those samples that differed. Student’s two-tailed T-test was used for pairwise comparison of canonical and variant histone mobilities.

## Acknowledgements

We acknowledge University of Saskatchewan Health Sciences 6^th^ floor Virology Cluster for use of lab space and cell culture facilities, and the University of Saskatchewan Cancer Cluster for use of the Zeiss LSM 700 microscope.

