## Supplemental Information for "Neurotropic and non-neurotropic equid alphaherpesvirus 1 (EHV1) mobilize most histones within viral replication compartments"

### Supporting figure captions

**S1 Fig. Sequence comparison between equine and human histones H2A.B, H2A.X, macroH2A, and H1.2.** Amino acid sequences for equine or human histones H2A.B. (H2A Bbd type 2/3; XP_023489462.1 or NP_001017991, respectively), H2A.X (XP_023500737 or NP_002096, respectively), macroH2A (macro-H2A.1 isoform X2 XP_023473434.1 or macro-H2A.1 isoform 2 NP_001035248.1, respectively), and H1.2 (XP_005603676.1 or NP_005310, respectively) were pair-wise aligned using MUSCLE Alignment in Geneious Prime 2025.1.2. Identical residues are indicated by a dot. The conserved histone fold of core histones, encompassing three alpha helices and two interjoining loop regions, is highlighted in pink. The docking domain that mediates H2A-H2B dimer binding to the H3-H4 tetramer is highlighted for the H2A variants in blue. Human H2A.B has an amino-terminal arginine motif (RRR) that is shortened in equine H2A.B (highlighted in purple). The H2A.X residues phosphorylated in response to DNA damage (T136, S139) are conserved among equine and human (denoted by red asterix)(23). The macro domain of macroH2A (highlighted in green) is largely conserved between equine and human. The conserved DNA-binding winged-helix motif of linker histones, encompassing three alpha helices and a beta-sheet "wing", is denoted in brown. Amino acid residues for the highlighted regions are as denoted for human histones on HistoneDB 2.0(104).

**S2 Fig. H3.1 is most dynamic in mock-infected cell chromatin, whereas H3.3 is most dynamic in "large" EHV1 RCs.** EDerm cells were transfected with plasmids encoding GFP fused to H3.1 or H3.3 at least 40h prior to mock-infection or infection with 10 PFU/cell of abortogenic or neurotropic EHV1. Nuclear mobilities of GFP-H3.1 or -H3.3 were evaluated by FRAP between 5 and 6hpi. FRAP data for EHV1 infected cells were pooled for each histone and segregated by the presence (Large RC; Large) or absence (Small RC; Small) of clearly identifiable RCs, or diffuse H3 (Diffuse). (**A**) Line graphs present the FRAP of GFP-H3.1 (black) or -H3.3 (blue) in mock- or EHV1-infected cells. (**B**) Bar graph presents the average normalized level of free GFP-H3.3 expressed relative to the average normalized level of free GFP-H3.1 in the same compartment (set at 1). (**C**) Bar graph presents the average initial normalized fluorescence recovery rate for GFP-H3.3 expressed relative to the average initial normalized fluorescence recovery rate for GFP-H3.1 in the same compartment (set at 1). Error bars, SEM; dashed error bars represent SEM for H3.1. H3.1 $n \geq 46$ cells per treatment from 5 independent experiments; H3.3 $n \geq 60$ cells per treatment from 6 independent experiments. ** $P<0.01$; * $P<0.05$; ns, not significant, Student's two-tailed T-test pairwise comparison of variant H3.3 to canonical H3.1 for each.

#### S3 Fig. Variant H2A histones are differentially mobilized in EHV1 RCs.

EDerm cells were transfected with plasmids encoding GFP-H2A, -H2A.Z, -H2A.X, -macroH2A, or -H2A.B. At least 40h after transfection, cells were mock-infected or infected with 10 PFU/cell of abortogenic or neurotropic EHV1. Nuclear mobilities of GFP-H2A, -H2A.Z, -H2A.X, -macroH2A, or -H2A.B were evaluated by FRAP between 5 and 6hpi. FRAP data for EHV1 infected cells were pooled for each histone and segregated by the presence (Large RC; Large) or absence (Small RC; Small) of clearly identifiable RCs for histones H2A, H2A.Z, H2A.X, or macroH2A, or by mobility group for histone H2A.B. (**A**) Line graphs present the FRAP of GFP-H2A, -H2A.Z, -H2A.X, or -macroH2A in mock- or EHV1-infected cells. (**B**) Bar graphs present the average normalized level of free GFP-H2A.Z, -H2A.X, -macroH2A, or -H2A.B (mobility groups 1 to 4+) expressed relative to the average normalized level of free GFP-H2A in the same compartment (set at 1). For EHV1 infected cells, H2A.B mobility groups 1 to 4+ are expressed relative to the levels of free H2A in the infected-cell chromatin and large RC for comparison. (**C**) Bar graphs present the average initial normalized fluorescence recovery rate for GFP-H2A.Z, -H2A.X, -macroH2A, or -H2A.B (mobility groups 1 to 4+) expressed relative to the average initial normalized fluorescence recovery rate for GFP-H2A in the same compartment (set at 1). For EHV1 infected cells, H2A.B mobility groups 1 to 4+ are expressed relative to the average initial normalized fluorescence recovery rate of H2A in the infected-cell chromatin and large RC for comparison. Error bars, SEM. $n \geq 38$ cells per treatment from 4 independent experiments. ** $P<0.01$; * $P<0.05$; ns, not significant, Student's two-tailed T-test for pairwise comparison of variant H2A to canonical H2A for each.

**S4 Fig. Canonical and variant H2A free pool levels do not relate to GFP-histone expression levels.** EDerm cells were transfected with plasmids encoding GFP-H2A, -H2A.Z, -H2A.X, -macroH2A, or -H2A.B. At least 40h after transfection, cells were mock-infected or infected with 10 PFU/cell of abortogenic or neurotropic EHV1. Nuclear mobilities of GFP-H2A, -H2A.Z, -H2A.X, -macroH2A, or -H2A.B were evaluated by FRAP between 5 and 6hpi. FRAP data for EHV1 infected cells were pooled for each histone and segregated by the presence (Large) or absence (Small) of clearly identifiable RCs for histones H2A, H2A.Z, H2A.X, or macroH2A, or by mobility group for H2A.B (1, 2, 3, 4+). Dot plots present the level of free GFP-H2A, -H2A.Z, -H2A.X, -macroH2A, or -H2A.B per individual cell plotted against its normalized total nuclear fluorescence intensity prior to photobleaching. $n \geq 38$ cells per treatment from 4 independent experiments.

**S5 Fig. Variant histone H2A.B is differentially mobilized in non-nucleolar chromatin.** EDerm cells were transfected with plasmids encoding GFP-H2A.B at least 40h prior to mock-infection or infection with 10 PFU/cell of abortogenic or neurotropic EHV1. Nuclear mobility of GFP-H2A.B was evaluated by FRAP between 5 and 6hpi. FRAP data for mock-infected cell chromatin or EHV1-infected cells were pooled and segregated by mobility group. Line graphs present FRAP of GFP-H2A.B in mock- or EHV1-infected cells. Error bars, SEM. $n \geq 38$ per treatment cells from 4 independent experiments.

**S6 Fig. Linker histone H1.2 free pool levels are independent of GFP-H1.2 expression levels.** EDerm cells were transfected with plasmids encoding GFP-H1.2. At least 40h after transfection, cells were mock-infected or infected with 10 PFU/cell of abortogenic or neurotropic EHV1. Nuclear mobility of GFP-H1.2 was evaluated by FRAP between 5 and 6hpi. FRAP data for EHV1 infected cells were pooled and segregated by the presence (Large) or absence (Small) of clearly identifiable RCs. Dot plot presents the level of free GFP-H1.2 per individual cell plotted against its normalized total nuclear fluorescence intensity prior to photobleaching. n≥38 cells per treatment from 4 independent experiments.

**S1 Table. Percent of EHV1-infected cells with "small" or "large" RCs.**

| | | % of cells with: | | |
|---|---|---|---|---|
| | | Small RCs | Large RCs | Diffuse |
| **H2B** | Abortogenic | 24 | 76 | - |
| | Neurotropic | 30 | 70 | - |
| **H4** | Abortogenic | 23 | 77 | - |
| | Neurotropic | 33 | 67 | - |
| **H3.1** | Abortogenic | 11 | 71 | 18 |
| | Neurotropic | 21 | 67 | 12 |
| **H3.3** | Abortogenic | 26 | 54 | 20 |
| | Neurotropic | 16 | 59 | 25 |
| **H2A** | Abortogenic | 29 | 71 | - |
| | Neurotropic | 13 | 87 | - |
| **H2A.Z** | Abortogenic | 8 | 92 | - |
| | Neurotropic | 23 | 77 | - |
| **H2A.X** | Abortogenic | 15 | 85 | - |
| | Neurotropic | 20 | 80 | - |
| **Macro H2A** | Abortogenic | 30 | 70 | - |
| | Neurotropic | 38 | 62 | - |
| **H1.2** | Abortogenic | 26 | 74 | - |
| | Neurotropic | 39 | 61 | - |

**S2 Table. Variant histone mobilities relative to canonical histone mobility.**

| | | | | | Free Histone | | | Fast Recovery Rate | | | Slow Recovery Rate | | |
|---|---|---|---|---|---|---|---|---|---|---|---|---|---|
| | | | | | Relative Level,% (avg ± SEM) | *P* | Cells with extreme increase[a], % | Relative Level,% (avg ± SEM) | *P* | Cells with extreme increase[a], % | Relative Level,% (avg ± SEM) | *P* | Cells with extreme increase[a], % |
| **H3.3** | **Mock** | Chromatin | H3.1 | | 100 ± 5 | - | 33 | 100 ± 12 | - | 13 | 100 ± 20 | - | 13 |
| | | | H3.3 | | 85 ± 5 | * | 39[b] | 60 ± 10 | * | 27[b] | 75 ± 8 | ns | 0[b] |
| | | Nucleolus | H3.1 | | 100 ± 4 | - | 17 | 100 ± 13 | - | 15 | 100 ± 21 | - | 11 |
| | | | H3.3 | | 103 ± 4 | ns | 17 | 56 ± 7 | ** | 18[b] | 120 ± 13 | ns | 5 |
| | **EHV** | Cell Chromatin | H3.1 | | 100 ± 4 | - | 18 | 100 ± 10 | - | 13 | 100 ± 11 | - | 13 |
| | | | H3.3 small | | 73 ± 5 | ** | 35[b] | 81 ± 8 | ns | 2[b] | 128 ± 14 | ns | 19 |
| | | | H3.3 large | | 105 ± 4 | ns | 14 | | | | | | |
| | | Small RC | H3.1 | | 100 ± 4 | - | 13 | 100 ± 16 | - | 14 | 100 ± 30 | - | 10 |
| | | | H3.3 | | 79 ± 4 | ** | 26[b] | 52 ± 8 | * | 46[b] | 275 ± 64 | * | 48 |
| | | Large RC | H3.1 | | 100 ± 3 | - | 19 | 100 ± 9 | - | 20 | 100 ± 15 | - | 16 |
| | | | H3.3 | | 113 ± 3 | ** | 30 | 84 ± 9 | ns | 17[b] | 135 ± 22 | ns | 19 |
| | | Diffuse | H3.1 | | 100 ± 4 | - | 21 | 100 ± 9 | - | 11 | 100 ± 14 | - | 14 |
| | | | H3.3 | | 112 ± 3 | * | 28 | 90 ± 5 | ns | 17 | 64 ± 9 | * | 45[b] |
| **H2A** | **Mock** | Chromatin | H2A | | 100 ± 3 | - | 19 | 100 ± 8 | - | 11 | 100 ± 9 | - | 16 |
| | | | H2A.Z | | 116 ± 3 | ** | 38 | 144 ± 12 | ** | 35 | 116 ± 10 | ns | 25 |
| | | | H2A.X | | 102 ± 3 | ns | 23 | 102 ± 7 | ns | 13 | 99 ± 14 | ns | 18 |
| | | | MacroH2A | | 91 ± 3 | ns | 28[b] | 95 ± 10 | ns | 21[b] | 110 ± 16 | ns | 15 |
| | | | H2A.B | 1 | 115 ± 11 | ns | 40 | 371 ± 54 | ** | 100 | | | |
| | | | | 2 | 142 ± 4 | ** | 83 | 728 ± 82 | ** | 100 | | | |
| | | | | 3 | 171 ± 7 | ** | 100 | 894 ± 83 | ** | 100 | | | |
| | | | | 4 | 194 ± 7 | ** | 100 | 1442 ± 131 | ** | 100 | | | |
| | | Nucleolus | H2A | | 100 ± 3 | - | 11 | 100 ± 8 | - | 15 | 100 ± 11 | - | 12 |
| | | | H2A.Z | | 107 ± 3 | ns | 15 | 101 ± 7 | ns | 8 | 121 ± 13 | ns | 26 |
| | | | H2A.X | | 107 ± 4 | ns | 23 | 92 ± 8 | ns | 15[b] | 116 ± 13 | ns | 30 |
| | | | MacroH2A | | 95 ± 3 | ns | 15 | 76 ± 7 | * | 16[b] | 139 ± 18 | ns | 36 |
| | | | H2A.B | | 74 ± 2 | ** | 56 | 567 ± 30 | ** | 100 | | | |
| | **EHV** | Cell Chromatin | H2A | | 100 ± 3 | - | 19 | 100 ± 5 | - | 19 | 100 ± 8 | - | 14 |
| | | | H2A.Z | | 127 ± 3 | ** | 28 | 131 ± 9 | ** | 27 | 130 ± 9 | * | 20 |
| | | | H2A.X | | 99 ± 3 | ns | 13[b] | 84 ± 5 | * | 18[b] | 103 ± 12 | ns | 8 |
| | | | MacroH2A | | 93 ± 3 | ns | 16[b] | 69 ± 5 | ** | 39[b] | 86 ± 7 | ns | 5[b] |
| | | | H2A.B | 1 | 116 ± 8 | ns | 30 | 234 ± 19 | ** | 90 | | | |
| | | | | 2 | 149 ± 5 | ** | 78 | 342 ± 16 | ** | 98 | | | |
| | | | | 3 | 149 ± 4 | ** | 80 | 348 ± 16 | ** | 95 | | | |
| | | | | 4+ | 238 ± 5 | ** | 100 | 493 ± 16 | ** | 100 | | | |
| | | Small RC | H2A | | 100 ± 5 | - | 20 | 100 ± 15 | - | 16 | 100 ± 14 | - | 22 |
| | | | H2A.Z | | 131 ± 8 | ** | 58 | 172 ± 26 | * | 45 | 69 ± 18 | ns | 19[b] |
| | | | H2A.X | | 133 ± 7 | ** | 64 | 136 ± 29 | ns | 21 | 144 ± 35 | ns | 43 |
| | | | MacroH2A | | 101 ± 4 | ns | 19 | 86 ± 8 | ns | 8[b] | 115 ± 20 | ns | 21 |
| | | Large RC | H2A | | 100 ± 2 | - | 17 | 100 ± 8 | - | 12 | 100 ± 10 | - | 16 |
| | | | H2A.Z | | 105 ± 2 | ns | 18 | 117 ± 9 | ns | 20 | 163 ± 17 | ** | 28 |
| | | | H2A.X | | 99 ± 2 | ns | 18 | 66 ± 6 | ** | 22[b] | 99 ± 11 | ns | 17 |
| | | | MacroH2A | | 89 ± 2 | ** | 30[b] | 64 ± 8 | ** | 32[b] | 117 ± 14 | ns | 17 |
| | | | H2A.B | 1 | 53 ± 4 | ** | 100 | 117 ± 10 | ns | 10 | | | |
| | | | | 2 | 67 ± 2 | ** | 85 | 179 ± 8 | ** | 59 | | | |
| | | | | 3 | 67 ± 2 | ** | 91 | 182 ± 8 | ** | 57 | | | |
| | | | | 4+ | 108 ± 2 | * | 27 | 257 ± 8 | ** | 96 | | | |

[a] percentage of cells > 1SD above the average value for the canonical histone in mock infected cell chromatin

[b] percentage of cells > 1SD below the average value for the canonical histone in mock infected cell chromatin

** *P* <0.01; * *P* <0.05

#### S3 Table. Percent of cells per H2A.B mobility group

| | | Mock (% of cells) | EHV1 Abortogenic (% of cells) | EHV1 Neurotropic (% of cells) |
|---|---|---|---|---|
| H2A.B Mobility Group | 1 | 13 | 0 | 13 |
| | 2 | 31 | 32 | 37 |
| | 3 | 38 | 21 | 29 |
| | 4/4+ | 18 | 47 | 21 |

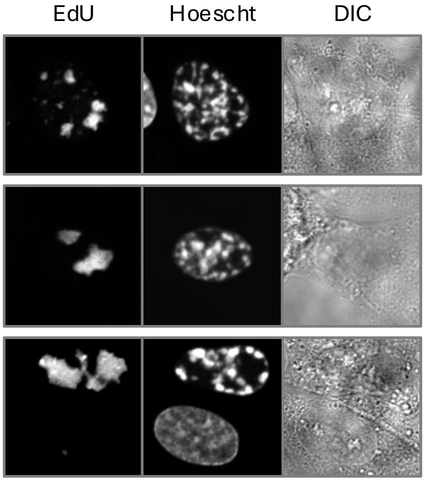
EdU
Hoescht
DIC

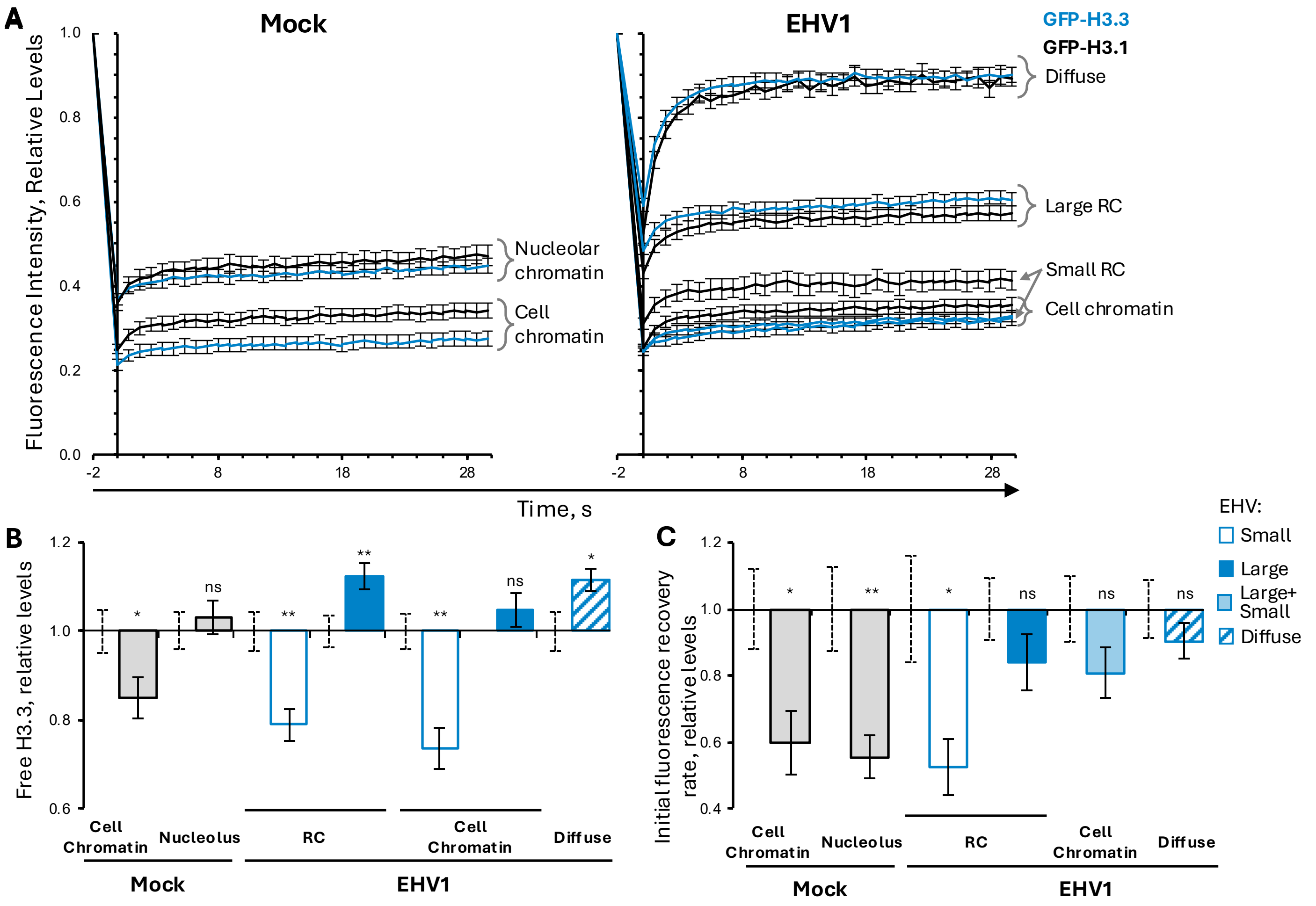
A
Mock
EHV1
GFP-H3.3
GFP-H3.1
Fluorescence Intensity, Relative Levels
1.0
0.8
0.6
0.4
0.2
0.0
-2
8
18
28
Nucleolar chromatin
Cell chromatin
Diffuse
Large RC
Small RC
Cell chromatin
Time, s
B
Free H3.3, relative levels
1.2
1.0
0.8
0.6
ns
*
**
Cell Chromatin
Nucleolus
RC
Diffuse
Mock
EHV1
C
Initial fluorescence recovery rate, relative levels
0.4
EHV:
Small
Large
Large+ Small
Diffuse

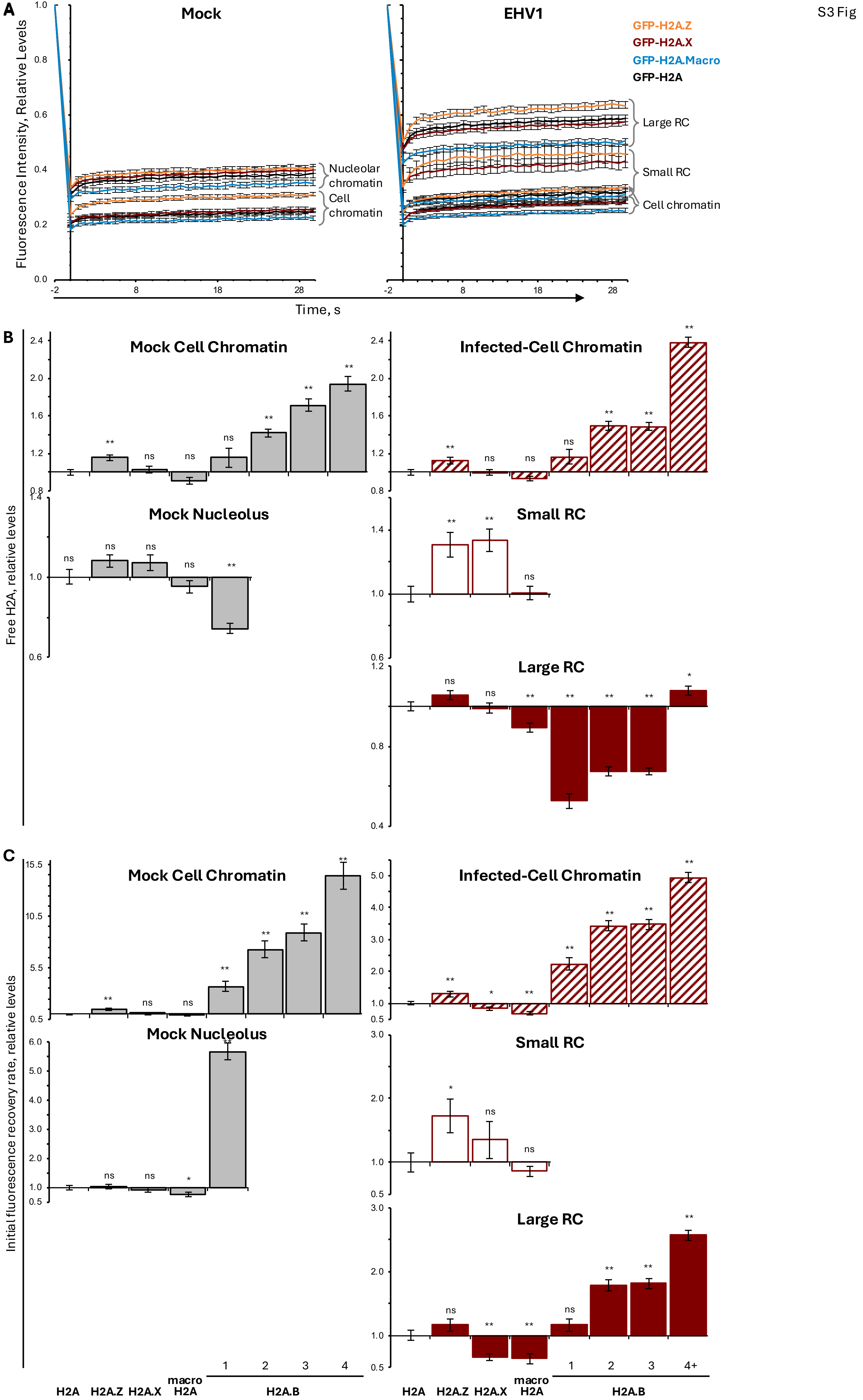
A
Mock
EHV1
GFP-H2A.Z
GFP-H2A.X
GFP-H2A.Macro
GFP-H2A
Fluorescence Intensity, Relative Levels
Nucleolar chromatin
Cell chromatin
Large RC
Small RC
Cell chromatin
Time, s
B
Free H2A, relative levels
Mock Cell Chromatin
Infected-Cell Chromatin
Mock Nucleolus
Small RC
Large RC
C
Initial fluorescence recovery rate, relative levels
Mock Cell Chromatin
Infected-Cell Chromatin
Mock Nucleolus
Small RC
Large RC
H2A
H2A.Z
H2A.X
macro H2A
H2A.B
1
2
3
4
4+

S4 Fig

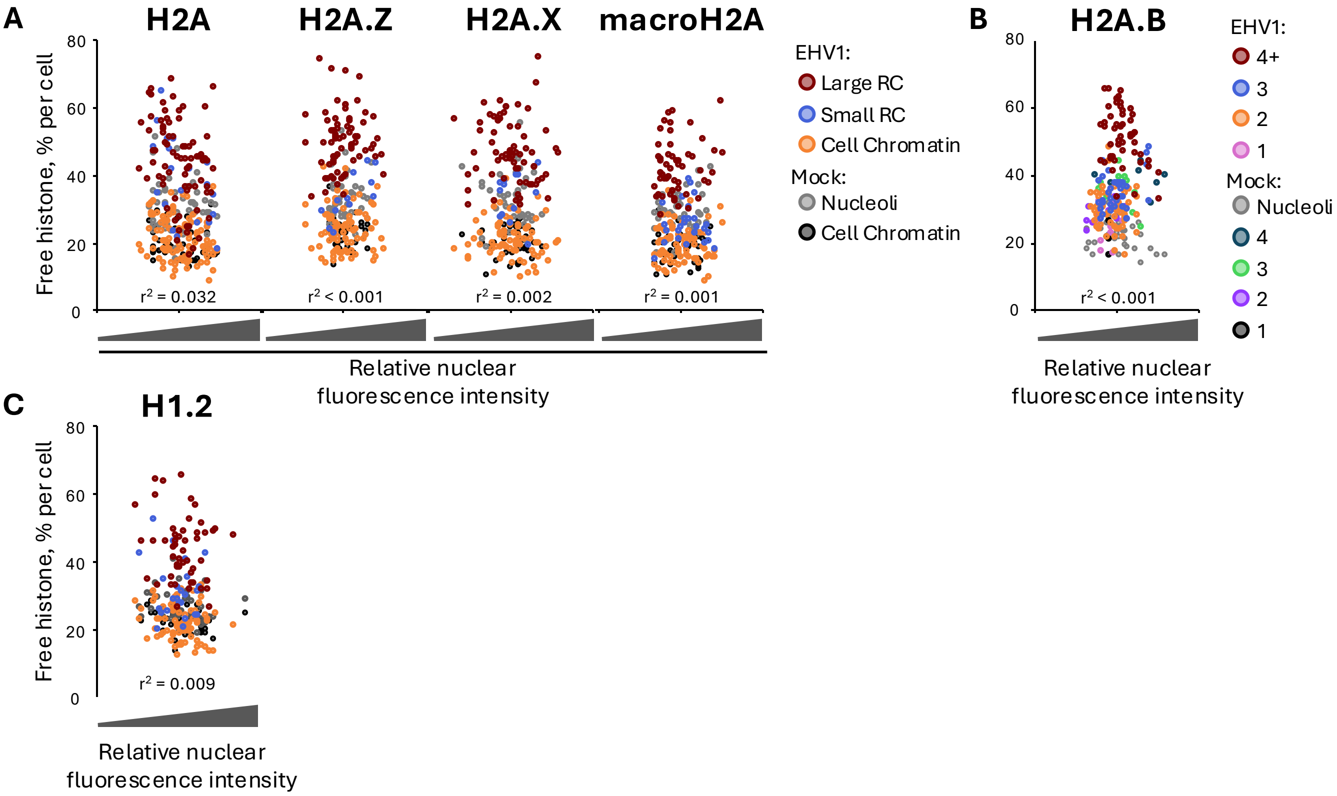
A
H2A
H2A.Z
H2A.X
macroH2A
Free histone, % per cell
80
60
40
20
0
r² = 0.032
r² < 0.001
r² = 0.002
r² = 0.001
Relative nuclear fluorescence intensity
EHV1:
Large RC
Small RC
Cell Chromatin
Mock:
Nucleoli
Cell Chromatin
B
H2A.B
r² < 0.001
Relative nuclear fluorescence intensity
EHV1:
4+
3
2
1
Mock:
Nucleoli
4
3
2
1
C
H1.2
r² = 0.009
Relative nuclear fluorescence intensity

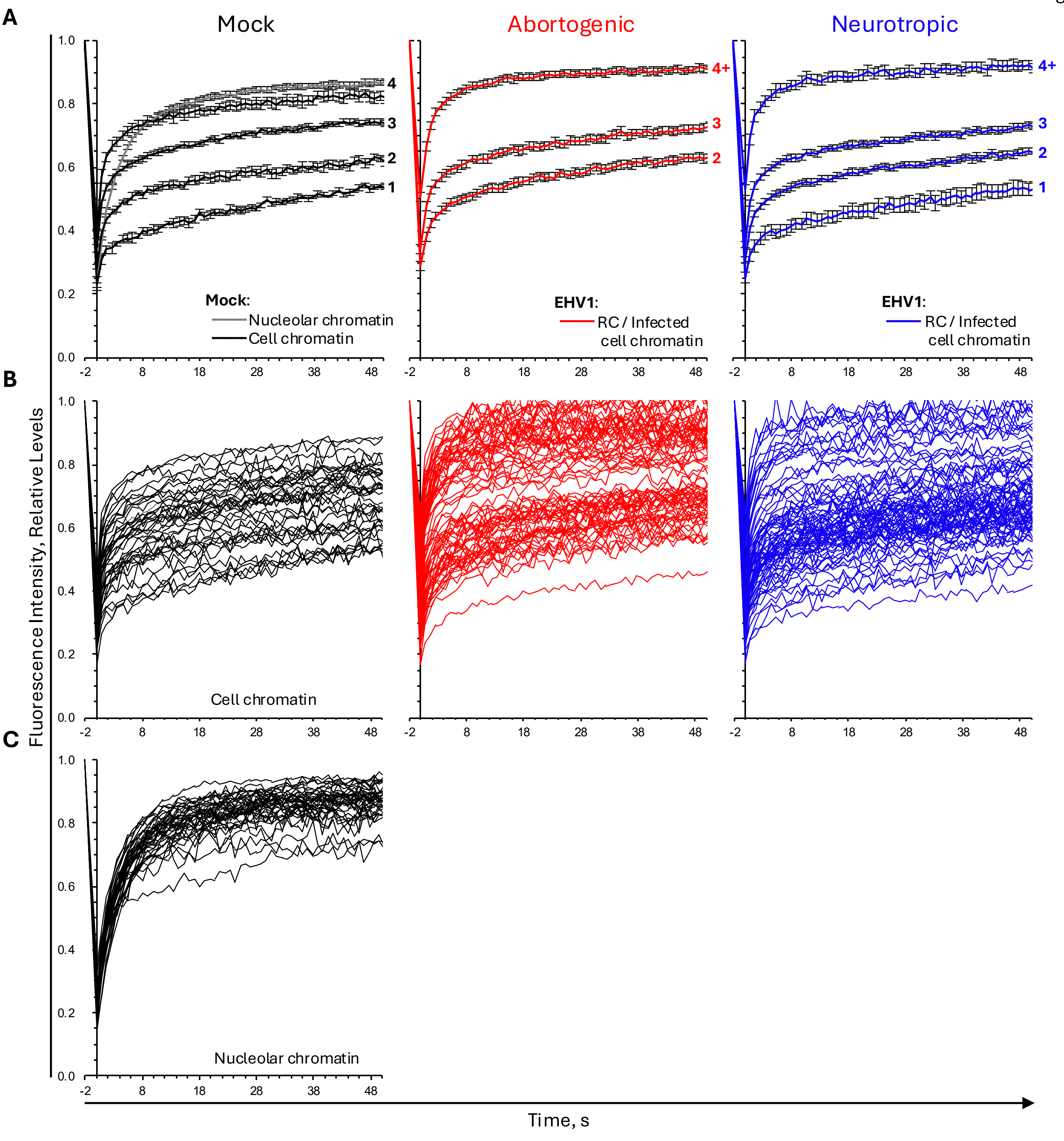
A
Mock
Abortogenic
Neurotropic
Mock:
Nucleolar chromatin
Cell chromatin
EHV1:
RC / Infected
cell chromatin
4
3
2
1
4+
B
Cell chromatin
C
Nucleolar chromatin
Fluorescence Intensity, Relative Levels
Time, s
1.0
0.8
0.6
0.4
0.2
0.0
-2
8
18
28
38
48

## H2A.X

Histone Fold (positions ~27–90); Docking Domain (positions ~81–119)

```
                  10         20         30         40         50         60         70         80         90        100
Consensus  MSGRGKTGGK ARAKAKSRSS RAGLQFPVGR VHRLLRKGHY AERVGAGAPV YLAAVLEYLT AEILELAGNA ARDNKKTRII PRHLQLAIRN DEELNKLLGG
Equine     .......... .......... .......... .......... .......... .......... .......... .......... .......... ..........
Human      .......... .......... .......... .......... .......... .......... .......... .......... .......... ..........

                 110        120        130        140
Consensus  VTIAQGGVLP NIQAVLLPKK TSATVGPKAP XGGKKATQAS QEY
Equine     .......... .......... .......... A......... ...
Human      .......... .......... .......... S......... ...
                                                *  *
```

#### macroH2A

Histone Fold; Docking Domain; Macro Domain

```
                  10         20         30         40         50         60         70         80         90        100
Consensus  MSSRGGKKKS TKTSRSAKAG VIFPVGRMLR YIKKGHPKYR IGVGAPVYMA AVLEYLTAEI LELAGNAARD NKKGRVTPRH ILLAVANDEE LNQLLKGVTI
Equine     .......... .......... .......... .......... .......... .......... .......... .......... .......... ..........
Human      .......... .......... .......... .......... .......... .......... .......... .......... .......... ..........

                 110        120        130        140        150        160        170        180        190        200
Consensus  ASGGVLPNIH PELLAKKRGS KGKLEAIIXP PPAKKAKSPS QKKPVSKKAG GKKGARKSKK QGEVSKAASA DSTTEGTPAD GFTVLSTKSL FLGQKLNLIH
Equine     .......... .......... ........M. .......... .......... .......... .......... .......... .......... ..........
Human      .......... .......... ........T. .......... .......... .......... .......... .......... .......... ..........

                 210        220        230        240        250        260        270        280        290        300
Consensus  SEISNLAGFE VEAIINPTNA DIDLKDDLGN TLEKKGGKEF VEAVLELRKK NGPLEVAGAA XSAGHGLPAK FVIHCNSPVW GXDKCEELLE KTVKNCLALA
Equine     .......... .......... .......... .......... .......... .......... L......... .......... .V........ ..........
Human      .......... .......... .......... .......... .......... .......... V......... .......... .A........ ..........

                 310        320        330        340        350        360        370
Consensus  DDKKLKSIAF PSIGSGRNGF PKQTAAQLIL KAISSYFVST MSSSIKTVYF VLFDSESIGI YVQEMAKLDA N
Equine     .......... .......... .......... .......... .......... .......... .......... .
Human      .......... .......... .......... .......... .......... .......... .......... .
```

## H1.2

Winged Helix Motif

```
                  10         20         30         40         50         60         70         80         90        100
Consensus  MSETAPAAPA AAPPAEKAPV KKKAXKKXXG XXRKASGPPV SELITKAVAA SKERSGVSLA ALKKALAAAG YDVEKNNSRI KLGLKSLVSK GTLVQTKGTG
Equine     .......... .......... ....T..PA. AR........ .......... .......... .......... .......... .......... ..........
Human      .......... .......... ....A..AG. TP........ .......... .......... .......... .......... .......... ..........

                 110        120        130        140        150        160        170        180        190        200
Consensus  ASGSFKLNKK AASGEAKPKX KKXGXXKPKK PXGAAKKPKK AXGXATPKKS AKKTPKKAKK PAAAXVTKKV AKSPKKAKXA KPKKAAKSAX KAVKPKAAKP
Equine     .......... .........A ..T.AA.... .A........ .T.A...... .......... ....A..... ........T. .........P ..........
Human      .......... .........V ..A.GT.... .V........ .A.G...... .......... ....T..... ........V. .........A ..........

                 210
Consensus  KVXKPKKAAP KKK
Equine     ..A....... ...
Human      ..V....... ...
```
